# CYP123, a sterol C4-methyl oxidase, acts with CYP51 to initiate mycobacterial lanosterol catabolism

**DOI:** 10.64898/2026.10.31.760369

**Authors:** Dongxu Yuan, Lauren R. Hatfield, Hannah Ryan, Harry Williams, Mochen Dong, Josie E. Parker, Yi Jin, Rémi Zallot

## Abstract

Mycobacteria exploit host cholesterol, both as a carbon source and as a source of catabolites proposed to modulate host immunity, but their capacity to catabolise other host sterols has not been established. Lanosterol, the first sterol intermediate of host cholesterol biosynthesis, accumulates in the activated macrophages these bacteria infect. Here we define the initiating steps of mycobacterial lanosterol catabolism, combining genomic enzymology, transcriptional profiling, mutagenesis, biochemistry, NMR and crystallography. We identify the cytochrome P450 CYP123 as a sterol C4-methyl oxidase. It is encoded with the lanosterol 14α-demethylase CYP51 in a co-oriented region conserved across most *Mycobacteriaceae*. The region carries no binding site for the cholesterol-responsive regulators KstR or KstR2, and in *M. abscessus* lanosterol induces both genes while cholesterol does not. CYP123 oxidises the C4α-methyl group of C14-demethylated sterols in three steps to the carboxylic acid, yet acts on neither lanosterol nor cholesterol, because Met82 excludes the C14-methylated precursor. An M82A variant binds and oxidises lanosterol. In a single reaction containing both recombinant *M. smegmatis* enzymes, CYP51 converts lanosterol to FF-MAS, which CYP123 then oxidises to the C4-carboxylic acid, so the two act in sequence. *M. smegmatis*, *M. phlei* and *M. abscessus* grow on lanosterol as a sole carbon source. Growth slows in *M. smegmatis cyp123* and *cyp51* mutants. Mycobacterial sterol catabolism therefore extends beyond cholesterol, and blocking the cholesterol route alone may not deprive these organisms of host sterols. Induction where lanosterol confers no growth advantage raises the possibility that the pathway also serves functions beyond carbon acquisition.

**Significance:** Mycobacteria feed on host cholesterol, a dependence long proposed as a drug target but never exploited therapeutically. This work shows that they also catabolise lanosterol, an early cholesterol precursor that accumulates in activated macrophages. Two cytochrome P450s encoded in a conserved, lanosterol-induced locus act in sequence: CYP51 removes the C14-methyl group, and CYP123 then oxidises the C4α-methyl group to a carboxylic acid. This explains why mycobacteria retain CYP51 despite being unable to make sterols, and identifies CYP123 as the first bacterial P450 sterol C4-methyl oxidase and a third enzymatic solution to C4-demethylation. The locus is conserved across more than 90% of *Mycobacteriaceae*, spanning pathogens of humans, livestock and the environment, and contributes to fitness during infection, making both enzymes attractive, azole-binding drug targets. Because cholesterol and lanosterol run through separate routes, depriving mycobacteria of host sterols would require blocking both.

## Introduction

The genus *Mycobacterium* encompasses organisms with major consequences for human and animal health (1). *Mycobacterium tuberculosis*, an obligate human pathogen and the causative agent of tuberculosis, caused an estimated 10.7 million cases and 1.23 million deaths in 2024, and remains the leading cause of death from a single infectious agent worldwide (2). Non-tuberculous mycobacteria (NTM), environmentally ubiquitous organisms commonly found in soil and water, are increasingly recognised as significant opportunistic pathogens (3), and include members of the *M. avium* complex, *M. abscessus*, *M. kansasii*, *M. intracellulare*, *M. gordonae*, and *M. xenopi*, frequently implicated in pulmonary and extrapulmonary infections, with a rising global prevalence (4, 5) estimated at 4% annually (6). The intrinsic antibiotic resistance of mycobacteria, due to low cell-envelope permeability, multidrug efflux systems, and an array of drug-modifying enzymes (7), necessitates prolonged, multi-drug regimens with side effects that often lead to poor patient compliance and the emergence of drug-resistant strains (8, 9). A deeper understanding of mycobacterial metabolism, particularly the exploitation of host-derived resources during infection and persistence (10, 11), could expose vulnerabilities that current regimens do not reach.

Cytochrome P450 enzymes (CYPs) have long been regarded as attractive targets in this context (12), as azole antifungals, which act by inhibiting P450s, inhibit the growth of *M. smegmatis* at submicromolar concentrations and bind tightly to *M. tuberculosis* P450s (13, 14). Mycobacterial genomes encode numerous CYPs (20 in *M. tuberculosis* alone), the vast majority of which have undefined biological functions (15, 16). The best characterised are those of cholesterol catabolism: CYP125 and CYP142 redundantly initiate degradation of the sterol side chain by sequential oxidation of its terminal carbon to the alcohol, aldehyde, and carboxylic acid, priming it for β-oxidation (17, 18), an oxidation that CYP124 also performs *in vitro* without supporting growth (16, 19, 20). That redundancy is not universal: *M. bovis* strains, including var. BCG, encode a defective CYP142, so CYP125 alone supports growth on cholesterol and its deletion abolishes that growth, whereas the equivalent *M. tuberculosis* mutant still grows (18, 19). Side-chain oxidation and ring degradation proceed as independent branches of the pathway (21), so impairment of one does not necessarily block the other. Cholesterol catabolism is a valued drug target: *M. tuberculosis* exploits host cholesterol for persistence in chronically infected lungs (22), *M. abscessus* requires the pathway for macrophage survival (23), and compounds blocking growth on cholesterol have been advanced as antitubercular candidates (24, 25). No therapy exploiting sterol catabolism has yet reached the clinic.

CYP51, the lanosterol 14α-demethylase central to sterol biosynthesis in eukaryotes and the primary target of antifungal azole drugs, is also present in *Mycobacterium* species (26). Although *Mycobacteriaceae* lack the capacity for *de novo* sterol synthesis, their CYP51 homologues retain lanosterol 14α-demethylase activity (26–28). Lanosterol carries two geminal methyl groups at C4, which in eukaryotes are removed one at a time in successive cycles of oxidation, decarboxylation and 3-keto reduction, so that the first cycle yields a 4α-monomethylsterol and only the second gives the 4-desmethyl product (29–32). Why mycobacteria retain CYP51 is therefore unclear, raising the possibility that it acts, with its neighbours in the region, on host-derived lanosterol during infection.

CYP123 and CYP51 are encoded adjacently and in the same orientation, and polymorphisms in *cyp123* have been associated with multidrug resistance (33, 34). The CYP51–CYP123 region is upregulated during macrophage infection of *M. bovis* var. BCG (35). All *M. bovis* var. BCG vaccine strains harbour an ancestral frameshift mutation in *cyp123*, proposed to render the enzyme inactive (36) (Figs. S1 and S2). CYP123 from *M. abscessus* binds azole and triazole antifungal drugs with micromolar affinity and acts on steroid hormones, hydroxylating 11-deoxycorticosterone and progesterone at a position tentatively assigned as C-17 (37). A *cyp123* (MAB_1216c) transposon insertion mutant showed the largest *in vivo* fitness defect of any *M. abscessus* P450 in a pooled transposon sequencing screen of a murine abscess infection model, in both the smooth and rough morphotypes (38), consistent with a role during infection (37). Yet the precise biological function of CYP123 remains unresolved, and its potential role in lanosterol catabolism has not been explored.

Whether lanosterol might be a relevant substrate during infection is not obvious from its abundance alone (39), but TLR4 activation in macrophages causes lanosterol to accumulate through type I interferon-dependent transcriptional repression of the host lanosterol 14α-demethylase CYP51A1 (40). Since both *M. tuberculosis* and *M. abscessus* activate TLR4 in macrophages (41, 42), lanosterol is likely to be enriched in the very macrophages that harbour intracellular mycobacteria. We therefore hypothesised that the CYP51–CYP123 operon constitutes a catabolic module for host-derived lanosterol, extending mycobacterial sterol utilisation beyond cholesterol.

Here we show that *cyp123* lies within a genomic region that is conserved across almost all *Mycobacteriaceae* and that invariably contains *cyp51*. Neither of the cholesterol-responsive regulators KstR and KstR2 has a binding site in the region. In *M. abscessus*, lanosterol induces this operon but cholesterol does not. *M. smegmatis*, *M. phlei* and *M. abscessus* grow on lanosterol as a sole carbon source, and *M. smegmatis cyp123* and *cyp51* mutants show slowed but not abolished growth, whereas *M. bovis* var. BCG, which carries an ancestral *cyp123* frameshift, barely grows on lanosterol. Recombinant *M. smegmatis* CYP123 binds FF-MAS, the direct product of CYP51, and its Δ14-reduced counterpart T-MAS, and catalyses stepwise oxidation of their C4α-methyl group to the alcohol, aldehyde, and carboxylic acid, but does not act on lanosterol or cholesterol. Reconstitution of CYP51 and CYP123 in a single reaction converts lanosterol through to the C4-carboxylic acid, establishing that the two enzymes act in sequence within a catabolic pathway. Crystal structures and molecular docking rationalise the selectivity of CYP123, which has a narrow hydrophobic pocket that excludes C14-methylated sterols. These findings reveal a previously unrecognised sterol catabolic pathway with implications for mycobacterial pathogenesis and drug development.

## Results

### CYP123 is conserved across *Mycobacteriaceae* within an invariant genomic context

To evaluate the distribution of CYP51 and CYP123 across species, a genomic enzymology approach (43) was applied. We generated Sequence Similarity Networks (SSNs) for the cytochrome P450 family (PF00067) in *Mycobacteriaceae* (Taxonomic ID 1762) using the Enzyme Function Initiative web tools. After filtering to exclude fragments and sequences shorter than 300 amino acids, 17,815 UniProt sequences were analysed. CYPs are classified into subfamilies at >55% sequence identity, corresponding to an Alignment Score Threshold (AST) of 131 for this dataset. At this threshold, the SSN resolved discrete clusters for characterised CYPs including CYP51, CYP123, CYP125, and CYP142 (Fig. 1A).

**Figure 1:**
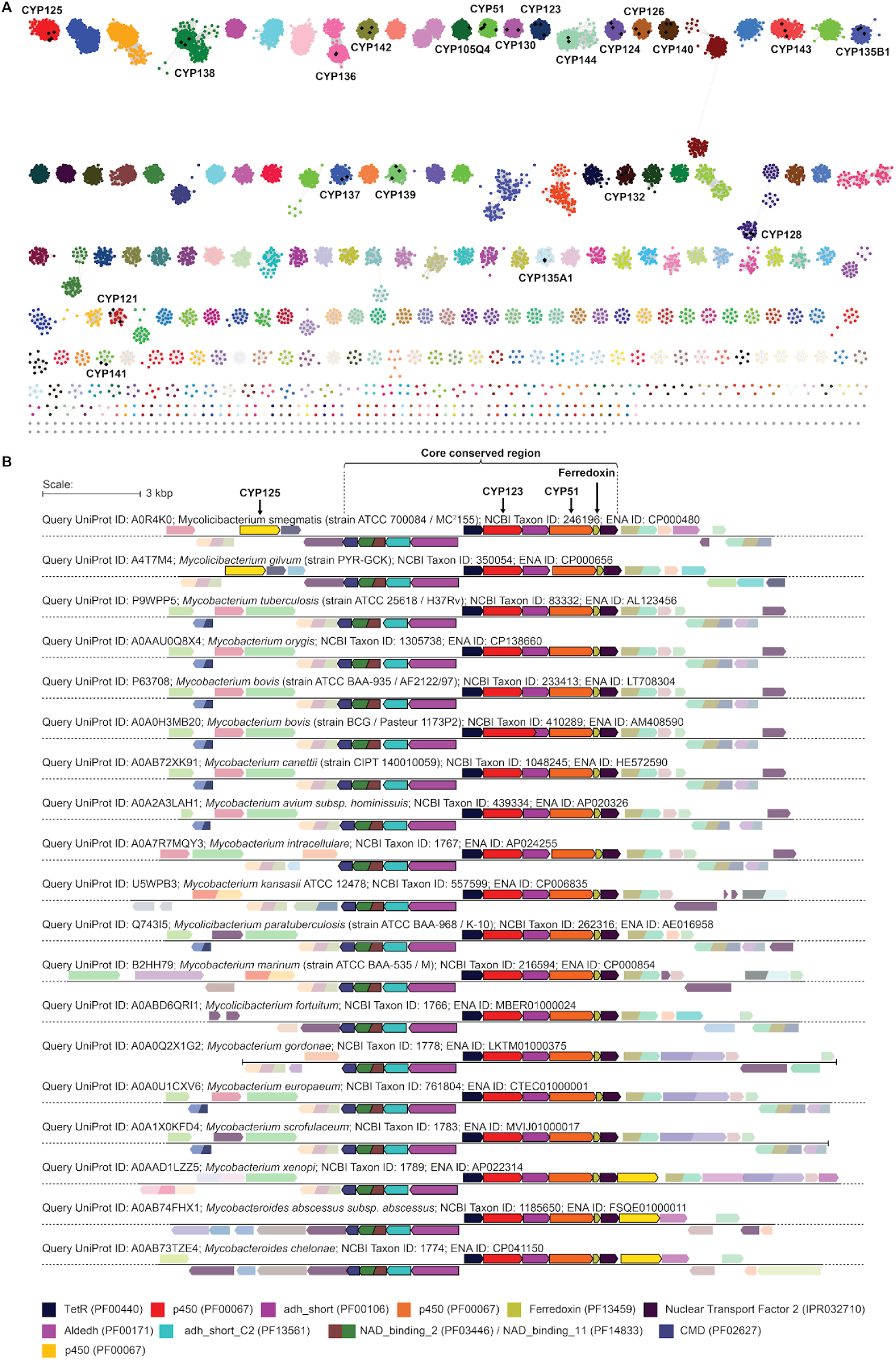
Sequence Similarity Network and Genome Neighbourhood Diagrams for cytochrome P450 (PF00067) sequences from *Mycobacteriaceae*. **(A)** SSN generated at an alignment score threshold of 131 (55% sequence identity) from 17,815 UniProt sequences (minimum length 300 amino acids, fragments excluded). Each cluster is coloured independently; labels indicate clusters containing characterised CYPs. Black diamonds denote SwissProt-annotated reference sequences. **(B)** Genome Neighbourhood Diagrams for 19 representative *Mycobacteriaceae* species, centred on CYP123 (Cluster 15). Genes are shown as arrows indicating transcriptional orientation, coloured by Pfam/InterPro family assignment (key at bottom). The bracket marks the core conserved region spanning from CYP123 to CYP51 and the adjacent ferredoxin; this block of 9 flanking genes, five on one side of CYP123 and four on the other, lying within a window of five genes either side, co-occurs in ≥93% of *Mycobacteriaceae* species (Table S1). CYP125 is located outside the core region but appears in the genomic neighbourhood of CYP123 in several species. Scale bar: 3 kbp.

The 341 *Mycobacteriaceae* species represented encompass the genera *Mycobacterium*, *Mycolicibacterium*, *Mycobacteroides*, and *Mycolicibacter*, spanning all environmental and pathogenic classes (Supplementary Data 1, Zenodo deposit, DOI 10.5281/zenodo.23534052). Sequences from the CYP123 cluster (Cluster 15) are present in 309 species (>90%), with similar prevalence for CYP51 (Cluster 13; 319 species, >93%), CYP125 (Cluster 1; 324 species, >95%), and CYP142 (Cluster 9; 292 species, >85%). The intersection of species containing CYP123, CYP51, and CYP125 encompasses 307 species (>90%). Among named *Mycobacterium* species, only *M. leprae* and *M. lepromatosis* lack all four of these genes.

The CYP123 cluster comprises 383 sequences with a length distribution tightly centred on 404 amino acids. A notable exception is A0A0H3MB20 from *M. bovis* var. BCG (strain Pasteur 1173P2), whose extended C-terminus results from an ancestral frameshift at residue 352 that extends the protein to 541 amino acids, proposed to render the enzyme inactive (36) (Figs. S1 and S2). Genome Neighbourhood Diagrams (GNDs) revealed that CYP123 lies within a genomic region conserved across most *Mycobacteriaceae* (Fig. 1B). In *M. smegmatis*, five genes share the orientation of *cyp123*: a TetR regulator, an uncharacterised short-chain dehydrogenase, CYP51, a ferredoxin, and an uncharacterised Nuclear Transport Factor 2 family protein. Four further genes lie on the opposite strand: a probable NAD-dependent aldehyde dehydrogenase, an oxidoreductase, an oxidoreductase fusion, and a 4-carboxymuconolactone decarboxylase. Within a window of five genes either side of *cyp123*, these nine show a minimum co-occurrence of 93% across *Mycobacteriaceae* (Table S1), and their shared orientation over the *cyp123*–*cyp51* interval is consistent with an operonic arrangement; in *M. tuberculosis*, the corresponding genes are predicted to form a single transcription unit (44). Because the region is near-invariant across the family, what is established in one species is expected to hold for the others that carry it. That expectation underpins the species used here: *M. smegmatis* as a tractable model for biochemical and genetic work, *M. bovis* var. BCG as a natural mutant of the locus, and *M. abscessus* as a clinically significant non-tuberculous pathogen.

CYP51 has been characterised as a lanosterol 14α-demethylase in several mycobacterial species (26), and the adjacent ferredoxin is proposed to support its activity (45); operonic linkage makes that ferredoxin a plausible partner for CYP123 as well. In several genomes, including but not limited to *M. abscessus*, *M. smegmatis*, *M. gilvum*, *M. xenopi*, and *M. chelonae*, a CYP125 paralogue is also found in the wider neighbourhood, whereas CYP142 is not, despite catalysing the same side-chain oxidation (Fig. S3), a selective association that may implicate CYP125 specifically in this pathway.

To ask whether the region is under cholesterol control, we searched it for binding sites of the TetR-type repressors KstR and KstR2, which have been mapped across all intergenic regions of the *M. smegmatis* and *M. tuberculosis* genomes (46, 47). No site of either regulator lies within the *cyp123*–*cyp51* region; in *M. smegmatis*, the nearest KstR site marks the boundary of the cholesterol catabolic gene cluster, some forty genes away. Control is instead likely to be local, through the TetR-family regulator encoded within the locus itself. As *Mycobacteriaceae* lack *de novo* sterol synthesis yet CYP51 acts on lanosterol, this region is proposed to be a dedicated lanosterol catabolic locus, outside the cholesterol-responsive regulon.

To test whether the arrangement is confined to *Mycobacteriaceae*, we built a second SSN encompassing all bacteria. CYP123 and its conserved genomic arrangement extend to phylogenetically related *Actinomycetales*, including *Rhodococcus*, *Nocardioides*, *Gordonia*, *Actinomadura*, *Dietzia*, *Nocardia* and *Prescottella*, and to *Saccharopolyspora* (Figs. S4, S5), suggesting that the pathway described here operates in related actinobacterial pathogens and environmental organisms alike. Notably, a CYP123 orthologue in *Rhodococcus* sp. P14, located within a gene cluster containing a short-chain dehydrogenase matching the conserved arrangement, has been reported to hydroxylate estrone within an estradiol degradation pathway (48). In *Antrihabitans stalactiti* and several *Nocardia* species, genes encoding elements of the Mce4 transport system, including MlaD and MlaE, which mediate cholesterol import in mycobacteria (49), are found adjacent to the lanosterol conserved core region (Fig. S6). In *Nocardia bovistercoris*, a ferredoxin reductase gene lies near the conserved block, though its functional compatibility with CYP123 and CYP51 remains to be determined.

### The CYP51–CYP123 operon is lanosterol-responsive in *M. abscessus*

In *M. abscessus*, *cyp123* (MAB_1216c), *cyp51* (MAB_1214c) and a *cyp125* paralogue (MAB_1211c) lie in one operon (Fig. 1B). The absence of KstR and KstR2 sites predicts that this locus responds to a signal other than cholesterol. We performed qRT-PCR on cultures grown in 7H9 with lanosterol, cholesterol or glycerol; sterols were delivered in a tyloxapol:ethanol (T:E) vehicle, and T:E alone was the reference condition. All p values are Bonferroni-corrected. As predicted, lanosterol induced all three genes, *cyp123* and *cyp51* by ∼6–7 log₂ fold (p = 0.006 and 0.010) and MAB_1211c by ∼4–5 log₂ fold (p = 0.035), while cholesterol induced none (p = 1.00, 1.00 and 0.65, respectively; Fig. 2, Table S2). Glycerol alone also raised *cyp51* expression (p = 0.007), while *cyp123* and MAB_1211c did not respond significantly (p = 0.052 and 1.00), so the operon-wide response was specific to lanosterol.

**Figure 2:**
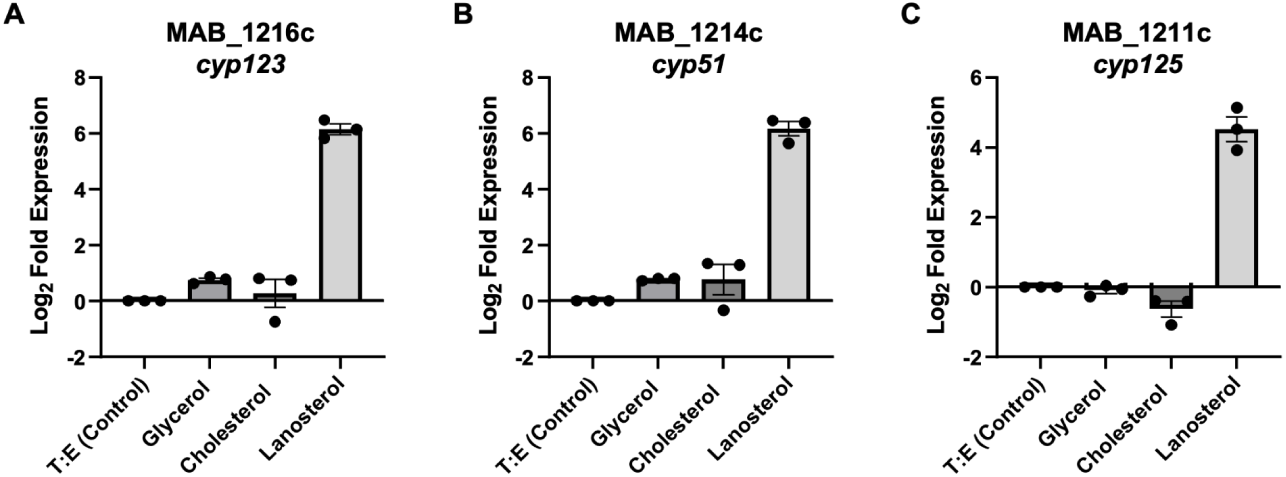
Lanosterol induces the CYP51–CYP123 operon in *M. abscessus*. Relative expression of **(A)** *cyp123* (MAB_1216c), **(B)** *cyp51* (MAB_1214c), and **(C)** *cyp125* (MAB_1211c) measured by qRT-PCR in *M. abscessus* cultured in 7H9 medium supplemented with T:E (control), glycerol (0.2%), cholesterol (0.5 mM), or lanosterol (1 mM). RNA was harvested at mid-log phase. Expression is shown as log₂ fold change relative to the T:E control, calculated using the Pfaffl method and normalised to *recA*. All three operonic genes are strongly induced by lanosterol (p = 0.006, 0.010, and 0.035 for *cyp123*, *cyp51*, and *cyp125* (MAB_1211c), respectively; Welch’s t-test with Bonferroni correction), with no significant cholesterol response. Cholesterol and lanosterol were supplied at different concentrations (0.5 mM and 1 mM, respectively), so the comparison between the two sterols is not concentration-matched. Bars represent mean expression from three biological replicates with individual replicate values overlaid as dots; error bars represent standard error of the mean.

*M. abscessus* encodes four CYP125 paralogues, of which only MAB_1211c lies within this locus. To distinguish co-regulation with the operon from a general sterol response, we tested all four against cholesterol, lanosterol, cholestanol and cholest-5-en-3-one. The operonic paralogue MAB_1211c responded only to lanosterol, matching *cyp123* and *cyp51* (Table S2, Fig. S7); this suggests co-regulation with the operon rather than a role in general sterol catabolism. Neither *cyp123* nor *cyp51* was assayed against cholestanol or cholest-5-en-3-one. Of the three non-operonic paralogues, only MAB_0611 responded to any sterol, being induced by cholesterol (p = 0.023), as expected for a gene lying within the cholesterol utilisation cluster. MAB_0613, two genes away in that cluster, and MAB_0101, elsewhere in the genome, responded to none.

In the same 7H9 medium, we compared growth on each sterol with vehicle alone. Of six sterols tested, cholesterol and cholestanol supported a modest increase in optical density over 72 h, and cholest-4-en-3-one an increase with large variance between replicates, whereas lanosterol, 7-ketocholesterol and cholest-5-en-3-one gave densities comparable to vehicle alone (Fig. S8). These data show that induction of the operon reflects the presence of lanosterol rather than a demand for sterol-derived carbon.

### Lanosterol supports growth of *M. smegmatis*, *M. phlei* and *M. abscessus*

To test lanosterol as a carbon source, we used a minimal medium rather than 7H9, with sterols delivered in tyloxapol. In this medium, *M. smegmatis*, *M. phlei*, *M. abscessus* and *M. bovis* var. BCG grew on glycerol (0.1% v/v) but not on medium alone or on tyloxapol, so neither asparagine nor the vehicle serves as a carbon source (Figs. 3, S9 and S10). In the same medium, cholesterol supported growth of *M. smegmatis* and *M. phlei* in a dose-dependent manner between 0.05 and 1.5 mM (Fig. S10), establishing that a sterol alone can sustain growth under these conditions.

**Figure 3:**
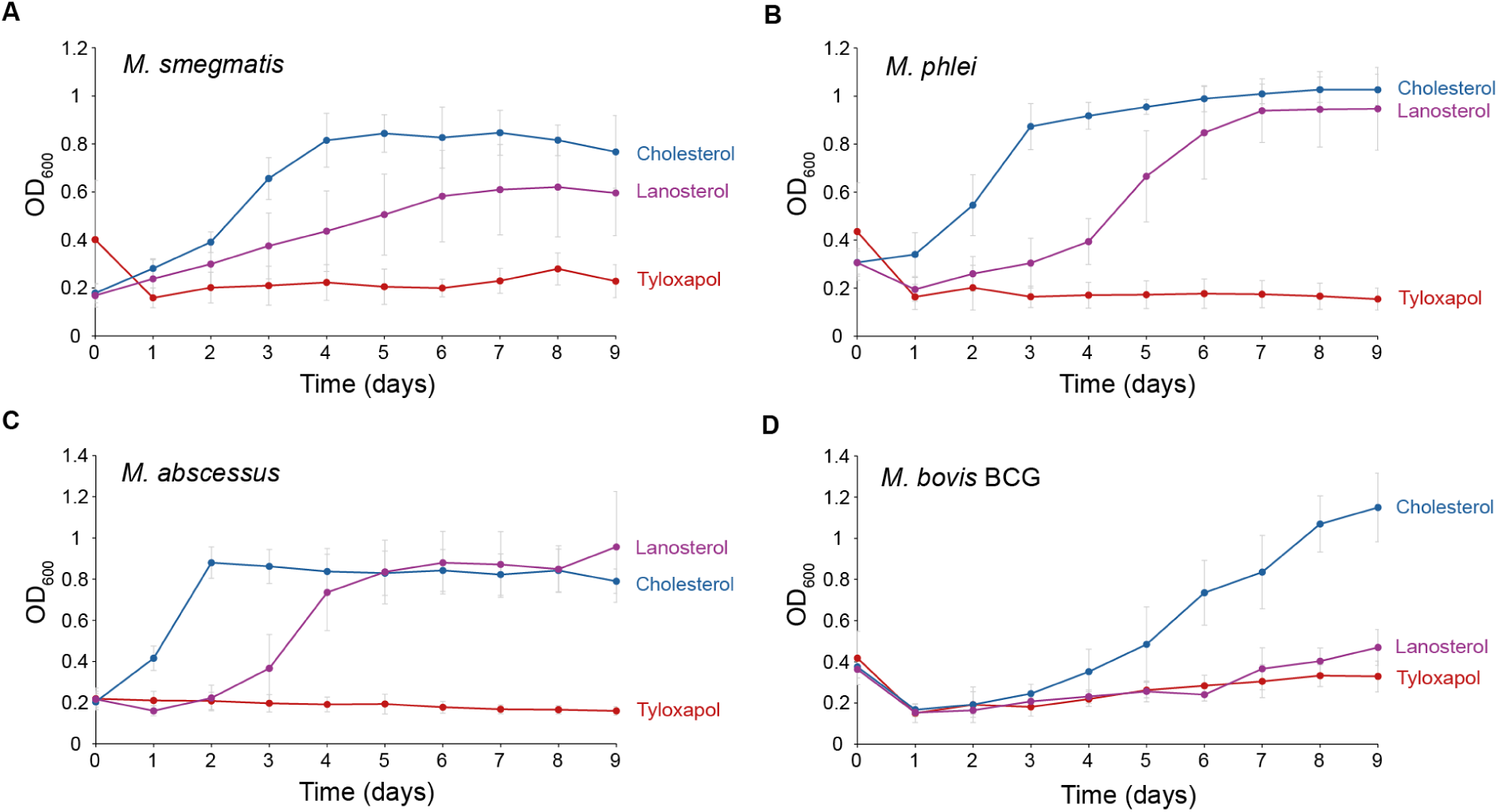
Growth of mycobacterial species on alternative carbon sources. Growth of *M. smegmatis* **(A)**, *M. phlei* **(B)**, *M. abscessus* **(C)** and *M. bovis* var. BCG **(D)** in minimal medium (MM) supplemented with cholesterol (1 mM) or lanosterol (1 mM) as the sole carbon source. Carbon sources were solubilised in tyloxapol before addition. A MM containing tyloxapol (2% v/v) control is included in each panel. OD₆₀₀ was measured every 24 h; data represent the mean ± SD of eight technical replicates. *M. bovis* var. BCG shows a reduced capacity to utilise lanosterol as a carbon source compared to the other species; it harbours a frameshift mutation in *cyp123* predicted to render the encoded enzyme non-functional.

On this basis, we tested lanosterol as the sole carbon source at 1 mM. *M. smegmatis*, *M. phlei* and *M. abscessus* all grew well above the tyloxapol control (Fig. 3A–C). *M. smegmatis* entered log phase later on lanosterol than on cholesterol and reached a lower final OD₆₀₀. *M. phlei* grew strongly on lanosterol, reaching a final OD₆₀₀ comparable to that on cholesterol and higher than on glycerol over the same period (Fig. 3B, S9). *M. abscessus* grew on lanosterol as the sole carbon source, entering log phase later than on cholesterol but reaching a comparable final OD₆₀₀ (Fig. 3C). *M. bovis* var. BCG behaved differently: it grew well on cholesterol (1 mM) and on glycerol (Fig. 3D, S9), but only marginally on lanosterol, barely above the tyloxapol control (Fig. 3D).

### Loss of CYP123 or CYP51 reduces the rate of growth on lanosterol

To test the contribution of CYP123 directly, we disrupted *M. smegmatis cyp123* by CRISPR-Cas-assisted editing (Fig. S11). We evaluated two independently isolated frameshift alleles: *cyp123*Δ17, a 17 bp deletion at gene nucleotides 742–758, and *cyp123Δ136ins228*, a 136 bp deletion at nucleotides 759–894 combined with a 228 bp insertion. Both lesions shift the reading frame and introduce a premature stop codon (Fig. S11), truncating the protein upstream of Cys348, the proximal haem ligand. The encoded enzyme is presumed inactive in both. Whole-genome sequencing confirmed the intended lesion in each strain and identified no plausible off-target site for the guide anywhere in the genome (Zenodo deposit, DOI 10.5281/zenodo.23534052). We complemented each mutant by integrating a single copy of wild-type *cyp123* at the mycobacteriophage L5 *attB* site under a constitutive promoter, verified as single-copy and identical in sequence to wild type.

On lanosterol (0.3 mM) as the sole carbon source, both mutants showed a delayed onset and a shallower subsequent rise in OD₆₀₀, indicating a reduced rate of biomass accumulation, while reaching a comparable final value (Fig. 4A, B). Wild type carrying the parental vector pML1335, which carries *gfipm2+*, was indistinguishable from untransformed wild type. In both mutants the complemented strain grew faster than its parent but more slowly than wild type, while final OD₆₀₀ values remained comparable (Fig. 4A, B). Incomplete rescue is unsurprising: the complementing copy is expressed from a constitutive promoter at a single ectopic site, uncoupled both from the lanosterol-responsive regulation of the native operon and from co-expression of the ferredoxin encoded within it.

**Figure 4:**
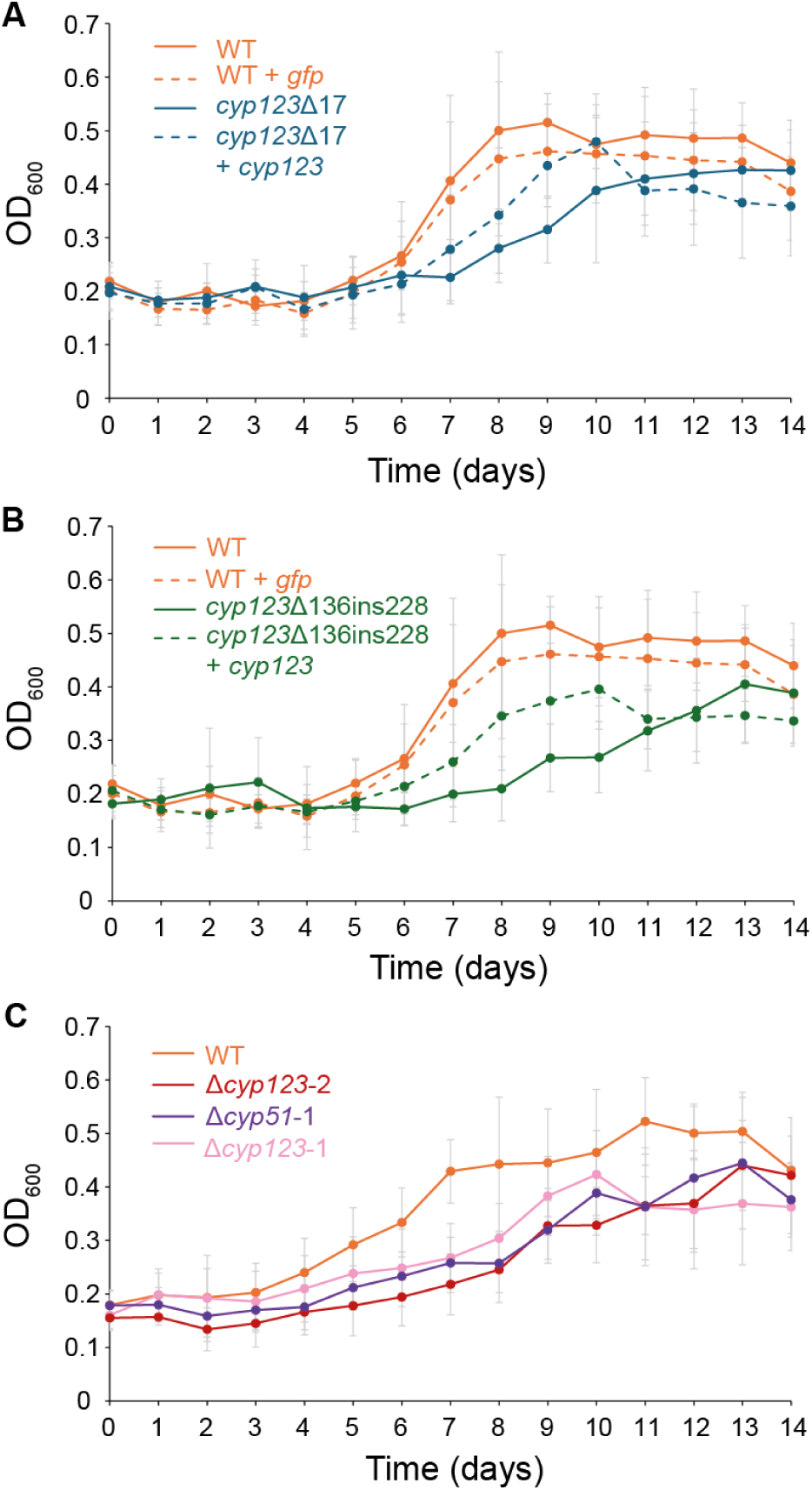
Growth of *M. smegmatis cyp123* and *cyp51* mutants on lanosterol as a sole carbon source. All strains were grown in minimal medium (MM) supplemented with lanosterol (0.3 mM) as the sole carbon source. **(A, B)** CRISPR frameshift mutants and their complemented derivatives: *cyp123Δ17* (solid blue) with *cyp123Δ17* + *cyp123* (dashed blue) in **(A)**, and *cyp123Δ136ins228* (solid green) with *cyp123Δ136ins228* + *cyp123* (dashed green) in **(B)**; the mutant also retains the NHEJ helper plasmid, which its complemented derivative has lost (Methods). Both panels also show wild-type *M. smegmatis* (solid orange) and wild type carrying the parental integrating vector pML1335 (dashed orange, labelled WT + *gfp*), whose *gfpm2+* cassette is replaced by *cyp123* to give the complementing construct. **(C)** ORBIT deletion mutants Δ*cyp123-1* (pink), Δ*cyp123-2* (red) and Δ*cyp51-1* (purple), with wild-type *M. smegmatis* (orange); these mutants could not be complemented (Methods). OD₆₀₀ was measured every 24 h; data represent the mean ± SD of six technical replicates.

To test whether the phenotype depends on the frameshift or on loss of the gene product, we replaced the *cyp123* coding sequence with a marked cassette. Two independent ORBIT replacements of *cyp123* (one retaining only the first 5 bp of the coding sequence, the other 61 bp at its ends) grew more slowly than wild type on lanosterol while reaching a comparable final OD₆₀₀ (Fig. 4C). Deleting *cyp51* by the same strategy reproduced this phenotype, consistent with the two enzymes acting in the same pathway. Because cassette insertion could in principle reduce expression of downstream genes in this closely packed operon (a polar effect), the *cyp51* and *cyp123* deletions cannot be separated on genetic grounds alone. Two observations argue against a purely polar explanation: the CRISPR frameshift alleles, which leave the locus structurally intact, give the same phenotype; and the sequential relationship between the two enzymes is established independently below, by reconstitution of the reaction *in vitro*.

All mutants retained the ability to use lanosterol, and the defect is one of rate rather than of capacity. This is expected if carbon can still be drawn from the side chain: degradation of the sterol side chain by β-oxidation proceeds independently of ring modification and is initiated by CYP125 and CYP142 (17, 18), both intact in *M. smegmatis*. Carbon derived from the lanosterol side chain should remain fully accessible to *cyp123* and *cyp51* mutants, which may account for the delay. Consistent with this, the phenotype was more pronounced at 0.3 mM than at 1 mM lanosterol (Figs. 4 and S12): when substrate is abundant, side-chain carbon alone may suffice, whereas at limiting concentrations the additional carbon released by ring processing may become important for growth. Growth on cholesterol was not impaired in these strains (Fig. S13), confirming that loss of CYP123 or CYP51 impairs lanosterol utilisation specifically rather than sterol catabolism in general.

From these findings, the reason *M. bovis* var. BCG grows so little on lanosterol can be inferred. Lanosterol is likely attacked by two independent branches: ring processing by CYP51 and CYP123, and side-chain degradation by CYP125 and CYP142. BCG carries an ancestral frameshift in *cyp123* predicted to inactivate the enzyme (36), so like the *M. smegmatis* mutants it likely lacks ring processing. Its side-chain route is further compromised, since its *cyp142* is defective (18), leaving CYP125 as its only side-chain oxidase (Fig. 3D, S9). Unlike *M. abscessus* and *M. smegmatis*, BCG has no CYP125 paralogue near *cyp123*; its single *cyp125* lies in the cholesterol utilisation cluster, some 3.1 Mb away. Growth on lanosterol therefore appears to track how many of these branches remain intact: wild-type *M. smegmatis* grows well, the *cyp123* and *cyp51* mutants grow more slowly yet reach the same final density, and BCG, impaired in both branches, grows only marginally. The comparison spans two species, so genotype is not the only variable, but the ordering tracks the branches that remain.

### Recombinant *M. smegmatis* CYP123 selectively binds C4-methylated sterols that lack the C14-methyl group

The genetic and transcriptional evidence places CYP123 within a lanosterol catabolic pathway but does not establish that the enzyme acts on lanosterol-derived sterols directly. CYP123 was therefore produced recombinantly and its ligand preferences measured. Of the *cyp123* genes cloned from *M. smegmatis*, *M. tuberculosis*, *M. marinum*, *Gordonia bronchialis*, *Nocardia afiricana* and *Rhodococcus coprophilus*, only the *M. smegmatis* enzyme could be produced recombinantly in useful yield. The *M. bovis* var. BCG gene could not be expressed in *E. coli* even after the ancestral frameshift was corrected and the gene recoded and codon-optimised (Zenodo deposit, DOI 10.5281/zenodo.23534052).

To characterise the enzyme directly, we purified recombinant C-terminal His-tagged *M. smegmatis* CYP123 and examined it by UV-vis spectroscopy (Fig. S14). Like mycobacterial CYP51 (50), CYP123 is a soluble enzyme, lacking the N-terminal hydrophobic extension that anchors eukaryotic sterol-processing P450s to the endoplasmic reticulum. The enzyme exists predominantly in the low-spin ferric state, with a Soret peak at 418 nm and α-band (568 nm) and β-band (536 nm) absorptions of comparable intensity, consistent with canonical cytochrome P450 spectral properties (51). On reduction and CO binding, the ferrous–CO spectrum showed a split Soret band with peaks at 450 nm and 418 nm (Fig. S14), indicating the characteristic P450–CO complex together with a fraction of inactive P420 (51).

To determine whether CYP123 binds lanosterol-derived compounds, we titrated a panel of steroidal ligands against ferric CYP123 and monitored binding by UV-visible difference spectroscopy (Fig. 5, Table 1). Lanosterol and dihydrolanosterol produced no detectable spectral change at any concentration tested (Fig. 5A, C), and neither did cycloartenol, which also retains the 14α-methyl group (Fig. S15G). Their 14α-demethylated counterparts are bound tightly. FF-MAS (follicular fluid meiosis-activating sterol; 4,4-dimethyl-5α-cholesta-8,14,24-trien-3β-ol), the direct product of CYP51-catalysed 14α-demethylation (26), gave the largest response and the tightest binding in the panel, with an apparent *K*_D_ below 0.05 μM (Fig. 5D, Table 1). T-MAS (testis meiosis-activating sterol; 4,4-dimethyl-5α-cholesta-8,24-dien-3β-ol), formed from FF-MAS by sterol Δ14-reductase in eukaryotic cells (52), gave a type I difference spectrum (53) with a peak at 389 nm and a trough at 421 nm and an apparent *K*_D_ of 0.16 ± 0.02 μM (Fig. 5B). Removal of the C14-methyl group is required for CYP123 binding. We used T-MAS for much of the characterisation that follows, because conversion was more extensive under the conditions tested and the compound was more readily available.

**Figure 5:**
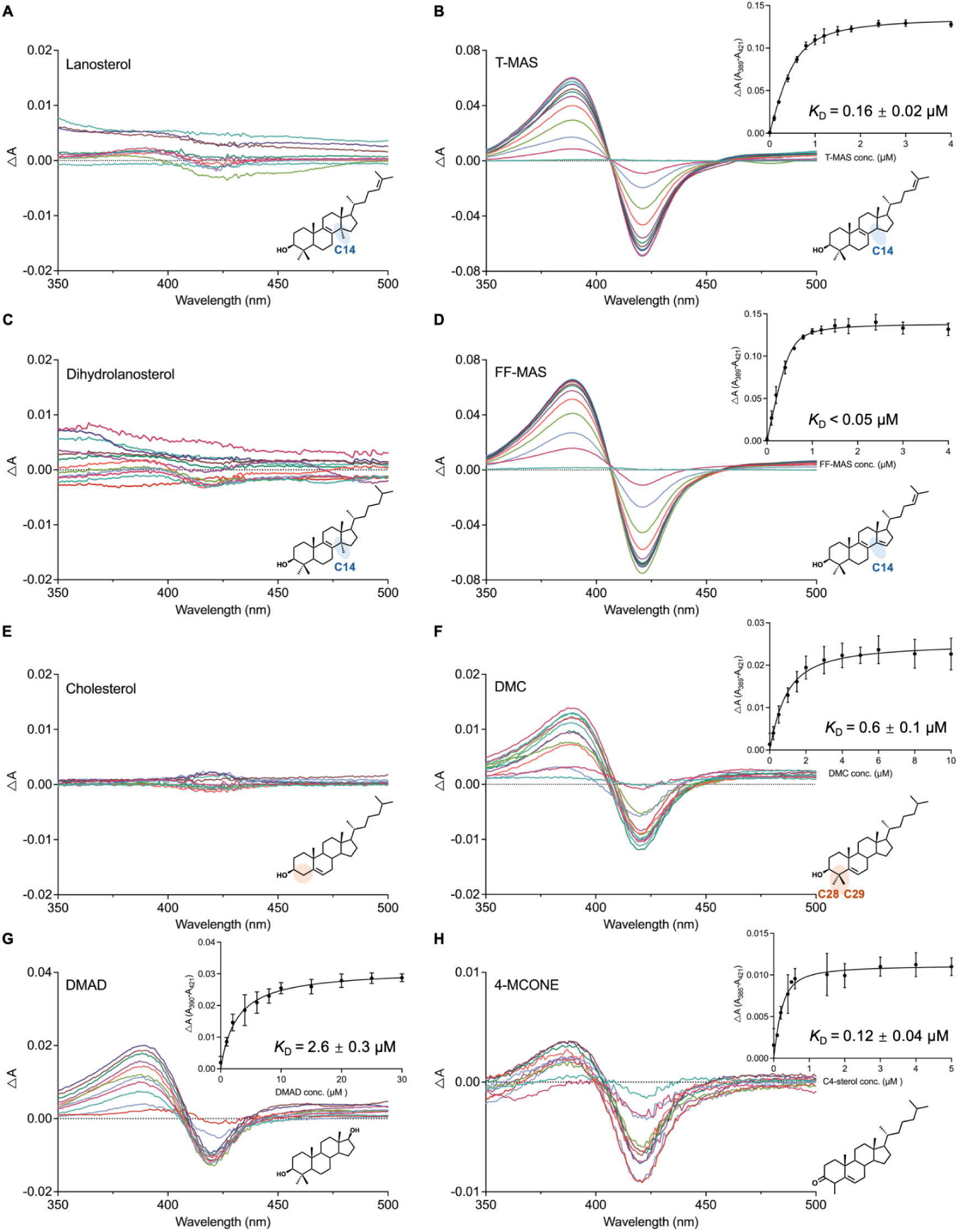
Binding spectra and affinity measurements of sterol ligands with *M. smegmatis* CYP123. Difference spectra recorded on titration of purified recombinant CYP123 (oxidised ferric form, 0.5 μM, 50 mM KPi pH 7.4, 0.1 mM EDTA) with ligands delivered in 40% (w/v) hydroxypropyl-β-cyclodextrin (HPCD), against a reference containing protein and vehicle only. In panels A–F, the left column shows ligands producing no response and the right column their responsive counterparts; panels G and H both show type I responses. **(A)** Lanosterol and **(B)** T-MAS, which differ only in the 14α-methyl group (highlighted blue) and so isolate that feature; FF-MAS, the direct CYP51 product, differs from lanosterol at Δ14 as well and so appears in the less direct comparison below. **(C)** Dihydrolanosterol and **(D)** FF-MAS, which differ not only in the 14α-methyl group but also at Δ14 and Δ24, and so are not related by a single demethylation step. **(E)** Cholesterol and **(F)** DMC, which differ only in the C4 *gem*-dimethyl group (highlighted orange; C-28, C-29). **(G)** DMAD, lacking the C17 side chain. **(H)** 4-MCONE, carrying a single C4-methyl group on a 3-oxo-Δ⁴ ring A. Insets show peak-to-trough absorbance difference (ΔA) against ligand concentration fitted to the Morrison equation for tight-binding ligands (Table 1); no fit is shown for panels A, C and E, where no concentration-dependent change was observed. Note the differing y-axis scales. Data are mean ± SEM of 3 independent titrations.

**Table 1:** Ligand binding parameters for *M. smegmatis* CYP123. Dissociation constants (*K*_D_) were determined by UV-vis difference spectroscopy and fitted to the Morrison equation: ΔA = ΔA_max_ × [([E] + [L] + *K*_D_) − (([E] + [L] + *K*_D_)² − 4[E][L])^0.5^] / 2[E], where [E] is the enzyme concentration, [L] is the ligand concentration, ΔA is the peak-to-trough absorbance difference and ΔA_max_ is the maximum peak-to-trough absorbance difference. Sterols were delivered in 40% (w/v) HPCD; estrone, progesterone and DOC were tested in both HPCD and DMSO, and the vehicle is indicated for each. ND: no measurable binding observed under the stated conditions. Values represent mean ± SEM of 3 independent titrations, except where footnoted. NQ: a response was observed but could not be quantified.

| Ligand | *K*<sub>D</sub> (μM) | Vehicle |
|---|---|---|
| **Type I response** |  |  |
| 4,4-dimethyl-5α-cholesta-8,14,24-trien-3β-ol (FF-MAS) | <0.05<sup>a</sup> | HPCD |
| 4,4-dimethyl-5α-cholesta-8,24-dien-3β-ol (T-MAS) | 0.16 ± 0.02 | HPCD |
| 4,4-dimethylcholest-5-en-3β-ol (DMC) | 0.6 ± 0.1 | HPCD |
| 4,4-dimethyl-5α-androstane-3β,17β-diol (DMAD) | 2.6 ± 0.3 | HPCD |
| 4-methylcholest-4-en-3-one (4-MCONE)<sup>b</sup> | NQ | HPCD |
| **Reverse type I response** |  |  |
| 11-deoxycorticosterone (DOC) | 102 ± 17 | DMSO |
| **No measurable binding** |  |  |
| Lanosterol | ND | HPCD |
| Dihydrolanosterol | ND | HPCD |
| Cholesterol | ND | HPCD |
| Cycloartenol | ND | HPCD |
| Progesterone<sup>c</sup> | NQ | DMSO/HPCD |
| Estrone | ND | DMSO/HPCD |

<sup>a</sup>Lies an order of magnitude below the enzyme concentration used, at the limit resolvable by a tight-binding fit; reported as an upper bound. <sup>b</sup>A type I response was observed but could not be quantified on the same basis as the others: the amplitude was low (ΔAmax ≈ 0.01) and the effective enzyme concentration converged to 0.2 μM during fitting. The fit returned 0.12 ± 0.04 μM, which should not be read as an affinity. <sup>c</sup>No characteristic difference spectrum in HPCD; in DMSO, a type I response that inverted to reverse type I during the titration (Fig. S15).

To isolate the contribution of C4 methylation, we used a commercially available synthetic probe rather than a natural sterol. DMC (4,4-dimethylcholest-5-en-3β-ol) differs from cholesterol only in the C-28 and C-29 methyl groups, so the pair reports on C4 methylation alone. Cholesterol produced no measurable response, whereas DMC gave a type I response with a *K*_D_ of 0.6 ± 0.1 μM (Fig. 5E, F). C4 methylation is required for binding, independently of the other features that distinguish lanosterol-derived sterols from cholesterol.

To probe the C17 side chain and the C4 *gem*-dimethyl group, we tested two further ligands. DMAD (4,4-dimethyl-5α-androstane-3β,17β-diol) retains the 4,4-dimethyl-3β-ol motif but lacks the C17 side chain, and is bound with a *K*_D_ of 2.6 ± 0.3 μM (Fig. 5G), around an order of magnitude weaker than T-MAS, indicating that the side chain contributes to binding without being essential. 4-MCONE (4-methylcholest-4-en-3-one) carries a single C4-methyl group, so it tests whether the gem-dimethyl arrangement is required. It also stands as the closest commercially available proxy for the monomethylated intermediate expected after one round of C4-demethylation. It gave a type I response of much smaller amplitude than those produced by FF-MAS or T-MAS (Fig. 5H). The fit returned 0.12 ± 0.04 μM, reported in the Table 1 footnote rather than as a *K_D_*: the response amplitude was too small to constrain the fit and the effective enzyme concentration converged well below the value used, so it should not be read as an affinity. Its ring A differs from that of the MAS sterols, carrying a 3-oxo-Δ⁴ group in place of the 3β-hydroxyl (54), so the comparison is not exact. A single C4-methyl group is nonetheless sufficient for a response, so the gem-dimethyl arrangement is not required. Because the amplitude of the spectral change reports the fraction of haem converted to the high-spin state, affinities derived from small-amplitude responses carry correspondingly greater uncertainty (Table 1).

To test ligands reported as substrates of CYP123 orthologues, we titrated estrone, described for the *Rhodococcus* sp. P14 orthologue (48), and progesterone and DOC (11-deoxycorticosterone), described for the *M. abscessus* orthologue (37). In HPCD, none gave a difference spectrum (Fig. S15). In DMSO, DOC gave a reverse type I response with a *K*_D_ of 102 ± 17 μM, progesterone a type I response that inverted during the titration, and estrone no response (Table 1). Together, these titrations show that CYP123 binds sterols that are both C4-methylated and C14-demethylated, discriminating against the C14-methylated precursors and against cholesterol alike.

### CYP123 catalyses stepwise oxidation of the C4α-methyl group of sterols

To determine whether CYP123 oxidises the C4-methyl group, we reconstituted the reactions with redox partners. As a Class I cytochrome P450, CYP123 requires a ferredoxin (Fdx) and a ferredoxin-NAD(P)⁺ reductase (FNR) to deliver electrons from NAD(P)H (55). In initial tests with spinach SoFdx–SoFNR and a single addition of NADPH (10 mM), CYP123 oxidised T-MAS and FF-MAS to two products each (Fig. S16). Gas chromatography–mass spectrometry (GC-MS) of *N,O*-bis(trimethylsilyl)trifluoroacetamide (BSTFA)-derivatised extracts assigned them as the corresponding C4-alcohol and C4-aldehyde derivatives. To sustain turnover, we supplemented the reactions with CYP123, SoFdx–SoFNR and NADPH every 30 min. Under these conditions, GC-MS showed progressive conversion of T-MAS and FF-MAS through hydroxylated and aldehyde intermediates to the corresponding carboxylic acids (Fig. 6A–C). Products were assigned from mass shifts relative to the TMS-derivatised substrate: monohydroxylation (+88 Da: +16 Da for the added oxygen and +72 Da for silylation of the new hydroxyl), aldehyde formation (+14 Da, with no additional silylation) and carboxylation (+102 Da: +30 Da for conversion of the methyl to a carboxyl and +72 Da for silylation of that carboxyl, the derivatised acid thus carrying two TMS groups in all); mass spectra of the identified intermediates and products are provided in Figs. S17 and S18. These results establish CYP123 as a sterol C4-methyl oxidase that catalyses the sequential three-step oxidation of a C4-methyl group to the corresponding carboxylic acid.

**Figure 6:**
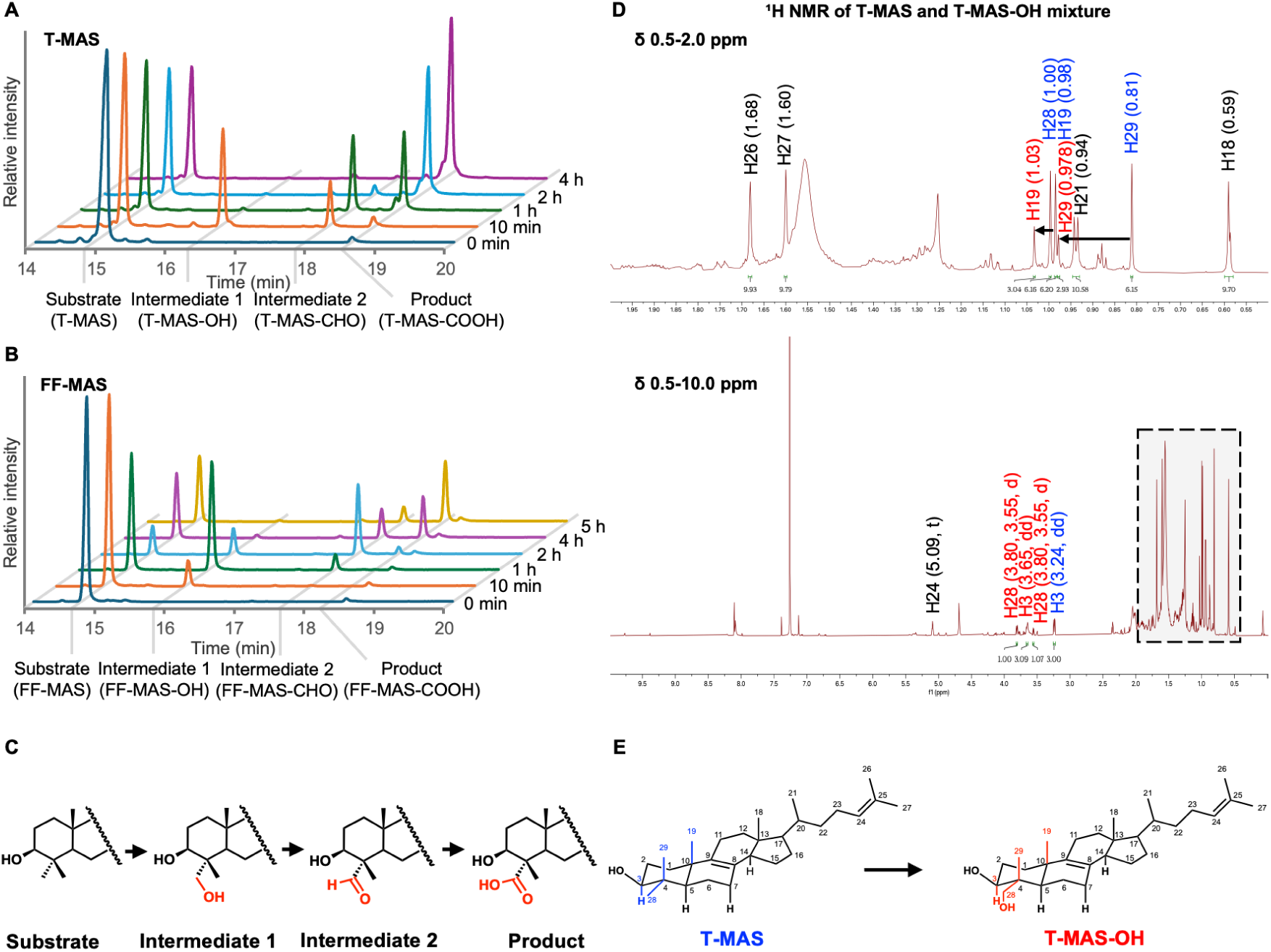
CYP123-catalysed stepwise oxidation of the C4α-methyl group of 14α-demethylated sterols. For the time courses in **(A)** and **(B)**, reactions contained 0.5 mM sterol substrate, 5 μM CYP123, 9 μM SoFdx, 1 μM SoFNR, 0.1 mg mL⁻¹ catalase and 10 mM NADPH, with enzyme and cofactors supplemented every 30 min; products were extracted at the indicated time points and analysed as BSTFA-derivatised extracts on an Agilent 7890B GC coupled to an Agilent 7200 Accurate-Mass Q-TOF fitted with an Agilent VF-5ms column. Mass spectra supporting the peak assignments are given in Figs. S17 and S18. **(A)** GC-MS total ion chromatograms showing the time course of T-MAS oxidation (0 min – 4 h), with sequential formation of T-MAS-OH (C4-alcohol), T-MAS-CHO (C4-aldehyde) and T-MAS-COOH (C4-carboxylic acid). **(B)** Equivalent time course for FF-MAS (0 min – 5 h), showing sequential formation of FF-MAS-OH, FF-MAS-CHO and FF-MAS-COOH. **(C)** Proposed stepwise oxidation pathway: C4-methyl → C4-alcohol → C4-aldehyde → C4-carboxylic acid. **(D)** 1D ¹H NMR spectrum of the preparative-scale reaction mixture of T-MAS and T-MAS-OH (Methods), recorded at 800 MHz in CDCl₃ and shown as two traces: the lower is the full spectrum and the upper an expansion of the methyl region, in which arrows mark the positions whose ¹H resonances shift between the two compounds. **(E)** Structures of T-MAS and T-MAS-OH as determined by NMR, showing oxidation at C-28, the C4α-methyl; positions whose ¹H resonances shift are numbered in blue for T-MAS and red for T-MAS-OH.

Under the conditions used, conversion to the carboxylic acid was more extensive for T-MAS than for FF-MAS, assessed by substrate consumption and acid production, although FF-MAS was the more tightly bound of the two (Table 1). No detectable turnover was observed for lanosterol, cholesterol, cycloartenol, estrone, dihydrolanosterol, progesterone or 11-deoxycorticosterone (Fig. S19A–G). 4-MCONE and DMAD, both of which bind CYP123 (Fig. 5, Table 1), underwent only limited oxidation, generating their corresponding hydroxylated products without progressing further (Fig. S19H, I). For 4-MCONE, the α,β-unsaturated enone formed by its C3-keto group and Δ4,5 double bond may prevent the C4-methyl group from adopting a catalytically competent orientation relative to the haem iron, whereas the absence of the sterol side chain in DMAD may compromise productive positioning within the active site. Binding alone is not sufficient for productive catalysis.

To determine the location of oxidation, intermediates and products from the enzymatic assays with T-MAS and FF-MAS were isolated and analysed by electrospray ionisation tandem mass spectrometry (ESI-MS/MS). Free sterols were detected as dehydrated molecular ions [M−H₂O+H]⁺ owing to water loss during ionisation (56, 57). High-resolution MS/MS spectra of the [M−H₂O+H]⁺ ions revealed fragmentation patterns matching those described by Münger et al. (57) for sterol starting materials and oxidised products (Figs. S20 and S21). Identification of side-chain elimination products in both starting materials and oxidised products argues against oxidation on the sterol tail. Smaller MS fragments narrowed the oxidation site to either the A or B ring, but NMR analysis was required to establish the precise location.

We scaled the reaction up from 0.1 mL to 10 mL containing 1 mg T-MAS (0.24 mM). To simplify the spectra of the mixture, we ran the reaction under deliberately limiting conditions (5 mM NADPH, 5 μM CYP123, 9 μM SoFdx, 1 μM SoFNR and 0.1 mg mL⁻¹ catalase for 10 h) so that conversion terminated predominantly at the alcohol intermediate, as confirmed by GC-MS (Fig. S22). To avoid sample loss, we collected spectra on the crude extract without further purification, affording a mixture of T-MAS and its oxidised alcohol intermediate T-MAS-OH in approximately 2:1 ratio (Figs. 6D and S23). ¹³C NMR (Figs. S24 and S25), HSQC (Figs. S26 and S27), HMBC (Figs. S28 and S29) and ROESY (Figs. S30 and S31) spectra were recorded. Complete ¹H and ¹³C assignments for T-MAS and T-MAS-OH are given in Tables S3 and S4.

Comparison of the ¹H NMR spectrum of the reaction mixture (Fig. 6D) with that of the T-MAS starting material (Fig. S23), revealed new singlet peaks at 0.978 and 1.03 ppm, assigned to the H-29 and H-19 methyl groups respectively. Multiplicity-edited HSQC (Fig. S27B) confirmed these as CH₃ peaks, with carbon environments at C-29 11.28 ppm and C-19 20.37 ppm. The assignment of H-29 was further supported by HMBC correlations (Fig. S29C) to C-4 (38.25 ppm), C-5 (47.01 ppm) and C-3 (79.11 ppm), together with the loss of the correlation to C-28 at 28.11 ppm and a new correlation to the peak at 75.88 ppm, the C-28 CH₂OH carbon. The CH₃ singlet at 1.00 ppm in T-MAS was replaced by two doublets at 3.55 and 3.80 ppm representing a diastereotopic CH₂ group, each proton showing geminal coupling to the other (²*J* = 10.1 Hz). This assignment was supported by HSQC correlations of both protons to the same C-28 carbon signal at 75.88 ppm (Fig. S27) (58). Downfield shifts of H-29 (from 0.81 ppm in T-MAS to 0.978 ppm), H-3 (from 3.24 ppm in T-MAS to 3.65 ppm) and H-19 (from 0.98 ppm in T-MAS to 1.03 ppm) are consistent with their proximity to the electron-withdrawing hydroxyl group at C-28 (59). In T-MAS, the two C4-methyl groups are differentiated as axial (C-29) and equatorial (C-28) within the sterol ring system (Fig. S32) (59, 60). ROESY analysis of T-MAS revealed correlations between the equatorial H-28 (1.00 ppm) and both H-3 (3.24 ppm) and H-5 (1.13 ppm), indicating that they lie on the same face of the ring system (Fig. S30B). In the T-MAS/T-MAS-OH mixture (Fig. S31B), a weak NOE correlation between H-3 (3.65 ppm) and one of the H-28 CH₂ doublets (3.55 ppm) further confirmed oxidation at the equatorial C4α-methyl group (C-28), which lies on the same face as C-3 (Fig. 6E).

### CYP51 and CYP123 act in sequence to initiate lanosterol catabolism

CYP123 does not act on lanosterol (Fig. S19), whereas CYP51 is a lanosterol 14α-demethylase (26, 28) and converts it to FF-MAS. We incubated lanosterol (0.1 mM) with purified *M. smegmatis* CYP51 (Fig. S33), alone or together with CYP123 (Fig. 7). CYP51 alone consumed 73% of the lanosterol and converted it to FF-MAS; adding CYP123 generated FF-MAS-COOH.

**Figure 7:**
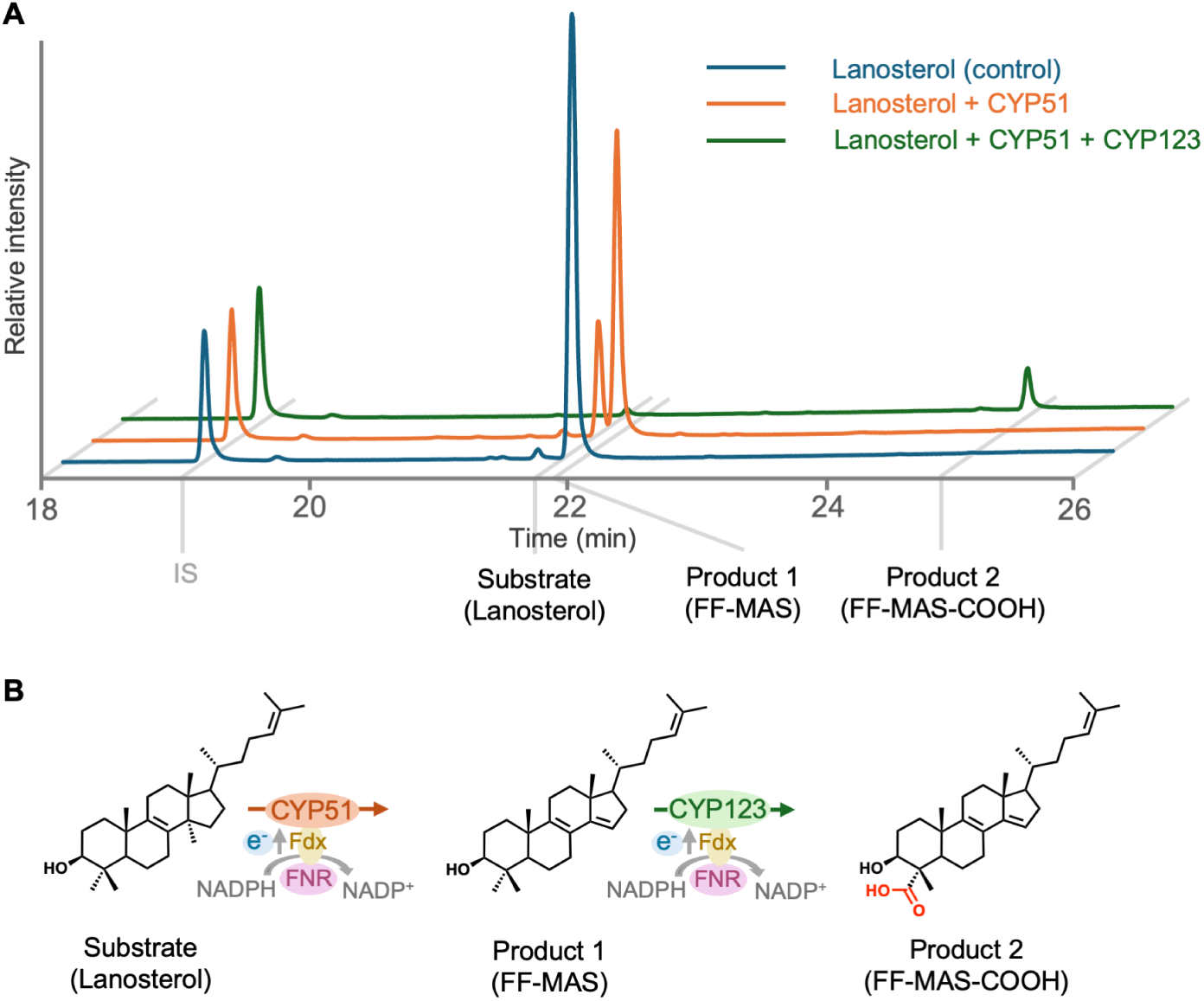
Coupled conversion of lanosterol by CYP51 and CYP123. **(A)** GC-MS total ion chromatograms of BSTFA-derivatised extracts from reactions containing 0.1 mM lanosterol in 50 mM KPi, incubated overnight at 30 °C. Traces, bottom to top: lanosterol with no enzyme; lanosterol with CYP51 (5 μM), SoFdx (9 μM) and SoFNR (1 μM); lanosterol with CYP51 and CYP123 (5 μM each), SoFdx (18 μM) and SoFNR (2 μM). Both enzyme reactions also contained catalase (0.1 mg mL⁻¹), NADPH (1 mM), G6PDH (0.5 U mL⁻¹) and G6P (5 mM). IS, cholesterol, used as an internal standard during lipid extraction from enzyme assay and prior to derivatisation. **(B)** The initiating cascade: CYP51-catalysed 14α-demethylation of lanosterol to FF-MAS, followed by CYP123-catalysed oxidation of the C4α-methyl group to FF-MAS-COOH. Electron transfer from NADPH via ferredoxin reductase (FNR) and ferredoxin (Fdx) supports P450 catalysis.

Because CYP123 cannot act on lanosterol itself, the appearance of a C4-oxidised product in the coupled reaction requires prior 14α-demethylation by CYP51. The two enzymes therefore operate in sequence, carrying lanosterol through to the C4-carboxylic acid. In eukaryotic sterol biosynthesis, FF-MAS is reduced to T-MAS by a dedicated Δ14-reductase before C4-demethylation can proceed. No such step was required here: the coupled reaction contained only CYP51 and CYP123, yet carried lanosterol through to FF-MAS-COOH, and CYP123 oxidises FF-MAS directly (Fig. 5, Table 1). Whether a Δ14-reductase acts at this point in the cell is a separate question. No homologue belonging to this enzyme family, which occurs in eukaryotes and in some bacteria (61), was identified by BLAST searches of *Mycobacteriaceae* proteomes, and the CYP51–CYP123 locus encodes no obvious candidate; whether an unrelated reductase acts elsewhere remains open. Both FF-MAS and T-MAS are intermediates of cholesterol biosynthesis in the host, although the extent to which either is accessible to mycobacteria during infection remains to be evaluated.

### A narrow active site excludes C14-methylated sterols

To define the basis of this selectivity at atomic resolution, we determined the structure of apo-CYP123 at 2.14 Å in space group P4₃2₁2, revealing a canonical cytochrome P450 fold (PDB: 9TJO; Table S5). In the crystal, CYP123 forms a homodimer in which the F–G loop contributes to the dimer interface through salt bridges between Arg40–Glu172′ and Arg171–Asp70′, occluding the active site (Fig. 8A). A hydrogen bond between the His383 and Asp173 side chains stabilises this closed conformation of the F–G loop, a widely recognised dynamic gate for substrate entry (62). The haem is coordinated by the proximal Cys348 thiolate, with a distal water molecule 2.3 Å from the iron, consistent with a resting-state ferric configuration (Fig. 8B). A PEG-bound structure at 2.19 Å (PDB: 9TJP; Table S5, Fig. S34) showed no conformational reorganisation with active-site occupancy by this cryoprotectant-derived ligand.

**Figure 8:**
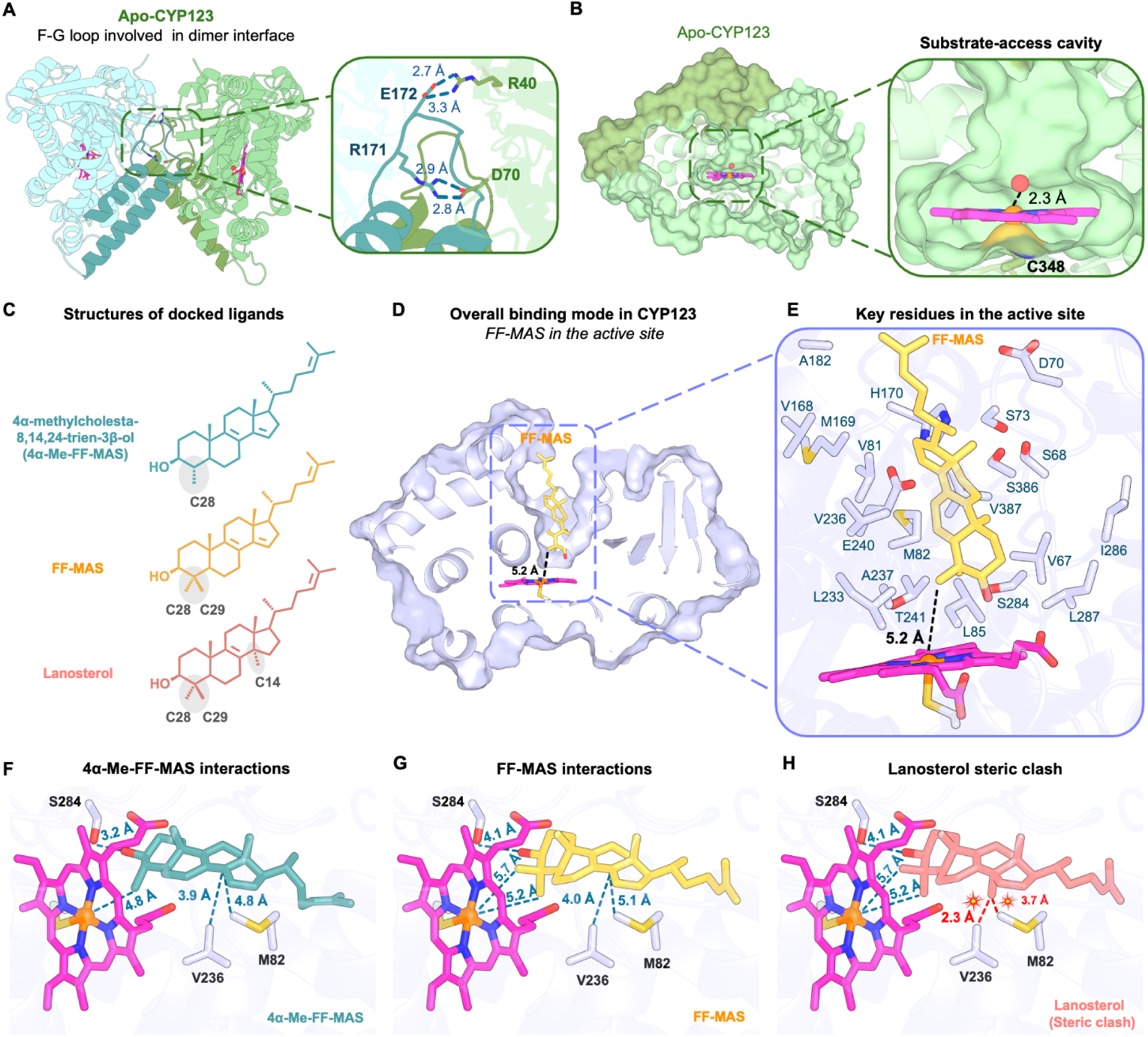
Crystal structure of apo *M. smegmatis* CYP123 and docking analysis of sterol substrates. Throughout, the haem is magenta with its iron as an orange sphere, FF-MAS is yellow, 4α-Me-FF-MAS teal, lanosterol salmon, and active-site residues blue-white; dashed lines indicate distances, reported in Å. **(A)** Crystal structure of apo-CYP123 at 2.14 Å resolution (PDB 9TJO). CYP123 adopts a canonical cytochrome P450 fold and forms a homodimer in which the F–G loop contributes to the dimer interface. Inset: interfacial polar interactions (Arg40–Glu172′ and Arg171–Asp70′) stabilising the closed apo arrangement. **(B)** Surface and active-site views of apo CYP123, with a distal water molecule 2.3 Å above the haem iron. **(C)** Chemical structures of the docked sterols, 4α-Me-FF-MAS (4α-methyl-5α-cholesta-8,14,24-trien-3β-ol) and FF-MAS, together with lanosterol. The C4-methyl groups are labelled C-28 and C-29, and the C14-methyl group of lanosterol is highlighted in grey. **(D)** Predicted binding mode of FF-MAS in the CYP123 active-site cavity, with the protein shown as a surface. **(E)** Enlarged active-site view with FF-MAS positioned above the haem; lining residues are labelled. **(F)** Side view of docked 4α-Me-FF-MAS. The C-28 methyl is oriented towards the catalytic iron (∼4.8 Å) and the C3-hydroxyl forms a hydrogen bond with Ser284 (3.2 Å). **(G)** Side view of docked FF-MAS. The C-28 methyl is oriented towards the catalytic iron (∼5.2 Å) and the C3-hydroxyl is oriented towards Ser284 (4.1 Å). **(H)** Active-site view with lanosterol superposed in the FF-MAS binding pose, showing the close contacts of the C14-methyl group with Met82 and Val236.

To probe substrate recognition, we docked FF-MAS into the apo CYP123 monomer in SwissDock. Poses were filtered on catalytic geometry rather than binding score alone, retaining those placing the C4α-methyl (C-28) towards the haem iron within the narrow, tunnel-like active site (Fig. 8D, G). This filter imposes rather than predicts a productive orientation; the retained poses therefore describe the cavity around a correctly oriented sterol and are not independent evidence that this orientation is adopted.

Twenty-one residues lie within 4.5 Å of the docked sterol, predominantly hydrophobic (Val67, Val81, Met82, Leu85, Val168, Met169, Ala182, Leu233, Val236, Ala237, Ile286, Leu287, Val387) with several polar residues (Ser68, Asp70, Ser73, His170, Glu240, Thr241, Ser284, Ser386) (Fig. 8E). The pocket accommodates the rigid tetracyclic core, with the aliphatic tail extending towards the F–G loop at the entrance; the C3-hydroxyl is oriented towards Ser284 (4.1 Å) and the haem pyrrole (O–H···π, 3.6 Å), and C-28 lies ∼5.2 Å from the haem iron (Fig. 8G). Superposing lanosterol onto the docked FF-MAS places its C14-methyl in close contact with Met82 and Val236, too short for the pocket to accommodate the sterol productively (Fig. 8H), rationalising the preference for C14-demethylated sterols (Table 1).

Because 4α-methyl-5α-cholesta-8,14,24-trien-3β-ol (4α-Me-FF-MAS), the substrate for a second round of C4 oxidation (Fig. 8C), is not commercially available, we modelled it instead; its closest available analogue, 4-MCONE, differs in ring A and is oxidised only as far as the C4-alcohol (Fig. 5, Fig. S19). In this pose, C-28 lies ∼4.8 Å from the haem iron and the C3-hydroxyl hydrogen bonds to Ser284 (3.2 Å) (Fig. 8F), raising the possibility that CYP123 acts in both successive rounds of C4-demethylation downstream of CYP51, although this remains to be established experimentally.

To test the prediction that Met82 and Val236 determine selectivity against C14-methylated sterols (Fig. 8H), we substituted both residues with alanine. The V236A variant showed no detectable type I response to lanosterol (Fig. 9A) and only minimal turnover, which did not proceed beyond the alcohol LAN-OH (Fig. 9C). The M82A variant, by contrast, bound lanosterol with a *K_D_* of 0.27 ± 0.02 μM (Fig. 9D) and converted it far more extensively, through to the aldehyde LAN-CHO (Fig. 9F). The M82A/V236A double variant behaved like M82A alone (*K*_D_ = 0.21 ± 0.06 μM; Fig. 9G, I). LAN-CHO was the furthest product detected within 4 h; whether the third oxidation is prevented or merely slower in this variant was not determined. Docking into each variant accounts for this asymmetry: in V236A, lanosterol remains obstructed by Met82 and the predicted pose is non-productive (Fig. 9B), consistent with the minimal turnover observed, whereas removing the Met82 side chain lets the sterol reposition against the haem in a catalytically compatible orientation in both M82A and M82A/V236A (Fig. 9E, H). All three variants retained the ability to bind T-MAS (Fig. S35), indicating that the substitutions did not compromise the integrity of the active site. These variants identify Met82 as the principal determinant of selectivity for C14-demethylated sterols, with Val236 neither sufficient alone nor additive in combination. That V236A nonetheless formed product without a detectable spectral response indicates that exclusion of lanosterol is leaky rather than absolute.

**Figure 9:**
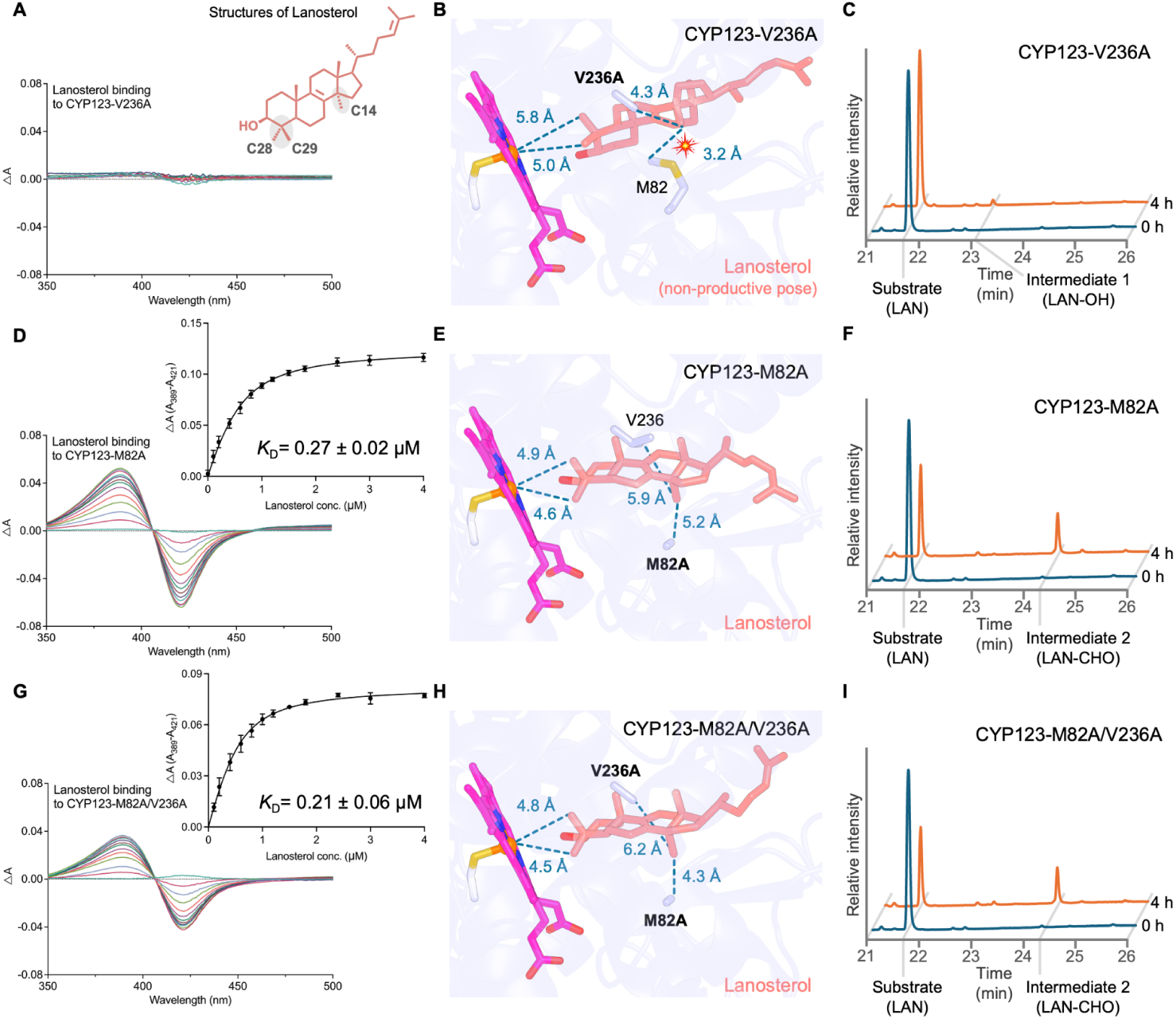
Effects of V236A and M82A substitutions on lanosterol binding and turnover by CYP123. **(A–C)** CYP123-V236A, **(D–F)** CYP123-M82A and **(G–I)** CYP123-M82A/V236A. For each variant: lanosterol binding **(A, D, G)**, the corresponding docking model **(B, E, H)** and GC-MS analysis of lanosterol turnover **(C, F, I)**. No characteristic spectral response was observed for CYP123-V236A; CYP123-M82A exhibited a typical type I spectrum with a *K*_D_ of 0.27 ± 0.02 μM, and the double variant a type I spectral response with a *K*_D_ of 0.21 ± 0.06 μM. Insets in **(D)** and **(G)** show the peak-to-trough absorbance differences plotted against lanosterol concentration, with the corresponding *K*_D_ values indicated. The structure of lanosterol is shown in **(A)**, with the C4-methyl groups (C-28, C-29) and the C14-methyl group highlighted in grey. Docking models show the predicted binding poses of lanosterol relative to the haem and residues Met82 and Val236, or their corresponding alanine substitutions; in CYP123-V236A the sterol adopts a non-productive pose and the red symbol marks its close contact with Met82 **(B)**. Selected distances between lanosterol, the haem and active-site residues are indicated by dashed lines and reported in Å. GC-MS traces are shown at 0 h and 4 h, with peaks labelled as lanosterol (LAN) and its hydroxylated (LAN-OH) and aldehyde (LAN-CHO) products. CYP123-V236A gave only minimal conversion, to LAN-OH **(C)**, whereas CYP123-M82A and CYP123-M82A/V236A converted lanosterol through to LAN-CHO **(F, I)**.

To assess the conservation of functionally important residues, we aligned CYP123 sequences from ten phylogenetically diverse actinobacteria, with *M. smegmatis* CYP123 as reference and representatives selected by CD-HIT from sequences sharing the conserved CYP123 genomic neighbourhood (Fig. S36, Table S6). Per-residue conservation was then mapped onto the apo structure in UCSF ChimeraX (63) using its Coloring by Sequence Conservation function, and displayed as a sequence logo generated with WebLogo (64) (Fig. S37). The haem-binding motif (residues 341–350) is FGSGAHFCLG in *M. smegmatis* and conforms to FGxGxHFCLG across the aligned sequences, matching the canonical P450 signature FGxGxHxCxG, with Cys348 providing the axial thiolate ligand; the conserved residues Arg96 and Arg289 may also coordinate the haem propionates. The I-helix motif (LMVVAGNETTTKLL, residues 233–246) lies deep within the active site adjacent to the haem and is fully conserved; it contains Glu240 and Thr241, the canonical acid–alcohol pair of cytochromes P450, both of which line the substrate pocket in the docked complexes. Conservation is also high across the regions lining the active-site cavity (Val67, Met82–Leu85, Leu233–Ala237, Glu240–Thr241 and Val387), which encompass binding-pocket residues identified by docking. Met82, the principal determinant of selectivity against C14-methylated sterols (Fig. 9), lies within one of these conserved stretches, and Val236 within the fully conserved I-helix motif, so the steric gate that excludes C14-methylated sterols is itself conserved, suggesting that the structural basis for sterol recognition is shared across the CYP123 family.

## Discussion

### Lanosterol catabolism extends mycobacterial sterol metabolism beyond cholesterol

Cholesterol catabolism has been widely recognised as a major determinant of mycobacterial infection, survival, persistence, and adaptation within the host (21, 22, 65). Cholesterol serves not only as a carbon source (66) but also generates catabolites with an emerging role as immunomodulators of the infected host, notably the ring-degradation products androst-4-ene-3,17-dione (AD) and androsta-1,4-diene-3,17-dione (ADD), which suppress LPS-induced inflammatory responses (67). The role of lanosterol, the first sterol intermediate in host cholesterol biosynthesis, has been underappreciated, despite evidence that it accumulates in activated macrophages and modulates innate immune responses (40). Our findings establish that mycobacteria catabolise lanosterol through a conserved CYP51–CYP123 genomic region: following import by a transporter that remains to be identified, CYP51 removes the C14-methyl group and CYP123 then oxidises the C4α-methyl group of the resulting intermediates to the carboxylic acid. Alongside the previously characterised sterol-metabolising P450s CYP51, CYP125 and CYP142, CYP123 broadens the known role of P450 enzymes in mycobacterial sterol metabolism.

The functional annotation of CYP123 illustrates how genomic enzymology can drive the assignment of biological roles to uncharacterised enzymes. Rather than screening arbitrary substrates, a genomic enzymology approach generated testable hypotheses from conserved genomic context: the operonic linkage of *cyp123* with *cyp51*, a known lanosterol 14α-demethylase, predicted that CYP123 would act on lanosterol-derived intermediates rather than on cholesterol derivatives or steroid hormones. This prediction contrasts with the conclusions of enzymatic screening alone, which identified the *Rhodococcus* sp. P14 orthologue as an estrone hydroxylase (48) and *M. abscessus* CYP123 as a steroid hydroxylase acting on progesterone and 11-deoxycorticosterone (37). None of these compounds is bound appreciably by *M. smegmatis* CYP123 (Table 1). Viewed through the lens of genomic enzymology, the reported infection phenotype of a *cyp123* mutant is more parsimoniously explained by a role in lanosterol catabolism (38), providing carbon or signalling metabolites from a host sterol intermediate accessible in the macrophage, than by steroid hormone hydroxylation.

### Resolution of the CYP51 paradox

This substrate logic resolves a long-standing paradox: why mycobacteria retain CYP51 despite their inability to synthesise sterols *de novo*. The initial characterisation of *M. tuberculosis* CYP51 suggested that oxidative portions of a sterol biosynthetic pathway were present in prokaryotes (26), yet no cholesterol could be detected in *M. smegmatis* grown on minimal medium, and the organism appears to lack the oxidosqualene cyclase required to initiate sterol ring formation (27). *M. tuberculosis* CYP51 has consequently been regarded as an “orphan” P450 in a non-sterol-producing organism (68). If CYP51 is considered only as a biosynthetic lanosterol C14-demethylase, its conservation in mycobacteria is difficult to rationalise.

Our results support an alternative interpretation: CYP51 is retained, at least in part, to generate C14-demethylated intermediates that CYP123 can process. The reconstituted reaction makes this explicit. CYP123 does not act on lanosterol, but in the presence of CYP51 the sterol is carried through to the C4-carboxylic acid (Fig. 7A, B), so the two enzymes constitute a sequential module in which CYP51’s product is CYP123’s substrate. Mycobacteria share with eukaryotes a reaction of sterol biosynthesis, but deploy it for the utilisation of exogenous sterol intermediates rather than for endogenous sterol production, which may expand the sterol carbon sources available during pathogenesis.

A key unresolved question is whether lanosterol or CYP51-derived intermediates are accessible to intracellular mycobacteria at physiologically relevant concentrations. *M. tuberculosis* CYP51 has previously been suggested to contribute little to carbon flux during infection because lanosterol is relatively scarce in mammalian cells compared with cholesterol (39). That view is recalibrated by evidence that lanosterol accumulates in TLR4-activated macrophages through type I interferon-dependent repression of host CYP51A1 (40), where it modulates membrane fluidity, reactive oxygen species production and phagocytic activity. Immune activation may therefore create host contexts in which early cholesterol biosynthetic intermediates become more accessible than previously assumed, precisely in the macrophage environments that mycobacteria infect.

### CYP123 represents a third solution to sterol C4-demethylation

In organisms capable of sterol biosynthesis, the two C4 methyl groups derive from cyclisation of 2,3-oxidosqualene. In yeast, their removal requires three enzymes acting in concert: the C4-methyl oxidase ERG25 (29), which oxidises the C4-methyl group to a carboxylic acid; the C3-sterol dehydrogenase/C4-decarboxylase ERG26 (31), which performs oxidative decarboxylation to yield a 3-keto sterol; and the 3-keto sterol reductase ERG27 (32), which regenerates the 3β-hydroxyl and resets the sterol for a second cycle. In *Methylococcus capsulatus*, a distinct bacterial system has been described: SdmA, a Rieske-type oxygenase, and SdmB, an NAD(P)-dependent reductase, convert 4,4-dimethylsterols to 4α-methylsterols using enzymes unrelated to those of the eukaryotic pathway and oxidising the 4β-rather than the 4α-methyl group, although the route still proceeds by decarboxylation (58). CYP123 represents a third enzymatic solution to this step: oxidation of the C4α-methyl group by a cytochrome P450.

The same chemistry has recently been described for CYP5122A1 of *Leishmania donovani*, a bona fide sterol C4-methyl oxidase (59); in *Leishmania major* the orthologue is essential whereas the canonical ERG25 is dispensable, indicating that the P450 carries this step *in vivo* (69). Together these suggest that P450s have been recruited to C4-methyl oxidation at least twice independently, in a trypanosomatid and in a bacterium. The two enzymes are nonetheless distant relatives, sharing around 30% sequence identity across the P450 core, below the 40% threshold that defines a P450 family, and CYP5122A1 carries an N-terminal membrane anchor absent from the soluble CYP123. The shared chemistry likely reflects independent recruitment within the superfamily rather than descent from a common C4-methyl oxidase. The two enzymes differ in ways that reflect their contexts. CYP5122A1 oxidises lanosterol directly, whereas CYP123 cannot act on lanosterol at all and requires prior C14-demethylation by CYP51. CYP123 also carries the oxidation through to the carboxylic acid, whereas the aldehyde is the principal characterised product of CYP5122A1. CYP123 is, to our knowledge, the first P450 C4-methyl oxidase described in a bacterium, and the first operating in a catabolic rather than a biosynthetic pathway.

Under the conditions tested, CYP123 produced a carboxylated sterol but did not complete C4-demethylation. This differs fundamentally from CYP51. There, CYP51 converts the 14α-methyl group successively to the alcohol and the aldehyde, then cleaves the C–C bond and liberates the carbon as formic acid, with concomitant introduction of the Δ14,15 double bond (26, 70); no carboxylate is formed, and demethylation is completed within the P450 reaction itself. CYP123 instead carries the C4-methyl group through to a carboxylic acid that accumulates, so that removal of the carbon would require dedicated downstream enzymes for decarboxylation and 3-keto reduction: an arrangement closer to the ERG25–ERG27 sequence than to CYP51. Additional oxidoreductase genes within the conserved locus are candidates for these steps, and their characterisation is in progress.

The crystal structures provide a structural basis for this substrate logic. CYP123 contains a narrow hydrophobic pocket that accommodates C14-demethylated, C4-methylated sterols (Fig. 8), with specificity appearing to rest primarily on the methylation state and geometry of the sterol core rather than by C17 side-chain conformation. A pose with the C4α-methyl directed towards the haem could be obtained for 4α-Me-FF-MAS as well as for FF-MAS, despite their different side-chain orientations, indicating that the cavity does not exclude the monomethylated sterol and leaving open that a single enzyme serves both rounds, as ERG25 does in yeast. The weaker binding and limited turnover of the tail-less sterol DMAD nonetheless indicate that the side chain is not dispensable and likely provides hydrophobic anchoring in the distal pocket. CYP123 acts preferentially on early lanosterol-derived intermediates that have undergone C14-demethylation but retain a C17 side chain, which would place C4α-methyl oxidation before complete side-chain degradation.

The structures also suggest how CYP123 enforces the order of the two reactions. Met82 excludes lanosterol from the CYP123 active site, and substituting this residue alone permits lanosterol binding and partial oxidation (Fig. 9). Met82 thus acts as a gatekeeper that prevents CYP123 from acting prematurely on C14-methylated sterols, so that productive turnover is largely confined to sterols from which CYP51 has removed the C14-methyl group.

A further distinction from the eukaryotic pathway is physical rather than chemical. In eukaryotes both the sterol 14α-demethylase and the C4-methyl oxidase ERG25 are integral membrane proteins of the endoplasmic reticulum. Mycobacterial CYP51 was the first sterol 14α-demethylase shown to be soluble (26), and it was that solubility which allowed the first CYP51 crystal structure to be determined (71). CYP123 is likewise soluble. Both enzymes of this module act on sterols without the membrane anchoring of their eukaryotic counterparts, a difference that is itself a point of divergence from their human homologues and that may make the pair unusually tractable for structural and inhibitor-binding work. Because sterols partition into membranes, how soluble enzymes meet them in the cell is an open question; in the experiments reported here we presented the sterols in HPCD rather than in a bilayer. Mycobacterial cholesterol catabolism shows that this is no obstacle: a host sterol is imported, and soluble P450s, CYP125 and CYP142, act on it inside the cell (17, 18, 22). The contrast with eukaryotes may therefore be functional rather than incidental: sterol biosynthesis acts on a substrate made in and retained by the membrane, whereas mycobacterial sterol catabolism acts on one that is taken up and consumed.

### The pathway is induced by lanosterol independently of carbon demand

The pathway is lanosterol-responsive and operates independently of the KstR-controlled cholesterol regulon, most likely through the TetR-family repressor encoded within the locus, acting on lanosterol or a lanosterol-derived metabolite. Induction occurs under conditions in which lanosterol confers no growth advantage (Fig. S8), raising the possibility that the pathway serves functions beyond carbon acquisition. One possibility is the generation of lanosterol-derived catabolites with immunomodulatory activity, analogous to the cholesterol-derived AD and ADD described above. This would be consistent with the fitness defect of a *cyp123* transposon insertion mutant in a pooled murine abscess screen (38), where the selective pressure is more plausibly immune modulation than carbon limitation. Brixius et al. reach the same conclusion about immune modulation from a different mechanism, steroid hormone hydroxylation (37).

The growth phenotypes are informative about pathway architecture. In mycobacterial cholesterol catabolism, side-chain degradation by β-oxidation proceeds concurrently with, but independently of, ring modification (21). If the same decoupling applies to lanosterol, loss of CYP51 or CYP123 should slow but not abolish catabolism, which is what the *M. smegmatis* mutants show. *M. bovis* var. BCG is the more severe case. Its ancestral *cyp123* frameshift (36) disables ring modification, while a defective *cyp142* (18) leaves a single, distant *cyp125* for side-chain degradation; it grows only marginally on lanosterol. BCG’s attenuation is attributed in large part to the RD1 deletion (72), but its compound impairment of sterol catabolism may contribute to the metabolic limitations of the vaccine strain. On the basis of the genomic enzymology analysis, we propose that in species carrying more than one *cyp125*, a particular subset of CYP125 homologues may be dedicated to the lanosterol route rather than to cholesterol. Three observations point that way: CYP125, but not CYP142, is repeatedly found beside the locus; in *M. abscessus* only the operonic paralogue responds to lanosterol; and transposon mutants of that paralogue are lost during infection, as are *cyp51* and *cyp123* mutants. Species with a *cyp125* beside the locus would use it for side-chain oxidation of lanosterol; those without would draw on the cholesterol cluster.

The contribution of this pathway during infection has not been tested directly here. All growth dependence experiments used defined carbon sources *in vitro*. The evidence that lanosterol catabolism matters *in vivo* comes from a published transposon sequencing screen of a murine abscess infection, in which *cyp123* and its operonic neighbours were depleted (38). Of 25 annotated P450s, *cyp123* (MAB_1216c) alone was required in both the smooth and rough morphotypes (log₂ fold change −6.8 and −8.0), and *cyp51*, the operon *cyp125* paralogue and the adjacent ferredoxin were also depleted (Dataset S6B of (38)). As these are pooled insertion mutants assayed without complementation, this suggests a requirement for the locus during infection rather than for CYP123 alone. Establishing whether CYP51 or CYP123 is required for survival or persistence within macrophages or in animal infection, and whether the products of the pathway act on the host, is the subject of ongoing work. The enzymology reported here likewise used a heterologous ferredoxin/ferredoxin reductase pair; reconstitution with the operon-encoded ferredoxin remains to be done.

### The locus is conserved across pathogens of humans, livestock and the environment

The reach of this pathway extends beyond human tuberculosis. Non-tuberculous mycobacteria cause pulmonary and extrapulmonary disease that is increasingly reported and difficult to treat (3, 5, 6, 9); in *M. abscessus* the locus is intact, is induced by lanosterol, supports growth on lanosterol as a sole carbon source, and is required during murine infection (38). *M. bovis* causes bovine tuberculosis in cattle, buffalo and wildlife (73, 74), a recognised zoonosis for which a One Health framework is increasingly advocated (75, 76). *M. avium* subsp. *paratuberculosis* causes Johne’s disease in ruminants, and has been proposed as a zoonotic agent although the association remains debated (77). The locus is conserved in over 90% of *Mycobacteriaceae* species examined (Supplementary Data 1) and in related actinobacterial pathogens such as *Rhodococcus* and *Nocardia* (Figs. S4, S5), broadening the potential reach of therapeutic strategies targeting it. The locus is absent from *M. leprae* and *M. lepromatosis*, which lack all four sterol-metabolising P450s examined. Most mycobacteria, however, are environmental organisms of soil and water (3), where the dominant sterols are the 4-desmethyl end products of plant and fungal biosynthesis, principally phytosterols and ergosterol. The routes to these sterols nonetheless pass through 4,4-dimethyl- and 4α-methylsterol intermediates structurally analogous to the substrates of CYP123 (54, 78), and a subset of these accumulates rather than remaining transient: 4α-methyl-Δ7-sterols such as 24-methylene- and 24-ethylidenelophenol reach a few percent of total sterols in plant tissue, and some organisms, including the methanotroph *Methylococcus* capsulatus, accumulate 4-methylsterols as major constituents (54). These sterols lack the C14 methyl group that excludes substrates from the CYP123 active site, and would be accessible to the enzyme directly. Decaying plant and microbial biomass would therefore present 4α-methylsterols as a minor but recurring fraction, raising the possibility that this module evolved to exploit environmental sterols and was later co-opted for host sterol utilisation. The identification of CYP123 as a sterol C4-methyl oxidase may also have industrial value: mycobacteria, particularly *M. neoaurum*, are the workhorses of the phytosterol-to-steroid industry (79), yet no dedicated C4-demethylation enzymes have been identified in these strains and the corresponding steps remain unexploited.

### Sterol catabolic P450s are tractable, azole-binding targets

Cholesterol catabolism has long been proposed as a drug target, yet no therapy exploiting it has reached clinical use. The most advanced clinical candidate, GSK2556286, acts indirectly, raising cAMP through the adenylyl cyclase Rv1625c to downregulate the pathway rather than inhibiting any of its enzymes (24, 25). Direct inhibition of the catabolic enzymes themselves has nonetheless been achieved: fragment-based development against CYP125 and CYP142 has produced dual inhibitors that bind both enzymes with nanomolar affinity and are active against *M. tuberculosis* in whole-cell assays, including intracellular bacteria and a multidrug-resistant strain (80). Sterol catabolic P450s are tractable targets whose inhibition can translate into antimycobacterial activity.

Lanosterol catabolism extends this to a second pathway, with CYP51 and CYP123 providing two further conserved active sites. Both are established azole-binding enzymes: mycobacterial CYP51 has been crystallised with azole inhibitors bound (71), and the *M. abscessus* CYP123 orthologue binds azole and triazole antifungals with micromolar affinity (37). That shared azole affinity offers a starting point for repurposing rather than requiring chemical matter to be developed afresh. With four of the twenty *M. tuberculosis* P450s now implicated in sterol utilisation, this enzyme class may represent a recurring point of vulnerability.

## Conclusion

This study defines the initiating steps of mycobacterial lanosterol catabolism. We show that CYP123 is a sterol C4-methyl oxidase, oxidising the C4α-methyl group of C14-demethylated sterols in three steps through the alcohol and the aldehyde to the carboxylic acid, and that Met82 excludes lanosterol itself, so that CYP51 must act first and the two enzymes together carry the sterol through in a single reaction. The locus that encodes them occurs in more than 90% of *Mycobacteriaceae* species, responds to lanosterol rather than to cholesterol, and lies outside the KstR-controlled regulon that governs cholesterol catabolism. Collectively, these findings place a second host sterol within the mycobacterial catabolic repertoire, and they suggest why bacteria that synthesise no sterols of their own have retained CYP51 throughout the family. They also add a third enzymatic solution to sterol C4-demethylation, one that a cytochrome P450 performs in a catabolic rather than a biosynthetic pathway, alongside the ERG25 route of fungi and the Rieske oxygenase route of methanotrophs. Notably, the sterols this pathway consumes accumulate in the macrophages these organisms inhabit, which may make lanosterol catabolism relevant to infection, although whether it does will require testing *in vivo*. The pathway described here illuminates how far mycobacterial sterol metabolism extends beyond cholesterol.

## Acknowledgements

We would like to thank Dominic Brinkworth, Andrew G.S. Warrilow, Steven L. Kelly (Swansea University), and Callum Breeze (Cardiff University) for preliminary data acquisition. Further, we thank colleagues at the Manchester Institute of Biotechnology (MIB), University of Manchester, in particular, Caroline Gehin for help with sterol sample preparation, Michael Trelore for assistance in running the GC samples, Matt Cliff and Carlo Bawn for assistance with the NMR and data analysis. We thank Maira Hernandez-Guzman and Katherine Hollywood at the University of Manchester Faculty of Science & Engineering Mass Spectrometry and Separations Facility for GC-MS analysis. We thank Colin Levy and Mary Ortmayer (MIB) for their assistance with crystallographic data collection and refinement, and the Diamond Light Source for access to beamline I03 and I04 under proposal number mx31850 and mx38021. We thank the U.S. Department of Energy Systems Biology Knowledgebase (KBase) for access to the KBase Data Lakehouse, on which the genome sequencing analyses were performed. R.Z. is supported by the Academy of Medical Sciences Springboard Award (SBF009/1016) and has previously been supported by the European Union’s Horizon 2020 Research and Innovation Programme under the Marie Skłodowska-Curie Individual Fellowships grant agreement (H2020-MSCA-IF-2018, 839116, deCrYPtion) and by the Manchester Metropolitan University Research Accelerator Grant (2681386). The authors acknowledge the Wellcome Trust and the Royal Society for their support of H. R. and Y. J., with Y. J. being a Wellcome Trust Sir Henry Dale Fellow (218568/Z/19/Z). H. W. is supported by a Cardiff University School of Biosciences Studentship.

## Author contributions

All authors contributed to the manuscript and approved the final version. **CRediT author statement: Dongxu Yuan**: Data curation, Formal analysis, Investigation, Validation, Visualization, Writing – review & editing. **Lauren R. Hatfield**: Data curation, Formal analysis, Investigation, Validation, Visualization, Writing – review & editing. **Hannah Ryan**: Investigation, Formal analysis, Writing – review & editing. **Harry Williams**: Investigation, Formal analysis. **Mochen Dong**: Investigation, Formal analysis. **Josie E. Parker**: Investigation, Formal analysis, Supervision, Writing – review & editing. **Yi Jin**: Conceptualization, Data curation, Formal analysis, Funding acquisition, Investigation, Visualization, Supervision, Writing – original draft, Writing – review & editing. **Rémi Zallot**: Conceptualization, Data curation, Formal analysis, Funding acquisition, Investigation, Project administration, Supervision, Validation, Visualization, Writing – original draft, Writing – review & editing.

## Materials and Methods

### Bioinformatics

The Enzyme Function Initiative (EFI) web tools (https://efi.igb.illinois.edu/) were used to generate and analyse Sequence Similarity Networks (SSNs) (43, 81). All analyses were performed in July 2025 using the EFI-implemented versions of UniProt (2025–03) and InterPro (82), with ENA data downloaded in June 2025. The Taxonomy tool was used to define an initial dataset of UniRef90 sequences from the PF00067 family restricted to bacteria. From this, full UniProt SSNs were generated for PF00067 members of the family *Mycobacteriaceae* (taxonomic ID 1762); selected analyses were performed on the complete bacterial PF00067 UniRef90 set. To reduce noise, fragments and sequences shorter than 300 amino acids were excluded. Alignment Score Thresholds (ASTs) were defined at the intersection of the median sequence identity and the corresponding alignment score, such that edges were drawn between sequences sharing at least 40% identity for grouping into a family and more than 55% identity for grouping into a subfamily. For the *Mycobacteriaceae* SSN these corresponded to ASTs of 79 and 131, respectively; for the bacterial UniRef90 dataset they were 73 and 126. SSNs were analysed using Color SSN and Cluster Analysis to retrieve UniProt identifier lists, sequence-length distributions and colour-mapping files for clusters of interest. A species was scored as encoding a cluster member when at least one of its representative genomes contained a sequence within that cluster. Reported cluster sequence counts refer to the full UniProt set; CD-HIT was used only to select representative sequences for the alignment figures. Of the 17,815 sequences analysed, 17,620 fall into 268 clusters and 195 are singletons. SSNs were further analysed with the Genome Neighbourhood Tool using default parameters and, separately, more stringent settings (window size 5, minimum co-occurrence 50%) to determine gene-neighbour co-occurrence ratios and to generate Genome Neighbourhood Diagrams. Networks were visualised and annotated in Cytoscape v3.10.3 (83) with the BridgeDb plug-in (84). Sequence alignments were performed with Clustal Omega v1.2.4 and visualised using the EMBL-EBI suite of tools (85). For the alignment of bacterial CYP123 sequences, Genome Neighbourhood Tool output was obtained for the members of Cluster 24 of the bacterial PF00067 SSN (Fig. S4) with genome neighbourhood data in ENA (899 sequences), from which the 472 whose neighbourhoods carry the conserved gene set of Table S6 were retained and reduced with CD-HIT. This selection was made before the 300 amino acid length minimum was applied, so one of the 472 sequences is shorter than 300 amino acids; it is not among the representatives selected below. Representative sequences for alignment figures were selected with CD-HIT v4.8.1 (86), clustering at ≥70% identity over ≥80% of sequence length (cd-hit -c 0.7 -n 5 -aS 0.8). CD-HIT was run in Google Colab using a notebook archived at https://doi.org/10.5281/zenodo.23503573. Protein structures were visualised and analysed in PyMOL (Schrödinger, LLC) and UCSF ChimeraX (63).

### Strains and Culture Conditions

*Mycobacterium smegmatis mc²155* (ATCC 700084) was the primary mycobacterial strain used in this study. Under the revised taxonomy of the family this species is *Mycolicibacterium smegmatis*, and *M. abscessus* is *Mycobacteroides abscessus*; the genus name *Mycobacterium* is used for both throughout, following common usage. *Mycobacterium abscessus* (ATCC 19977) was used for transcription studies, and the growth of *Mycobacterium phlei* (DSM 43239) and *Mycobacterium bovis* var. BCG (NCTC 14044) was monitored on a range of carbon sources. Genomic DNA was extracted from *Gordonia bronchialis* (NCTC 10667), *Rhodococcus coprophilus* (NCTC 10994), *Mycobacterium marinum* (NCTC 2275) and *Nocardia afiricana* (NCTC 13184); genomic DNA from *M. tuberculosis* H37Rv was also used as a PCR template. *Escherichia coli* XL1-Blue MRF′ (Stratagene) was used for routine cloning, *E. coli* DH5α for Gibson assembly, and *E. coli* Rosetta2 (DE3) for recombinant protein expression. All strains used in this study are listed in Table S7.

Mycobacterial strains were maintained in Middlebrook 7H9 medium supplemented with 0.5% (v/v) tyloxapol and 0.2% (v/v) glycerol. *M. smegmatis, M. phlei* and *M. bovis* var. BCG cultures were additionally supplemented with 10% (v/v) ADC (bovine albumin, dextrose and catalase); *M. abscessus* was grown without ADC.

### Cloning

Genomic DNA was extracted following established methods (87). All primers and oligonucleotides are listed in Table S8. Cloning of the *cyp123* orthologues that could not be produced in useful yield, including the *M. bovis* var. BCG gene and its recoded derivatives, is described in the Zenodo deposit (DOI 10.5281/zenodo.23534052).

#### M. smegmatis cyp123

The *cyp123* gene was amplified from *M. smegmatis* mc²155 genomic DNA using primers listed in Table S8 and cloned into pET28a for recombinant expression, giving a C-terminally hexahistidine-tagged product. Insert and vector were digested with NcoI and XhoI according to the manufacturer’s protocol and purified using the PCR Purification Kit. Ligation was performed with T4 DNA Ligase at an insert-to-vector ratio of 3:1 and incubated at 16 °C overnight. The ligation mixture was transformed into *E. coli* XL1-Blue MRF′ by heat shock at 42 °C for 45 s; transformants were recovered in SOC medium at 37 °C for 1 h with shaking at 220 rpm and plated on LB agar containing 50 μg mL⁻¹ kanamycin. Colony PCR with Dream Taq Green DNA Polymerase and T7 forward and reverse primers were used to identify clones carrying an insert of the expected size. Plasmid DNA was extracted using the MiniPrep Kit and the insert verified by Sanger sequencing with the same primers.

#### M. smegmatis cyp51

The *cyp51* gene was amplified from *M. smegmatis* mc²155 genomic DNA using primers carrying 20 bp overlaps complementary to linearised pET28a. Gibson assembly was performed with NEBuilder HiFi DNA Assembly Master Mix at an approximate insert-to-vector molar ratio of 3:1, incubated at 50 °C for 1 h and transformed into chemically competent *E. coli* DH5α by heat shock at 42 °C for 45 s. Transformants were recovered and plated as above, screened by colony PCR and verified by Sanger sequencing. The confirmed plasmid, encoding a C-terminally hexahistidine-tagged protein, was transformed into *E. coli* BL21(DE3) for expression.

#### CYP123 active-site variants

Site-directed mutagenesis was performed by whole-plasmid PCR with Q5 High-Fidelity DNA Polymerase, using pET28-CYP123 as the template. The M82A and V236A single variants were generated independently with the corresponding mutagenic primer pairs (pET28-MsCYP123-M82A-F/R and pET28-MsCYP123-V236A-F/R, respectively; Table S8). The amplified products were treated with *Dpn*I digestion, purified using the QIAquick PCR Purification Kit and transformed into *E. coli* DH5α competent cells. Individual colonies were picked, and the introduced mutations were verified by Sanger sequencing. The sequence-confirmed plasmids were designated pET28-CYP123-M82A and pET28-CYP123-V236A. To construct the M82A/V236A double variant, a second round of whole-plasmid PCR mutagenesis was carried out using a confirmed single-mutant plasmid as the template under the same conditions. The resulting construct was verified by Sanger sequencing and designated pET28-CYP123-M82A/V236A. All primers used for mutagenesis are listed in Table S8.

### Construction and Complementation of *M. smegmatis* Mutants

#### CRISPR-assisted editing

*cyp123* (MSMEG_5861) was edited using the CRISPR-assisted genome editing method of Yan et al. (88). Protospacer adjacent motif (PAM) sequences within *cyp123* were identified with CRISPOR (89), and sgRNAs were designed as the 20 bases immediately upstream of the PAM 5′ end, preceded by a BbsI site. The sgRNA used targeted nucleotide 762 of *cyp123* (Table S8), and both alleles reported here derive from it. sgRNAs were shuttled in pYC2085 (Addgene #158721), a gift from Yi-Cheng Sun (88). Constructs were electroporated into competent *M. smegmatis* carrying either pNHEJ-recX-sacB (Addgene #158718) or pNHEJ-recAmu-sacB (Addgene #158717), also gifts from Yi-Cheng Sun (88). Cells were recovered in 7H9 supplemented with ADC and plated on 7H10 supplemented with 10% (v/v) OADC (oleic acid, bovine albumin, dextrose and catalase), kanamycin (20 μg mL⁻¹) and hygromycin B (50 μg mL⁻¹). Helper plasmids were removed by growth on 7H10 containing 10% (v/v) OADC and 2% (w/v) sucrose. Whole-genome sequencing showed that *cyp123*Δ136ins228 retains the helper plasmid, which has sustained a 1,311 bp deletion truncating sacB; the reconstructed plasmid is deposited as GenBank QB217605 and so is not curable by sucrose counter-selection; its complemented derivative has lost it.

Two *cyp123* alleles were obtained and validated by amplicon sequencing (Fig. S11), with whole-genome sequencing used to confirm the absence of off-target mutations: a 17 bp deletion at nucleotides 742–758 causing a frameshift (*cyp123*Δ17) and a 136 bp deletion spanning nucleotides 759–894 with a 228 bp insertion at nucleotide 758, also causing a frameshift (*cyp123*Δ136ins228). Nucleotide positions are numbered from the *cyp123* start codon. The two frameshift alleles *cyp123*Δ17 and *cyp123*Δ136ins228 were used for phenotypic characterisation.

#### ORBIT recombineering

The ORBIT strategy of Murphy et al. (90) was used to delete *cyp123* (MSMEG_5861) and *cyp51* (MSMEG_5863) in *M. smegmatis* using the recommended plasmids pKM461 and pKM464, replacing the target gene with a payload plasmid. The published method was followed with two alterations: recovery after electroporation was extended beyond the suggested overnight incubation, and plating was on 7H10 agar supplemented with OADC and hygromycin B (25 μg mL⁻¹). The payload plasmid integrates in full, so the resulting cassette carries two resistance genes: *aph(7″)-Ia*, conferring the hygromycin resistance used for selection in *M. smegmatis*, and *cat* from the pKM464 backbone, which is selectable only in *E. coli*. The integrated cassette is deposited as ORBIT-cat-hyg (GenBank QB217607). The *cyp51* cassette carries no synthetic ribosome-binding site, unlike the two *cyp123* cassettes. Two *cyp123* and one *cyp51* mutants were obtained and validated by amplicon sequencing (Fig. S11). The three ORBIT mutants are designated Δ*cyp123*::*hyg*-1, Δ*cyp123*::*hyg*-2 and Δ*cyp51*::*hyg*-1, abbreviated hereafter to Δ*cyp123*-1, Δ*cyp123*-2 and Δ*cyp51*-1. In Δ*cyp123*-1 the payload plasmid replaced *cyp123* from nucleotide 31, and in Δ*cyp123*-2 from nucleotide 6; in Δ*cyp51*-1 it replaced *cyp51* from nucleotide 31. Whole-genome sequencing confirmed the absence of off-target mutations in Δ*cyp123*-1, Δ*cyp123*-2 and Δ*cyp51*-1.

#### Complementation

Mutants were complemented using the integrating plasmid pML1335 (91), together with pBS-Int (92), as previously described (93). The *gfipm2+* cassette of pML1335 was replaced with *cyp123*, giving pCYP123-comp (GenBank QB217606), allowing integration at the L5 *attB* site under the control of the constitutive Psmyc promoter. The ORBIT mutants could not be complemented by this route: the hygromycin resistance marker of pML1335 is nucleotide-identical to that of the ORBIT cassette over 1,236 bp and so affords no selection in a cassette background. Complementation controls therefore derive from the CRISPR alleles. Mutant and complemented strains were verified by PCR and by Oxford Nanopore whole-genome sequencing.

#### Genomic DNA extraction

Genomic DNA for amplicon and whole-genome sequencing was extracted from 10 mL log phase cultures of each strain using Zymo Research Quick-DNA Fungal/Bacterial Microprep Kit, and quantified and assessed for integrity by NanoDrop and Qubit.

#### Amplicon sequencing

Genomic DNA from five strains (wild type, *cyp123*Δ17, *cyp123*Δ136ins228, Δ*cyp123*-1 and Δ*cyp123*-2) was used to amplify *cyp123* and from two strains (wild type and Δ*cyp51*-1) to amplify *cyp51*. *M. smegmatis cyp123* and *cyp51* were amplified using Phusion High-Fidelity DNA polymerase and ORBIT detection primers (Table S8) to amplify outside the gene. Amplicons were cleaned using AMPure XP beads before quantification and assessment of quality by agarose gel, NanoDrop and Qubit. Amplicons were sequenced using the SQK-RBK114-24 rapid barcoding kit run using a FLO-MIN114 (R10.4.1) flow cell on an Oxford Nanopore MinION Mk1D. Demultiplexed raw FASTQ reads underwent quality assessment, filtering, and trimming via fastcat v0.18.6. Reads were aligned to the *Mycobacterium smegmatis* mc²155 *cyp123* (MSMEG_5861) or *cyp51* (MSMEG_5863) reference sequence using minimap2 v2.28.

#### Whole-genome sequencing

Genomic DNA from eight strains (wild type, *cyp123*Δ17, *cyp123*Δ136ins228, Δ*cyp123*-1, Δ*cyp123*-2, Δ*cyp51*-1, and the *cyp123*Δ17 and *cyp123*Δ136ins228 complemented strains) was sequenced on a single multiplexed Oxford Nanopore MinION Mk1D run using a FLO-MIN114 (R10.4.1) flow cell and SQK-RBK114-24 rapid barcoding. Reads were basecalled on-instrument with dna_r10.4.1_e8.2_400bps_hac@v5.2.0, filtered with Filtlong v0.3.1, assembled *de novo* with Flye v2.9.6 (--nano-hq) and polished with Medaka v2.2.2 using the model matched to the basecaller, giving 23–78× depth per strain. Assemblies were annotated with Bakta v1.12.0 and assessed with QUAST v5.3.0. Target loci were extracted from each polished assembly and aligned against the corresponding wild-type sequence to determine the structure of each lesion, and read pileups spanning each lesion were classified to confirm clonality. Off-target variants were called with Clair3 v2.0.2 using the matched ONT model and intersected with whole-genome assembly alignments (MUMmer v4.0.1); coverage was computed with mosdepth v0.3.14 and alignments handled with minimap2 v2.31 and SAMtools v1.24. Variants were called against the laboratory wild-type assembly rather than the public mc²155 reference, which differs from it by 139 SNPs and 357 indels that would otherwise be reported as editing off-targets. Candidate off-target sites were additionally screened by searching both strands of the wild-type assembly for near-matches to each protospacer. A single clonal SNV shared by *cyp123*Δ17 and its complemented derivative survived filtering; it is absent from every other strain including the wild type and so marks that lineage rather than an editing event. Analyses were performed on the KBase Data Lakehouse, a computational resource of the U.S. Department of Energy Systems Biology Knowledgebase (KBase) (94, 95).

### Protein Expression and Purification

Recombinant *M. smegmatis* CYP123 and CYP51 were expressed in *E. coli* Rosetta2 (DE3) and BL21(DE3) carrying the corresponding pET28a-derived plasmids. Overnight pre-cultures were grown at 37 °C in LB medium containing 50 μg mL⁻¹ kanamycin and with 34 μg mL⁻¹ chloramphenicol included for Rosetta2 (DE3) cultures. Pre-cultures (0.1% v/v) were used to inoculate autoinduction medium (96), which was incubated at 37 °C with shaking at 220 rpm for 4 h and then at 25 °C overnight. Cultures expressing CYP123 and CYP51 were supplemented with 2 mM δ-aminolevulinic acid to support haem biosynthesis.

Cells were harvested by centrifugation at 6,900 × g for 30 min at 4 °C and stored at −80 °C. Pellets were resuspended in lysis buffer (50 mM potassium phosphate, KPi, pH 7.4, 500 mM sodium acetate, 1 mM tris(2-carboxyethyl)phosphine, 10% (v/v) glycerol, 1 mM phenylmethylsulfonyl fluoride) and lysed in a high-pressure cell disruptor (Constant Systems Ltd, UK) at 20 kpsi. Insoluble material was removed by centrifugation at 25,400 × g for 40 min at 4 °C.

Proteins were purified by Ni-NTA affinity chromatography followed by size-exclusion chromatography (SEC). CYP123 and CYP51 were purified on a 5 mL HisTrap column (Cytiva) using an ÄKTA Start system. Columns were equilibrated in buffer A (100 mM KPi, pH 7.4, 10% (v/v) glycerol), washed with buffer B (100 mM KPi, pH 7.4, 500 mM NaCl, 20% (v/v) glycerol), and eluted with a linear gradient of 10–80% buffer B containing 1% (w/v) histidine (26). Ni-eluted fractions were purified by SEC using a HiLoad 16/600 Superdex 75 pg column (Cytiva) equilibrated with buffer C (100 mM KPi, pH 7.4, 500 mM NaCl), isocratic elution was performed at a flow rate of 0.6 mL/min. Protein fractions were collected based on absorbance at 420 nm (A₄₂₀), analysed by SDS-PAGE, pooled on the basis of purity, concentrated, and buffer-exchanged into storage buffer (50 mM KPi, pH 7.4, 10% (v/v) glycerol) using Vivaspin 20 centrifugal concentrators (Sartorius). Purified proteins were stored at −80 °C. All CYP123 variants were expressed and purified using the same procedures as those used for wild-type CYP123.

### CO-Difference Spectroscopy

Cytochrome P450 concentration was determined by the carbon monoxide difference spectrum assay (97). Purified CYP123 and CYP51 were diluted into 1 mL of assay buffer (50 mM KPi, pH 7.4, 0.1 mM EDTA) in a quartz cuvette and UV-visible spectra were recorded between 250 and 600 nm on a BioSpectrometer kinetic spectrophotometer (Eppendorf). The protein was reduced with sodium dithionite, the cuvette sealed, and approximately 1 cm³ of CO gas introduced by syringe. Spectra were recorded at 1 min intervals until formation of the ferrous–CO complex was complete. Difference spectra were generated in GraphPad Prism by subtracting the oxidised spectrum from the ferrous–CO spectrum, and the concentration of active P450 was calculated using ε₄₅₀₋₄₉₀ = 91 mM⁻¹ cm⁻¹ (98).

### Ligand-Binding Spectral Titrations

Titrations were performed on a Cary 1 Bio UV-visible spectrophotometer. CYP123 was diluted in 50 mM KPi (pH 7.4) containing 0.1 mM to 0.5 μM EDTA. Sterols were prepared as 1 mM stocks in 40% (w/v) hydroxypropyl-β-cyclodextrin (HPCD), a vehicle used previously to solubilise sterols for work with mycobacterial cytochrome P450s (18).

Lanosterol (catalogue no. T4970; purity 99.67%) was purchased from TargetMol. Cycloartenol (catalogue no. 08172; purity ≥90% by GC), cholesterol (catalogue no. C8667; purity ≥99%), estrone (catalogue no. E9750; purity ≥99% by HPLC), progesterone (catalogue no. P0130; purity ≥99% by HPLC), and 11-deoxycorticosterone (catalogue no. D6875; purity ≥97% by HPLC) were purchased from Sigma-Aldrich. Dihydrolanosterol (catalogue no. 700067P; purity >99% by TLC), T-MAS (catalogue no. 700073P; purity >99% by TLC), and FF-MAS (catalogue no. 700077P; purity >99% by TLC) were obtained from Avanti Research. 4-Methylcholest-4-en-3-one (4-MCONE; catalogue no. R209694), 4,4-dimethyl-5α-androstane-3β,17β-diol (DMAD; catalogue no. R190861), and 4,4-dimethylcholest-5-en-3β-ol (DMC; catalogue no. S957623) were obtained from AldrichCPR.

Ligands were added incrementally and absorbance spectra recorded between 350 and 500 nm after each addition. Peak-to-trough absorbance differences were plotted against ligand concentration and fitted to the Morrison equation for tight-binding ligands (99) to obtain apparent dissociation constants (*K_D_*):

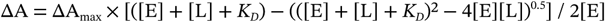

where ΔA is the observed peak-to-trough absorbance difference, ΔA_max_ the maximum absorbance change, and [E] and [L] the total enzyme and ligand concentrations. Titrations were performed in 3 independent replicates and are reported as mean ± SEM.

### Enzyme Activity Assays

Sterol substrates were prepared as stock solutions in 40% (w/v) HPCD as described above. Reactions were performed in 50 mM KPi (pH 7.4). For CYP123 assays, reaction mixtures contained 0.5 mM sterol substrate, 5 μM CYP123, 0.1 mg mL⁻¹ catalase (bovine liver catalase, Sigma-Aldrich, catalogue no. C9322) and an electron transfer system of 9 μM spinach ferredoxin (SoFdx, ferredoxin from *Spinacia oleracea*, Sigma-Aldrich, catalogue no. F3013) and 1 μM spinach ferredoxin reductase (SoFNR, ferredoxin-NADP⁺ reductase from *Spinacia oleracea*, Sigma-Aldrich, catalogue no. F0628). Stepwise oxidation assays were initiated with 10 mM NADPH and incubated at 30 °C for 0–4 h, with fresh enzyme components and cofactors added every 30 min. For CYP51–CYP123 cascade assays, 0.1 mM lanosterol, 1 mM NADPH, 0.5 U mL⁻¹ glucose-6-phosphate dehydrogenase (G6PDH), 5 mM glucose-6-phosphate (G6P) (100), 0.1 mg mL⁻¹ catalase and the SoFdx–SoFNR electron transfer system were incubated overnight at 30 °C with either 5 μM CYP51 alone (9 μM SoFdx, 1 μM SoFNR) or 5 μM each of CYP51 and CYP123 (18 μM SoFdx, 2 μM SoFNR), the redox partners being scaled with total P450. Reactions were quenched and extracted with 1,000 μL ethyl acetate containing 5 μM cholesterol as internal standard; extracts were dried by vacuum centrifugation, resuspended in 20 μL anhydrous pyridine and derivatised with 30 μL BSTFA + 1% (v/v) TMCS for 1 h at 80 °C.

### Preparative-Scale Reaction and NMR Sample Preparation

For NMR analysis, the CYP123 reaction was scaled up from 0.1 mL to 10 mL, containing 1 mg T-MAS (0.24 mM). To favour accumulation of the alcohol intermediate (T-MAS-OH), the reaction was performed under limiting conditions with 5 mM NADPH, 5 μM CYP123, 9 μM SoFdx, 1 μM SoFNR, and 0.1 mg mL⁻¹ catalase at 30 °C for 10 h. The predominant formation of T-MAS-OH under these conditions was confirmed by GC-MS (Fig. S22). The reaction was quenched and extracted with ethyl acetate (3 × 3 mL). The combined organic phases were washed with water, dried over anhydrous Na₂SO₄, and evaporated to dryness under a stream of nitrogen. T-MAS-OH was not separated from unreacted T-MAS. To preserve sample recovery and NMR signal intensity, the crude extract (a mixture of T-MAS and T-MAS-OH at approximately 2:1, estimated from the integrals of H-29 and H-19 in T-MAS-OH and H-28 in T-MAS) was dissolved in 550 μL CDCl₃ and analysed directly.

### GC-MS

Derivatised sterols were analysed on two platforms, based on a protocol described in (100). Routine assays were run on an Agilent 7890B GC coupled to a 5975 series MSD with a VF-5ht non-polar column (30 m × 0.25 mm × 0.10 μm). Accurate-mass measurements for product identification were made on an Agilent 7890B GC coupled to a 7200 Accurate-Mass Q-TOF GC/MS (Agilent) with a VF-5ms non-polar column (30 m × 0.25 mm × 0.25 μm). Ionisation was by electron impact at 70 eV with helium as carrier gas at 1 mL min⁻¹. Data were acquired in splitless mode with source and quadrupole temperatures of 230 °C and 150 °C and an injector temperature of 300 °C. The temperature programme was: 60 °C for 2 min; 50 °C min⁻¹ to 270 °C, held 8 min; 20 °C min⁻¹ to 325 °C, held 2 min.

### LC-MS/MS

Assay extracts in ethyl acetate (5 μL) were separated on an ACQUITY UPLC BEH Shield C18 column (130 Å, 1.7 μm, 2.1 mm × 150 mm) using a gradient of water and methanol, each containing 0.1% (v/v) formic acid. Mass analysis was performed on a coupled Waters Vion IMS Q-ToF in ESI-positive mode with capillary voltage 3.00 kV, source temperature 130 °C, desolvation temperature 600 °C, cone gas 35 L h⁻¹, desolvation gas 850 L h⁻¹, sampling cone 40 V, source offset 80 V and a scan range of 50–1,000 *m/z*. Free sterols ionised as [M−H₂O+H]⁺, which were selected as precursor ions; MS/MS spectra were acquired at 35 V.

### NMR

Spectra were recorded on a Bruker AVIII 500 MHz spectrometer with a ¹H/¹⁹F/¹³C/¹⁵N QCI-F cryoprobe and z-gradients, and on a Bruker 800 MHz spectrometer with a ¹H/¹³C/¹⁵N TCI cryoprobe. Approximately 1 mg of T-MAS, or of a T-MAS/T-MAS-OH mixture, was dissolved in 550 μL CDCl₃. Chemical shifts are reported in ppm and coupling constants in Hz, referenced to residual CHCl₃ in CDCl₃ (δ_H_ 7.26 ppm, δ_C_ 77.15 ppm).

### Crystallisation and Data Collection

Apo CYP123 was screened against a range of commercial crystallisation screens, and condition A9 of the HR2-130 Crystal Screen Kit (Hampton Research) was identified as a hit. Following optimisation, crystals were obtained by the sitting-drop vapour diffusion method, mixing 0.2–0.4 mM protein solution (approximately 9–18 mg mL⁻¹) with an equal volume of precipitant (0.2 M ammonium acetate, 0.1 M sodium citrate tribasic pH 5.6, 28–32% (w/v) PEG-4000). Crystals appeared within 7–14 days at 4 °C. Cryoprotection with PEG-200 resulted in a PEG molecule occupying the active site, affording the CYP123–PEG complex; perfluoropolyether was therefore used as the cryoprotectant for the apo structure. Crystals were flash-frozen in liquid nitrogen and diffraction data collected at 100 K on beamlines I03 and I04 at the Diamond Light Source.

### Structure Solution and Refinement

Reflections were processed with the xia2 pipeline (101). Structures were solved by molecular replacement using the AlphaFold-predicted (102) model of CYP123 in MOLREP (103), followed by alternating rounds of manual model building in Coot (104) and refinement in REFMAC5 (105) within CCP4i2 (106). Active-site cavity volumes were calculated with CASTp (107). Coordinates and structure factors have been deposited in the Protein Data Bank under accession codes 9TJO (apo) and 9TJP (PEG-bound).

### Molecular Docking

Docking was performed in SwissDock (Attracting Cavities 2.0) (108) using the apo CYP123 monomer as a receptor, with crystallographic waters removed. Calculations used the buried-cavity prioritisation mode with medium sampling exhaustivity. FF-MAS (4,4-dimethyl-5α-cholesta-8,14,24-trien-3β-ol) and 4α-methyl-5α-cholesta-8,14,24-trien-3β-ol (4α-Me-FF-MAS) were docked. Poses were ranked initially by the SwissDock scoring function, but because CYP123 oxidises the C4α-methyl group the binding modes used for structural analysis were selected on catalytic orientation rather than score alone: productive poses were those in which the C4α-methyl group (C-28) was directed towards the haem iron within the narrow active-site cavity. Residues within 4.5 Å of the docked ligand were taken to define the binding pocket. To assess the basis for the absence of lanosterol binding, lanosterol was modelled by superposition onto the selected pose and its additional C14-methyl group examined for steric clashes with active-site residues.

### Growth on Alternative Carbon Sources in Minimal Medium

*M. smegmatis*, *M. phlei*, *M. abscessus* and *M. bovis* var. BCG growth on alternative carbon sources was assessed in a minimal medium (MM) as described previously (22, 109–111), containing zinc sulphate (0.1 mg L⁻¹), magnesium sulphate heptahydrate (0.5 g L⁻¹), disodium phosphate (2.5 g L⁻¹), calcium chloride (0.5 mg L⁻¹), potassium dihydrogen phosphate (1 g L⁻¹), ammonium iron(II) sulphate (50 mg L⁻¹), asparagine (0.5 g L⁻¹) and vitamin B12 (10 μg mL⁻¹), with asparagine as the sole nitrogen source.

Glycerol was prepared as a 137 mM (1% v/v) stock and added to MM to a final concentration of 13.7 mM (0.1% v/v). Sterols cholesterol and lanosterol were solubilised at 10 mM in 20% (v/v) tyloxapol with continuous stirring at 90 °C.

For growth curves, log-phase cultures (100 mL) were pelleted, washed three times in PBS to remove residual glycerol, and resuspended to OD₆₀₀ ≈ 0.5. Cultures were inoculated into MM, with or without supplement, to a final OD₆₀₀ ≈ 0.025. Growth assays were conducted at 37 °C (except *M. abscessus* which was incubated at 30 °C) and 200 rpm and growth was monitored on a BMG FLUOstar plate reader over 6–14 days (assay dependent), with readings reported at 24 h intervals. Assays were performed with 3, 6 or 8 technical replicates as stated in each figure legend, and growth curves are reported as mean ± SD.

### Growth of *M. abscessus* under Transcription Assay Conditions

Sterols were dissolved in tyloxapol:ethanol (T:E, 1:1), which also served as the vehicle control. Cultures were prepared as above, washed in sterile dH₂O, and resuspended to OD₅₉₅ 0.1 in 7H9 containing 0.5% (v/v) T:E (1:1). Growth was monitored on a BMG FLUOstar plate reader at 37 °C and 200 rpm over 72 h, with OD₅₉₅ read hourly. Replicate numbers are stated in each figure legend.

### Transcription Studies

Transcription of *cyp123*, *cyp51* and the four *cyp125* paralogues of *M. abscessus* was measured in the presence of different sterols. Transcription of *cyp51* and *cyp123* was assayed in T:E, glycerol, cholesterol and lanosterol; the four *cyp125* paralogues were additionally assayed in cholestanol and cholest-5-en-3-one. *M. abscessus* was grown in 100 mL 7H9 supplemented with T:E (control), 0.2% (v/v) glycerol, or sterol. RNA was extracted between OD₅₉₅ 0.2 and 0.3 using Lysing Matrix B and TRIzol reagent. Genomic DNA was removed with a gDNA Eliminator column and the RNeasy Mini Kit, followed by TURBO DNA-free treatment (Invitrogen). cDNA was synthesised with SuperScript IV VILO reverse transcriptase.

Primers were optimised and qPCR performed on a QuantStudio 7 or StepOne Real-Time PCR System using iTaq Universal SYBR Green Supermix. Reactions contained 200 nM primers. Primer efficiencies were determined from standard curves generated with control cDNA samples corresponding to 0.039–5 ng μL⁻¹ RNA, and *C_t_* values for treated and control samples were determined at 1.25 ng μL⁻¹ RNA. Relative expression was calculated by the Pfaffl method (112), which applies efficiency-corrected *C_t_* values and so provides more accurate quantification than ΔΔ*C_t_* approaches when primer efficiencies differ. The housekeeping gene was *recA*, previously evaluated as a candidate reference gene for RT-qPCR across different carbon sources in another bacterium (113). Assays were performed in 3 biological replicates. Statistical significance between conditions was assessed by unpaired Welch’s *t*-test on individual biological replicates, which accommodates unequal variances and sample sizes. The resulting p values were adjusted by Bonferroni correction to account for multiple comparisons.

### Data Availability

All bioinformatics data files, including the SSNs and the Supplementary Data cited in the text, have been deposited in Zenodo under DOI 10.5281/zenodo.23534052. The Google Colab notebook used to run CD-HIT is archived under DOI 10.5281/zenodo.23503573. Atomic coordinates and structure factors have been deposited in the Protein Data Bank under accession codes 9TJO and 9TJP. Nanopore sequencing reads and genome assemblies for the eight *M. smegmatis* strains used in this study are available from the European Nucleotide Archive under study accession PRJEB128689. The reconstructed helper plasmid pNHEJ-recX-sacB-d1311, the *cyp123* complementation vector pCYP123-comp and the ORBIT cassette ORBIT-cat-hyg have been deposited in GenBank under accession numbers QB217605, QB217606 and QB217607, respectively. Source data for the growth curves, the enzyme titrations and the chromatographic and spectroscopic analyses are available from the corresponding author on reasonable request.

### Declaration of Generative AI Use

Generative AI was used in the preparation of this manuscript. Anthropic Claude was used to assist with drafting, editing and structuring the text, and with checking internal consistency and references. It was also used to develop analysis code and to assist with analysis and interpretation of the whole-genome sequencing data. It was not used to generate data. All analyses were inspected and all content reviewed and verified by the authors, who take full responsibility for the work.

