## Supplementary materials for "CYP123, a sterol C4-methyl oxidase, acts with CYP51 to initiate mycobacterial lanosterol catabolism"

**Dongxu Yuan,[a,b,†] Lauren R. Hatfield,[c,†] Hannah Ryan,[a,b,†] Harry Williams,[d] Mochen Dong,[e] Josie E. Parker,[d] Yi Jin,[a,b,*] and Rémi Zallot[c,*]**
[a]Manchester Institute of Biotechnology, the University of Manchester, M1 7DN, UK
[b]Department of Chemistry, School of Natural Sciences, Faculty of Science and Engineering, the University of Manchester, M13 9PL, UK
[c]School of Biological and Chemical Sciences, School of Science and Engineering, Manchester Metropolitan University, Manchester M1 5GD, UK
[d]Molecular Biosciences Division, School of Biosciences, Cardiff University, CF10 3AX, UK
[e]School of Physical, Chemical, and Environmental Sciences, Cardiff University, CF10 3AT, UK
[†]These authors contributed equally to this work.

```
tr|I7G922|I7G922_MYCS2              --MSLPTVEPVLDPYDYDFHEDPYPYYKRLRDEAPLYRNDERNFWALSRHEDVLKGFRNS  58
sp|P9WPP5|CP123_MYCTU               MTVRVGDPELVLDPYDYDFHEDPYPYYRRLRDEAPLYRNEERNFWAVSRHHDVLQGFRDS  60
sp|P63708|CP123_MYCBO               MTVRVGDPELVLDPYDYDFHEDPYPYYRRLRDEAPLYRNEERNFWAVSRHHDVLQGFRDS  60
tr|A0A0H3MB20|A0A0H3MB20_MYCBP      MTVRVGDPELVLDPYDYDFHEDPYPYYRRLRDEAPLYRNEERNFWAVSRHHDVLQGFRDS  60
                                      : :   * *****************:***********:******:***.***:***:*

tr|I7G922|I7G922_MYCS2              TALSNKHGVSLDPVSRNDEAHRVMSFLALDDPAHLRLRTLVSKGFTPRRIRELEARVTEI  118
sp|P9WPP5|CP123_MYCTU               TALSNAYGVSLDPSSRTSEAYRVMSMLAMDDPAHLRMRTLVSKGFTPRRIRELEPQVLEL  120
sp|P63708|CP123_MYCBO               TALSNAYGVSLDPSSRTSEAYRVMSMLAMDDPAHLRMRTLVSKGFTPRRIRELEPQVLEL  120
tr|A0A0H3MB20|A0A0H3MB20_MYCBP      TALSNAYGVSLDPSSRTSEAYRVMSMLAMDDPAHLRMRTLVSKGFTPRRIRELEPQVLEL  120
                                    ***** :****** **..**:****:**:*******:**************** :* *:

tr|I7G922|I7G922_MYCS2              AVQHLEIALQSDSFDFVDDFAGKLPMDVISELMGVPEEDRVRIRALADGVMHREDGVADV  178
sp|P9WPP5|CP123_MYCTU               ARIHLDSALQTESFDFVAEFAGKLPMDVISELIGVPDTDRARIRALADAVLHREDGVADV  180
sp|P63708|CP123_MYCBO               ARIHLDSALQTESFDFVAEFAGKLPMDVISELIGVPDTDRARIRALADAVLHREDGVADV  180
tr|A0A0H3MB20|A0A0H3MB20_MYCBP      ARIHLDSALQTESFDFVAEFAGKLPMDVISELIGVPDTDRARIRALADAVLHREDGVADV  180
                                    *  **: ***::***** :*************:***: **.*******.*:*********

tr|I7G922|I7G922_MYCS2              PASAIAASGELLVYYADMVKRRRRNVSDDLTSALVEAEIDGDKLTDDEIMAFLFLMVVAG  238
sp|P9WPP5|CP123_MYCTU               PPPAMAASIELMRYYADLIAEFRRRPANNLTSALLAAELDGDRLSDQEIMAFLFLMVIAG  240
sp|P63708|CP123_MYCBO               PPPAMAASIELMRYYADLIAEFRRRPANNLTSALLAAELDGDRLSDQEIMAFLFLMVIAG  240
tr|A0A0H3MB20|A0A0H3MB20_MYCBP      PPPAMAASIELMRYYADLIAEFRRRPANNLTSALLAAELDGDRLSDQEIMAFLFLMVIAG  240
                                    *  *:*** **: ****:: . **. :::*****: **:***:*:*:*********:**

tr|I7G922|I7G922_MYCS2              NETTTKLLANAAYWGFKNPDQLAPVFDDHSQIPLWVEETLRYDTSSQILARAVVEDLTFY  298
sp|P9WPP5|CP123_MYCTU               NETTTKLLANAVYWAAHHPGQLARVFADHSRIPMWVEETLRYDTSSQILARTVAHDLTLY  300
sp|P63708|CP123_MYCBO               NETTTKLLANAVYWAAHHPGQLARVFADHSRIPMWVEETLRYDTSSQILARTVAHDLTLY  300
tr|A0A0H3MB20|A0A0H3MB20_MYCBP      NETTTKLLANAVYWAAHHPGQLARVFADHSRIPMWVEETLRYDTSSQILARTVAHDLTLY  300
                                    ***********.**. ::*.*** ** ***:**:*****************:*..***:*
                                                                                 Frameshift
tr|I7G922|I7G922_MYCS2              GTTVPAGDVLVLLAGSANRDERAFDDPDEYRIGRDIGSKLVSFGSGAHFCLGAHLARMEA  358
sp|P9WPP5|CP123_MYCTU               DTTIPEGEVLLLLPGSANRDDRVFDDPDDYRIGREIGCKLVSFGSGAHFCLGAHLARMEA  360
sp|P63708|CP123_MYCBO               DTTIPEGEVLLLLPGSANRDDRVFDDPDDYRIGREIGCKLVSFGSGAHFCLGAHLARMEA  360
tr|A0A0H3MB20|A0A0H3MB20_MYCBP      DTTIPEGEVLLLLPGSANRDDRVFDDPDDYRIGREIGCKLVSFGSGAHFCLGLTWPGWKP  360
                                    .**:* *:**:** ******:*.*****:*****:**.*************      :

tr|I7G922|I7G922_MYCS2              RVALAELFKRIRGYDVDESASVRVHSSSVRGFAHLPITVEKR-----------------  400
sp|P9WPP5|CP123_MYCTU               RVALGALLRRIRNYEVDDDNVVRVHSSNVRGFAHLPISVQAR-----------------  402
sp|P63708|CP123_MYCBO               RVALGALLRRIRNYEVDDDNVVRVHSSNVRGFAHLPISVQAR-----------------  402
tr|A0A0H3MB20|A0A0H3MB20_MYCBP      GWPWARCCVGSATT---KSTTTTSCASIPATCADLPICRSACRPGNATLRTSPRPADHRR    417
                                        .            .. .   :*     *.***  .

tr|I7G922|I7G922_MYCS2              ------------------------------------------------------------  400
sp|P9WPP5|CP123_MYCTU               ------------------------------------------------------------  402
sp|P63708|CP123_MYCBO               ------------------------------------------------------------  402
tr|A0A0H3MB20|A0A0H3MB20_MYCBP      RRRIVGHRRGHRNRTRRPRVSGRLGGPPHGQVGRAGRQNPRRRWRGGSLPPRRDRSRVGE    477

tr|I7G922|I7G922_MYCS2              ------------------------------------------------------------  400
sp|P9WPP5|CP123_MYCTU               ------------------------------------------------------------  402
sp|P63708|CP123_MYCBO               ------------------------------------------------------------  402
tr|A0A0H3MB20|A0A0H3MB20_MYCBP      IVCGANGRGTRRGRTAGVQCRRHAPGTAARGQHRGLCRAGSDTPGRCQPAGHGRATGHGG    537

tr|I7G922|I7G922_MYCS2              ----    400
sp|P9WPP5|CP123_MYCTU               ----    402
sp|P63708|CP123_MYCBO               ----    402
tr|A0A0H3MB20|A0A0H3MB20_MYCBP      TPAR    541
```

**Figure S1: Comparative protein sequence alignment of CYP123 enzymes highlighting the altered C-terminus in *M. bovis var. BCG*.** Multiple sequence alignment of CYP123 enzymes from *M. smegmatis* mc$^2$155, *M. tuberculosis* H37Rv, *M. bovis* BAA-935, and *M. bovis var. BCG* Pasteur 1173P2. The sequences aligned correspond to the UniProt IDs I7G922, P9WPP5, P63708, and A0A0H3MB20, respectively. The alignment was generated and visualised using the EMBL-EBI CLUSTAL OMEGA (v1.2.4) tool. The sequence conservation among these species is notably high. Specifically, CYP123 from *M. bovis* and *M. tuberculosis* share 74.1% sequence identity with *M. smegmatis*. However, the sequence similarity drops to 49.4% between *M. bovis var. BCG* and *M. smegmatis*. This reduced conservation is attributed to a single-nucleotide deletion, which leads to a frameshift (from position 352). This frameshift is expected to produce a different C-terminus (highlighted and in bold in the alignment) and is proposed to result in an inactive enzyme form.

```
CP001663.1:5928894-5930096     ------ATGAGTTTGCCGACCGTCGAACCCGTACTCGACCCGTACGACTACGACTTCCAC     54
AL123456.3:c860072-858864      ATGACCGTCCGCGTCGGTGACCCCGAACTGGTCCTGGACCCCTACGACTACGACTTTCAC     60
LT708304.1:c862261-861053      ATGACCGTCCGCGTCGGTGACCCCGAACTGGTCCTGGACCCCTACGACTACGACTTTCAC     60
AM408590.1:c891853-890228      ATGACCGTCCGCGTCGGTGACCCCGAACTGGTCCTGGACCCCTACGACTACGACTTTCAC     60
                                      *  *  *      *  *****  ** ** ***** ************** ***

CP001663.1:5928894-5930096     GAGGATCCGTATCCGTACTACAAGCGACTTCGCGACGAGGCGCCGCTGTACCGCAACGAC     114
AL123456.3:c860072-858864      GAAGACCCGTACCCGTATTATCGTCGGCTGCGGGACGAGGCCCCGCTGTACCGTAACGAG     120
LT708304.1:c862261-861053      GAAGACCCGTACCCGTATTATCGTCGGCTGCGGGACGAGGCCCCGCTGTACCGTAACGAG     120
AM408590.1:c891853-890228      GAAGACCCGTACCCGTATTATCGTCGGCTGCGGGACGAGGCCCCGCTGTACCGTAACGAG     120
                               ** ** ***** ***** **    ** ** ** ******** *********** *****

CP001663.1:5928894-5930096     GAACGCAACTTCTGGGCGCTGTCGCGCCACGAGGACGTGCTCAAGGGATTCCGCAACAGC     174
AL123456.3:c860072-858864      GAACGCAATTTCTGGGCGGTGTCGCGGCACCACGACGTGCTGCAAGGCTTCCGGGACAGC     180
LT708304.1:c862261-861053      GAACGCAATTTCTGGGCGGTGTCGCGGCACCACGACGTGCTGCAAGGCTTCCGGGACAGC     180
AM408590.1:c891853-890228      GAACGCAATTTCTGGGCGGTGTCGCGGCACCACGACGTGCTGCAAGGCTTCCGGGACAGC     180
                               ******** ********* ******* *** * ********  * ** *****  *****

CP001663.1:5928894-5930096     ACCGCGCTGTCCAACAAGCACGGTGTGTCGCTGGATCCGGTGTCGCGCAACGACGAGGCG     234
AL123456.3:c860072-858864      ACGGCGTTGTCGAATGCCTATGGGGTATCCCTGGATCCGTCCTCACGCACTTCTGAGGCG     240
LT708304.1:c862261-861053      ACGGCGTTGTCGAATGCCTATGGGGTATCCCTGGATCCGTCCTCACGCACTTCTGAGGCG     240
AM408590.1:c891853-890228      ACGGCGTTGTCGAATGCCTATGGGGTATCCCTGGATCCGTCCTCACGCACTTCTGAGGCG     240
                               ** *** **** **      * ** ** ** *********   ** ****     ******

CP001663.1:5928894-5930096     CATCGCGTGATGTCGTTTCTCGCACTCGACGATCCGGCACACCTGCGGTTGCGCACGCTG     294
AL123456.3:c860072-858864      TACCGGGTAATGTCGATGCTGGCCATGGACGACCCCGCACATCTGCGGATGCGTACCTTG     300
LT708304.1:c862261-861053      TACCGGGTAATGTCGATGCTGGCCATGGACGACCCCGCACATCTGCGGATGCGTACCTTG     300
AM408590.1:c891853-890228      TACCGGGTAATGTCGATGCTGGCCATGGACGACCCCGCACATCTGCGGATGCGTACCTTG     300
                                * ** ** ****** * ** **  * ***** ** ***** ****** **** **  **

CP001663.1:5928894-5930096     GTGTCCAAGGGTTTCACGCCGCGGCGCATCCGGGAGCTGGAGGCGCGGGTCACCGAGATC     354
AL123456.3:c860072-858864      GTGTCCAAGGGCTTCACCCCACGGCGGATCCGTGAACTCGAGCCGCAGGTGCTCGAACTT     360
LT708304.1:c862261-861053      GTGTCCAAGGGCTTCACCCCACGGCGGATCCGTGAACTCGAGCCGCAGGTGCTCGAACTT     360
AM408590.1:c891853-890228      GTGTCCAAGGGCTTCACCCCACGGCGGATCCGTGAACTCGAGCCGCAGGTGCTCGAACTT     360
                               *********** ***** ** ***** ***** ** ** *** *** ***   ***  *

CP001663.1:5928894-5930096     GCCGTGCAGCATCTCGAAATTGCCCTGCAGTCGGACAGTTTCGACTTCGTCGACGACTTC     414
AL123456.3:c860072-858864      GCCCGCATTCACCTGGATTCGGCCCTGCAAACCGAAAGTTTCGATTTCGTAGCAGAATTC     420
LT708304.1:c862261-861053      GCCCGCATTCACCTGGATTCGGCCCTGCAAACCGAAAGTTTCGATTTCGTAGCAGAATTC     420
AM408590.1:c891853-890228      GCCCGCATTCACCTGGATTCGGCCCTGCAAACCGAAAGTTTCGATTTCGTAGCAGAATTC     420
                               ***      ** ** **    ********  * ** ******** ***** *  ** ***

CP001663.1:5928894-5930096     GCGGGCAAGCTTCCGATGGACGTCATCTCCGAGCTCATGGGCGTCCCCGAAGAGGACCGC     474
AL123456.3:c860072-858864      GCTGGCAAGCTGCCGATGGATGTGATTTCAGAGCTGATAGGCGTGCCCGACACCGACCGG     480
LT708304.1:c862261-861053      GCTGGCAAGCTGCCGATGGATGTGATTTCAGAGCTGATAGGCGTGCCCGACACCGACCGG     480
AM408590.1:c891853-890228      GCTGGCAAGCTGCCGATGGATGTGATTTCAGAGCTGATAGGCGTGCCCGACACCGACCGG     480
                               ** ******** ******** ** ** ** ***** ** ***** *****    *****

CP001663.1:5928894-5930096     GTGCGGATCCGTGCGCTGGCCGACGGTGTCATGCACCGTGAGGACGGCGTGGCCGACGTG     534
AL123456.3:c860072-858864      GCCCGCATCCGCGCGCTGGCCGATGCGGTGCTGCACCGCGAGGACGGCGTGGCCGATGTG     540
LT708304.1:c862261-861053      GCCCGCATCCGCGCGCTGGCCGATGCGGTGCTGCACCGCGAGGACGGCGTGGCCGATGTG     540
AM408590.1:c891853-890228      GCCCGCATCCGCGCGCTGGCCGATGCGGTGCTGCACCGCGAGGACGGCGTGGCCGATGTG     540
                               *  ** ***** *********** *  ** ******* **************** ***

CP001663.1:5928894-5930096     CCGGCCTCGGCCATCGCGGCCTCCGGTGAACTGCTCGTGTACTACGCCGACATGGTCAAG     594
AL123456.3:c860072-858864      CCCCCGCCGGCGATGGCGGCGTCGATCGAGCTGATGAGATATTACGCTGACCTGATCGCG     600
LT708304.1:c862261-861053      CCCCCGCCGGCGATGGCGGCGTCGATCGAGCTGATGAGATATTACGCTGACCTGATCGCG     600
AM408590.1:c891853-890228      CCCCCGCCGGCGATGGCGGCGTCGATCGAGCTGATGAGATATTACGCTGACCTGATCGCG     600
                               **  * **** ** ***** **    ** *** *    ** ***** *** ** **  *

CP001663.1:5928894-5930096     AGGCGTCGCCGCAACGTCAGCGACGACCTGACCTCGGCGCTGGTCGAGGCCGAGATCGAC     654
AL123456.3:c860072-858864      GAATTCCGGCGGCGGCCCGCGAACAATCTGACGTCGGCACTGCTGGCAGCCGAGCTCGAC     660
LT708304.1:c862261-861053      GAATTCCGGCGGCGGCCCGCGAACAATCTGACGTCGGCACTGCTGGCAGCCGAGCTCGAC     660
AM408590.1:c891853-890228      GAATTCCGGCGGCGGCCCGCGAACAATCTGACGTCGGCACTGCTGGCAGCCGAGCTCGAC     660
                                     ** **      *    ** * ***** ***** *** * *  ****** *****

CP001663.1:5928894-5930096     GGTGACAAGCTCACCGACGACGAGATCATGGCGTTCCTGTTCCTCATGGTCGTCGCCGGT     714
AL123456.3:c860072-858864      GGCGACCGGCTTTCCGACCAGGAAATCATGGCGTTCCTGTTTCTCATGGTGATCGCCGGC     720
LT708304.1:c862261-861053      GGCGACCGGCTTTCCGACCAGGAAATCATGGCGTTCCTGTTTCTCATGGTGATCGCCGGC     720
AM408590.1:c891853-890228      GGCGACCGGCTTTCCGACCAGGAAATCATGGCGTTCCTGTTTCTCATGGTGATCGCCGGC     720
                               ** ***  ***  ***** * ** ***************** ******** *******

CP001663.1:5928894-5930096     AACGAGACCACCACCAAACTCCTTGCCAATGCCGCCTATTGGGGTTTCAAGAACCCCGAT     774
AL123456.3:c860072-858864      AACGAGACCACCACCAAGCTACTGGCCAATGCCGTCTACTGGGCCGCCCACCACCCTGGC     780
LT708304.1:c862261-861053      AACGAGACCACCACCAAGCTACTGGCCAATGCCGTCTACTGGGCCGCCCACCACCCTGGC     780
AM408590.1:c891853-890228      AACGAGACCACCACCAAGCTACTGGCCAATGCCGTCTACTGGGCCGCCCACCACCCTGGC     780
                               ***************** ** ** ********** *** ****    * *  **** *

CP001663.1:5928894-5930096     CAGCTGGCGCCGGTGTTCGACGACCACAGCCAGATCCCCCTGTGGGTCGAGGAGACGCTG     834
AL123456.3:c860072-858864      CAGCTGGCCCGCGTATTCGCCGACCACTCCCGGATTCCGATGTGGGTGGAGGAAACCCTG     840
LT708304.1:c862261-861053      CAGCTGGCCCGCGTATTCGCCGACCACTCCCGGATTCCGATGTGGGTGGAGGAAACCCTG     840
AM408590.1:c891853-890228      CAGCTGGCCCGCGTATTCGCCGACCACTCCCGGATTCCGATGTGGGTGGAGGAAACCCTG     840
                               ******** *  ** **** *******  ** *** **  ******* ***** ** ***

CP001663.1:5928894-5930096     CGCTACGACACCTCCAGCCAGATCCTCGCGCGTGCGGTGGTGGAAGATTTGACGTTCTAC     894
AL123456.3:c860072-858864      CGCTACGACACGTCCAGCCAGATTCTGGCCCGCACCGTCGCGCACGATCTCACGTTGTAC     900
```

```
LT708304.1:c862261-861053       CGCTACGACACGTCCAGCCAGATTCTGGCCCGCACCGTCGCGCACGATCTCACGTTGTAC     900
AM408590.1:c891853-890228       CGCTACGACACGTCCAGCCAGATTCTGGCCCGCACCGTCGCGCACGATCTCACGTTGTAC     900
                                *********** *********** ** ** **  * ** * * * *** * ***** ***

CP001663.1:5928894-5930096      GGCACGACGGTGCCCGCGGGTGACGTGCTGGTGCTGCTGGCCGGTTCGGCCAACCGTGAC     954
AL123456.3:c860072-858864       GACACCACGATCCCCGAGGGTGAGGTGTTGCTGCTGCTACCGGGATCGGCCAACCGTGAC     960
LT708304.1:c862261-861053       GACACCACGATCCCCGAGGGTGAGGTGTTGCTGCTGCTACCGGGATCGGCCAACCGTGAC     960
AM408590.1:c891853-890228       GACACCACGATCCCCGAGGGTGAGGTGTTGCTGCTGCTACCGGGATCGGCCAACCGTGAC     960
                                * *** *** * **** ****** *** ** *******  * ** ***************

CP001663.1:5928894-5930096      GAGCGGGCGTTCGACGACCCCGACGAGTACCGCATCGGCCGTGACATCGGTTCCAAGCTC     1014
AL123456.3:c860072-858864       GACCGGGTGTTCGACGACCCGGACGACTATCGCATCGGCCGCGAAATCGGCTGCAAACTA     1020
LT708304.1:c862261-861053       GACCGGGTGTTCGACGACCCGGACGACTATCGCATCGGCCGCGAAATCGGCTGCAAACTA     1020
AM408590.1:c891853-890228       GACCGGGTGTTCGACGACCCGGACGACTATCGCATCGGCCGCGAAATCGGCTGCAAACTA     1020
                                ** **** ************ ***** ** *********** ** ***** * *** **
                                                                  Deletion
CP001663.1:5928894-5930096      GTCAGTTTCGGCAGCGGTGCCCATTTCTGCCTCGGCGCGCATCTGGCGCGTATGGAGGCC     1074
AL123456.3:c860072-858864       GTCAGTTTCGGCAGCGGTGCCCACTTCTGTCTGGGGGCTCACCTGGCCCGGATGGAAGCC     1080
LT708304.1:c862261-861053       GTCAGTTTCGGCAGCGGTGCCCACTTCTGTCTGGGGGCTCACCTGGCCCGGATGGAAGCC     1080
AM408590.1:c891853-890228       GTCAGTTTCGGCAGCGGTGCCCACTTCTGTCTGGG-GCTCACCTGGCCCGGATGGAAGCC     1079
                                ********************** ***** ** ** ** ** ***** ** ***** ***

CP001663.1:5928894-5930096      AGGGTCGCGCTGGCCGAGTTGTTCAAGCGGATCCGCGGATACGACGTGGACGAATCCGCG     1134
AL123456.3:c860072-858864       CGGGTGGCCCTGGGCGCGCTGCTGCGTCGGATCCGCAACTACGAAGTCGACGACGACAAC     1140
LT708304.1:c862261-861053       CGGGTGGCCCTGGGCGCGCTGCTGCGTCGGATCCGCAACTACGAAGTCGACGACGACAAC     1140
AM408590.1:c891853-890228       CGGGTGGCCCTGGGCGCGCTGCTGCGTCGGATCCGCAACTACGAAGTCGACGACGACAAC     1139
                                 **** ** **** ** * ** *   *********  ***** ** *****  *

CP001663.1:5928894-5930096      TCCGTGCGGGTGCATTCCAGCAGCGTCCGCGGTTTCGCTCACCTTCCGATCACAGTGGAG     1194
AL123456.3:c860072-858864       GTCGTGCGCGTCCATTCCAGCAACGTGCGCGGATTTGCCCATCTGCCGATCAGCGTGCAG     1200
LT708304.1:c862261-861053       GTCGTGCGCGTCCATTCCAGCAACGTGCGCGGATTTGCCCATCTGCCGATCAGCGTGCAG     1200
AM408590.1:c891853-890228       GTCGTGCGCGTCCATTCCAGCAACGTGCGCGGATTTGCCCATCTGCCGATCAGCGTGCAG     1199
                                  ****** ** ********** *** ***** ** ** ** ** *******  *** **

CP001663.1:5928894-5930096      AAACGCTAA---------------------------------------------------     1203
AL123456.3:c860072-858864       GCCAGGTAA---------------------------------------------------     1209
LT708304.1:c862261-861053       GCCAGGTAA---------------------------------------------------     1209
AM408590.1:c891853-890228       GCCAGGTAATGCCACGCTTCGAACCTCACCCCGCCCGGCGGACCACCGTCGTCGCCGGCG     1259
                                    * ***

CP001663.1:5928894-5930096      ------------------------------------------------------------     1203
AL123456.3:c860072-858864       ------------------------------------------------------------     1209
LT708304.1:c862261-861053       ------------------------------------------------------------     1209
AM408590.1:c891853-890228       CATCGTCGGGCATCGGCGCGGCCACCGCAACCGAACTCGCCGGCCGCGGGTTTCCGGTCG     1319

CP001663.1:5928894-5930096      ------------------------------------------------------------     1203
AL123456.3:c860072-858864       ------------------------------------------------------------     1209
LT708304.1:c862261-861053       ------------------------------------------------------------     1209
AM408590.1:c891853-890228       CCTTGGGGGCCCGCCGCATGGACAAGTTGGCCGAGCTGGTCGACAAAATCCGCGCCGACG     1379

CP001663.1:5928894-5930096      ------------------------------------------------------------     1203
AL123456.3:c860072-858864       ------------------------------------------------------------     1209
LT708304.1:c862261-861053       ------------------------------------------------------------     1209
AM408590.1:c891853-890228       GTGGCGAGGCGGTAGCCTTCCCCCTCGACGTGACCGATCCCGAGTCGGTGAAATCGTTTG     1439

CP001663.1:5928894-5930096      ------------------------------------------------------------     1203
AL123456.3:c860072-858864       ------------------------------------------------------------     1209
LT708304.1:c862261-861053       ------------------------------------------------------------     1209
AM408590.1:c891853-890228       TGGCGCAAACGGTCGAGGCACTCGGCGAGGTCGAACTGCTGGTGTCCAGTGCAGGCGACA     1499

CP001663.1:5928894-5930096      ------------------------------------------------------------     1203
AL123456.3:c860072-858864       ------------------------------------------------------------     1209
LT708304.1:c862261-861053       ------------------------------------------------------------     1209
AM408590.1:c891853-890228       TGCTCCCGGGACAGCTGCACGAGGTCAGCACCGAGGCCTTTGCAGAGCAGGTTCAGATAC     1559

CP001663.1:5928894-5930096      ------------------------------------------------------------     1203
AL123456.3:c860072-858864       ------------------------------------------------------------     1209
LT708304.1:c862261-861053       ------------------------------------------------------------     1209
AM408590.1:c891853-890228       ACCTGGTCGGTGCCAACCGGCTGGCCACGGCCGTGCTACCGGCCATGGTGGCACGCCGGC     1619

CP001663.1:5928894-5930096      -------     1203
AL123456.3:c860072-858864       -------     1209
LT708304.1:c862261-861053       -------     1209
AM408590.1:c891853-890228       GAGGTGA     1626
```

**Figure S2: Comparative nucleotide sequence alignment of CYP123 genes highlighting the frameshift mutation in *M. bovis var. BCG*.** Multiple sequence alignment of the genes encoding CYP123 enzymes from *M. smegmatis* mc$^{2}$155, *M. tuberculosis H37Rv, M. bovis BAA-935, and M. bovis var. BCG Pasteur* 1173P2. The sequences were obtained from genomes CP001663, AL123456, LT708304, and AM408590, corresponding to the UniProt IDs I7G922, P9WPP5, P63708, and A0A0H3MB20, respectively. The alignment was generated and visualised using the EMBL-EBI CLUSTAL OMEGA (v1.2.4) tool. In the sequence encoding CYP123 in *M. bovis var. BCG*, a single deletion mutation is observed (highlighted in yellow, positioned just after nucleotide 1055). This mutation is proposed to lead to a frameshift, resulting in a protein with an alternate C-terminus compared to other species, potentially leading to an inactive enzyme.

**Table S1: Co-occurrence of gene neighbours of CYP123 across 382 CYP123-encoding genomes in *Mycobacteriaceae*.** Gene neighbours of sequences from Cluster 15 (CYP123) in the *Mycobacteriaceae* PF00067 SSN at AST 131 (Fig. 1A). Co-occurrence was calculated from the Cluster Centre Genome Neighbourhood Network generated by the EFI Genome Neighbourhood Tool with a window size of ±5 genes. Only neighbours present in ≥50% of queried genomes are shown.

| Pfam | Pfam Description - Note | # of Queries with Pfam Neighbours | Co-occurrence Ratio | Median Distance | Co-occurrence |
|---|---|---|---|---|---|
| **Same orientation as CYP123** | | | | | |
| PF00440 | Bacterial regulatory proteins, tetR family | 376 | 376/382 | 1 | 0.98 |
| **PF00067** | **Cytochrome P450 - CYP123** | | **Query** | **0** | **CYP123** |
| PF00106 | short chain dehydrogenase | 370 | 370/382 | 1 | 0.96 |
| PF00067 | Cytochrome P450 - CYP51 | 367 | 367/382 | 2 | 0.96 |
| PF13459 | 4Fe-4S single cluster domain - Ferredoxin | 365 | 365/382 | 3 | 0.95 |
| none | Nuclear transport factor 2 family protein IPR032710 (NTF2) - No PFAM assigned | 360 | 360/382 | 4 | 0.94 |
| **Opposite orientation to CYP123** | | | | | |
| PF00171 | Aldehyde dehydrogenase family | 374 | 374/382 | 2 | 0.97 |
| PF13561 | Enoyl-(Acyl carrier protein) reductase | 365 | 365/382 | 3 | 0.95 |
| PF03446-PF14833 | NAD binding domain of 6-phosphogluconate dehydrogenase-NAD-binding of NADP-dependent 3-hydroxyisobutyrate dehydrogenase | 364 | 364/382 | 4 | 0.95 |
| PF02627 | Carboxymuconolactone decarboxylase family | 358 | 358/382 | 5 | 0.93 |

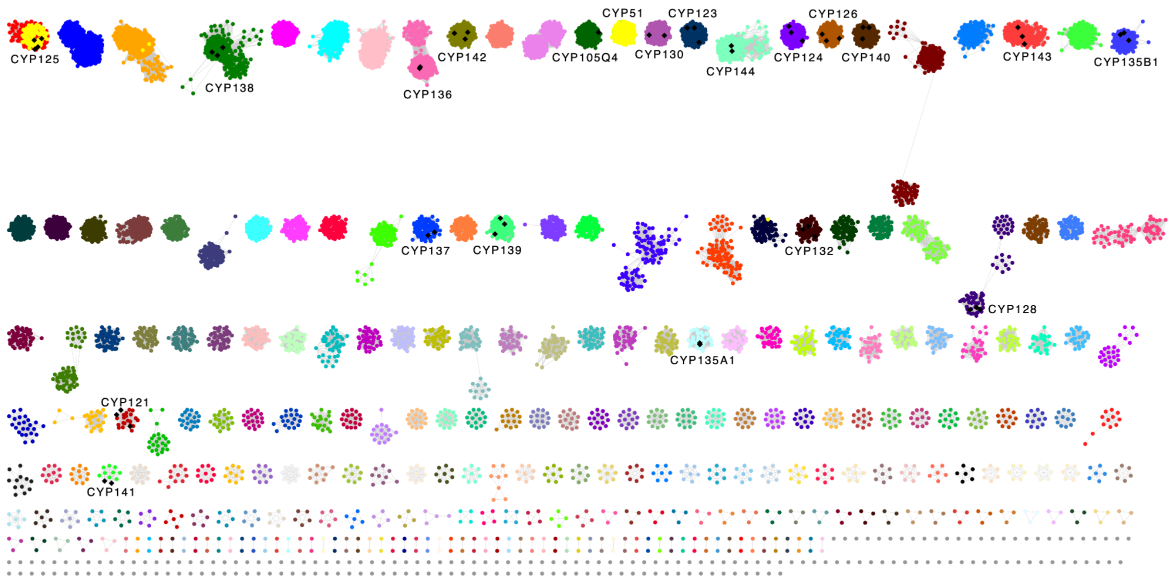


**Figure S3: Cytochrome P450 sequences neighbouring CYP123 in *Mycobacteriaceae* genomes.** The *Mycobacteriaceae* P450 SSN from Fig. 1A is shown with P450 sequences located within 10 genes on each side of CYP123 (Cluster 15) highlighted in yellow. CYP51 cluster members are identified as neighbours, consistent with their presence in the same operonic structure. Of the 383 CYP123 sequences queried, 156 have a CYP125 cluster member within the 10-gene window, indicating that CYP125 is frequently co-localised with the CYP123–CYP51 region. No CYP142 cluster members are found neighbouring CYP123-encoding genes. Cluster labels indicate characterised CYP subfamilies; unlabelled clusters contain sequences with no experimentally assigned function.

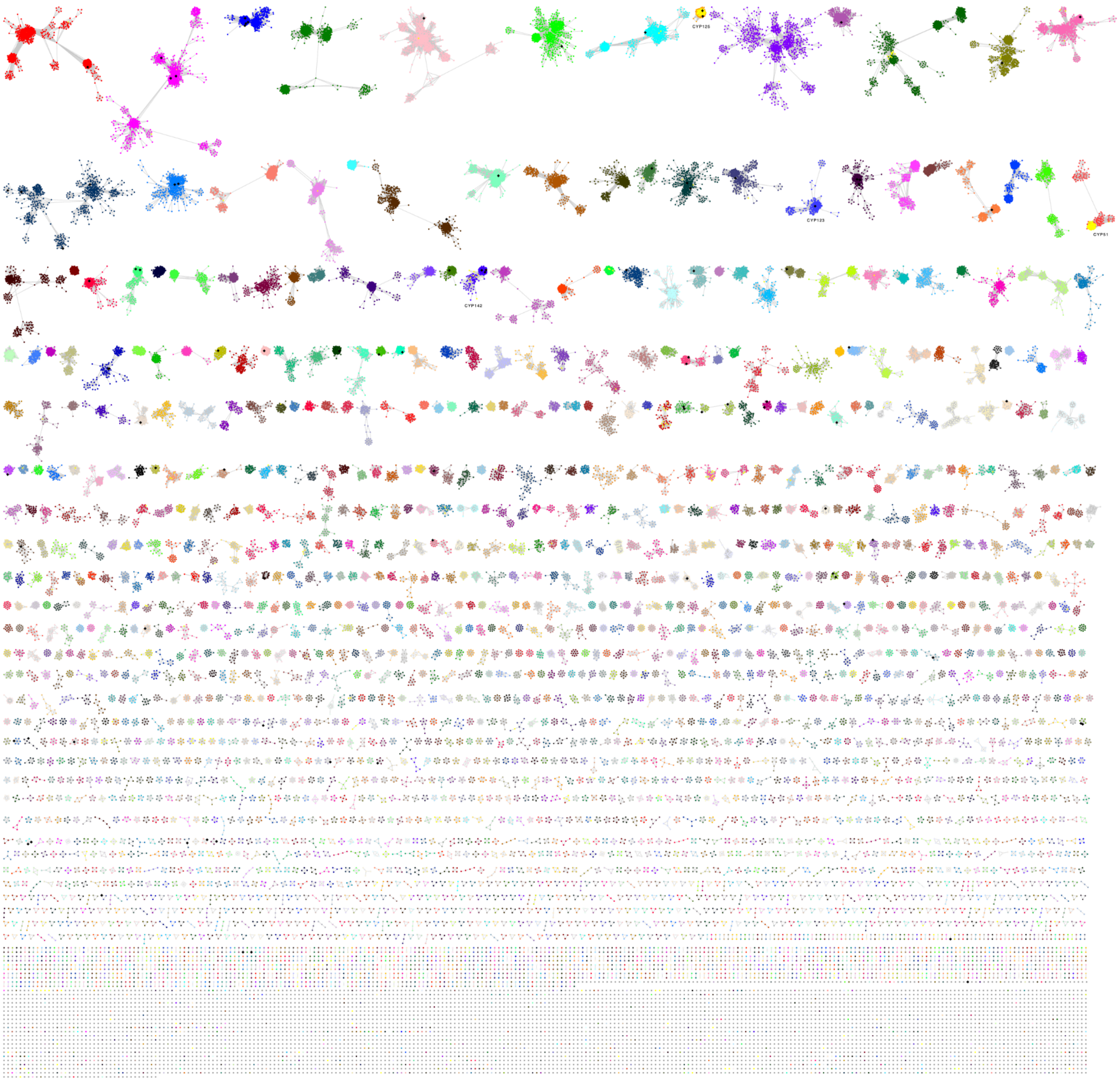


**Figure S4: Distribution of CYP123 across bacteria.** SSN of UniRef90 sequences belonging to PF00067 from bacteria, generated at an alignment score threshold of 126 (55% sequence identity). Cluster 24 (CYP123) comprises 389 UniRef90 sequences representing 902 UniProt sequences, distributed across the *Actinomycetales*, including *Rhodococcus*, *Nocardioides*, *Gordonia*, *Actinomadura*, *Dietzia*, *Nocardia* and *Prescottella*, and across *Saccharopolyspora*.

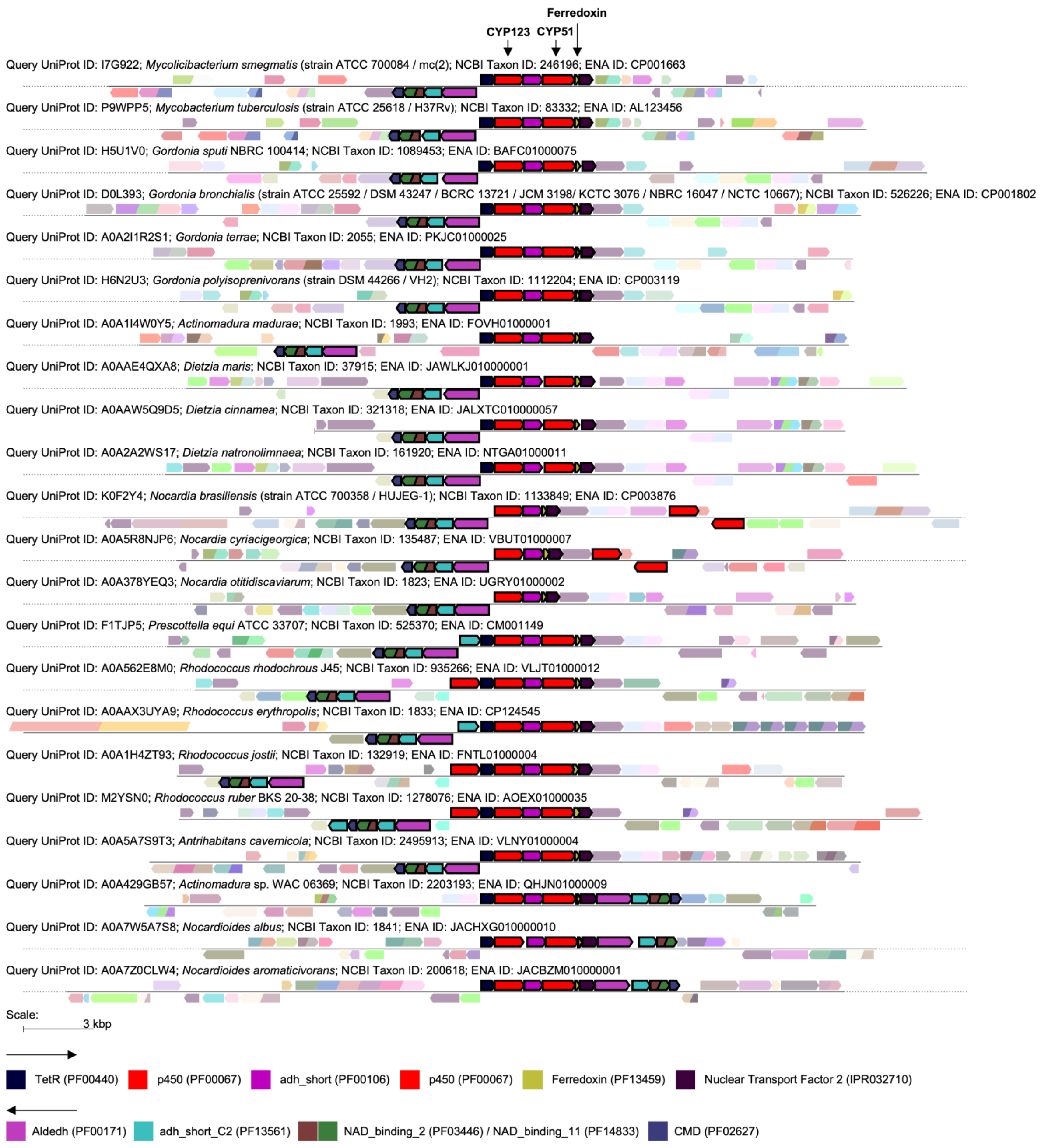


**Figure S5: Genome Neighbourhood Diagrams for bacterial CYP123 homologues.** Genome Neighbourhood Diagrams generated by EFI-GNT for all sequences in the CYP123 cluster (Cluster 24 from Fig. S4), displaying the genomic context (±10 genes) of CYP123 homologues across bacteria. Genes are coloured by Pfam/InterPro family assignment (key at bottom).

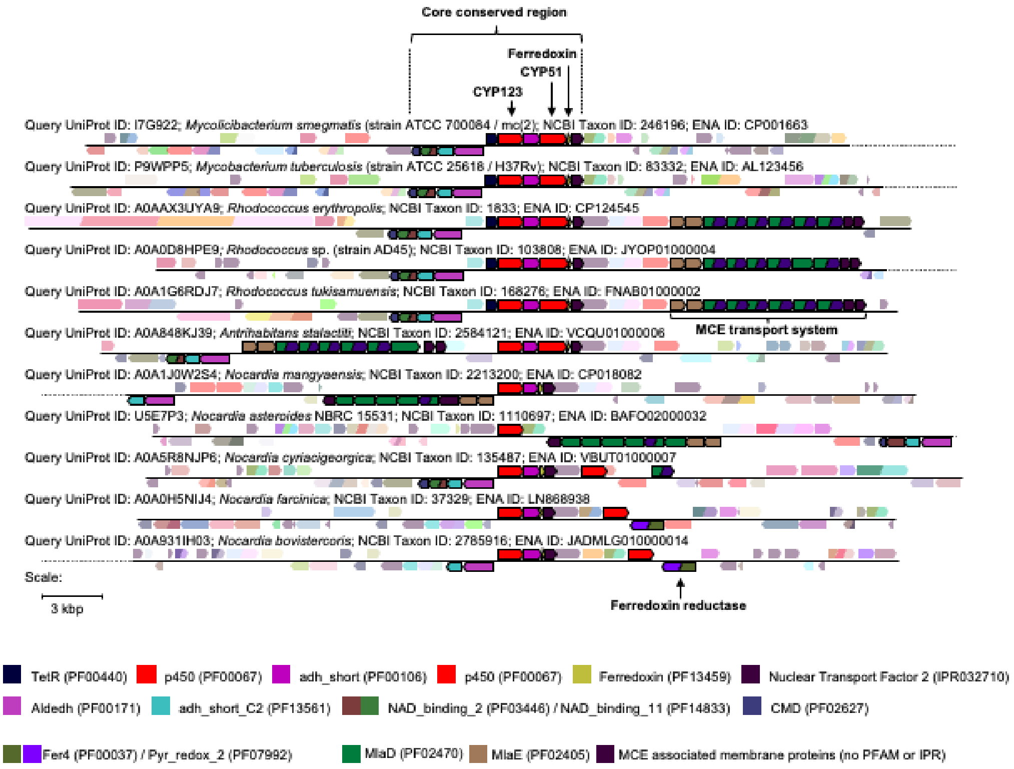


**Figure S6: Conservation of the CYP123–CYP51 genomic region in actinobacteria.** Annotated Genome Neighbourhood Diagrams for 11 representative species: *M. smegmatis* and *M. tuberculosis* (*Mycobacteriaceae* references), *Rhodococcus* erythropolis, *Rhodococcus* sp. AD45, R. tukisamuensis, Antrihabitans stalactiti, *Nocardia* mangyaensis, N. asteroides, N. cyriacigeorgica, N. farcinica, and N. bovistercoris. The bracket marks the core conserved region spanning CYP123, CYP51, and the adjacent ferredoxin. In A. stalactiti and several *Nocardia* species, genes encoding components of the Mce4 transport system, which mediates sterol import in mycobacteria (https://doi.org/10.7554/eLife.26969), are found adjacent to the core region. In N. bovistercoris, a ferredoxin reductase gene is located near the conserved block. Genes are shown as arrows indicating transcriptional orientation, coloured by Pfam/InterPro family assignment (key at bottom). Scale bar: 3 kbp.

**Table S2: Statistical analysis of differential gene expression.** P values from unpaired Welch's t-tests on individual biological replicates, comparing expression of each gene under the indicated growth conditions with the tyloxapol:ethanol (T:E) vehicle control, with Bonferroni correction for multiple comparisons; corrected values are capped at 1. Values <0.05 denote statistically significant differences. A dash indicates a condition that was not tested for that gene.

| | Gene identifier (MAB_x) | | | | | |
|---|---|---|---|---|---|---|
| **Condition** | **0101** | **0611** | **0613** | **1211c** | **1214c** | **1216c** |
| **Glycerol** | 1.0000 | 1.0000 | 0.1829 | 1.0000 | 0.0070 | 0.0523 |
| **Cholesterol** | 0.0721 | 0.0230 | 0.2130 | 0.6506 | 1.0000 | 1.0000 |
| **Lanosterol** | 1.0000 | 0.0674 | 0.0837 | 0.0353 | 0.0104 | 0.0056 |
| **Cholestanol** | 0.6779 | 0.0886 | 0.1292 | 0.1218 | - | - |
| **Cholest-5-en-3-one** | 0.5135 | 0.0715 | 0.2382 | 0.5369 | - | - |

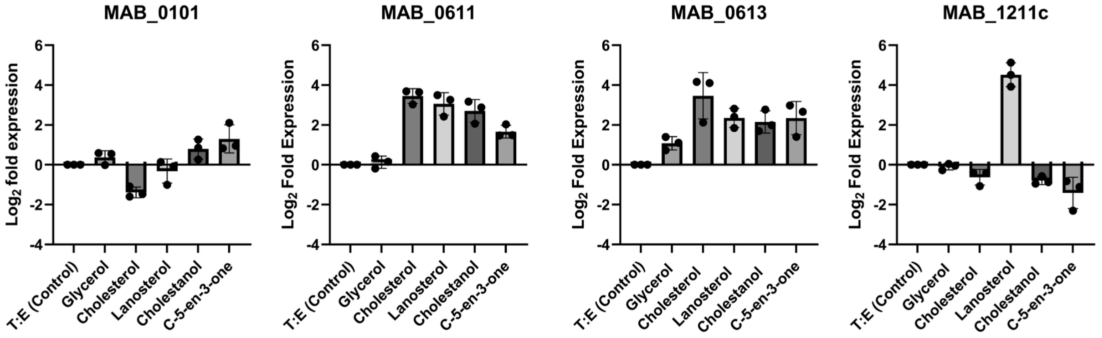


**Figure S7: Sterol responsiveness of CYP125 paralogues in *M. abscessus*.** Relative expression of the four CYP125 paralogues (MAB_0101, MAB_0611, MAB_0613, and MAB_1211c) measured by qRT-PCR in *M. abscessus* cultured in 7H9 medium supplemented with T:E (control), glycerol (0.2%), cholesterol (0.5 mM), lanosterol (1 mM), cholestanol (0.5 mM), or cholest-5-en-3-one (1 mM). RNA was harvested at mid-log phase. Expression is shown as $\log_2$ fold change relative to the T:E control, calculated using the Pfaffl method and normalised to *recA*. MAB_0611 and MAB_0613 respond to multiple sterols, though only the induction of MAB_0611 by cholesterol survives correction for multiple comparisons, whereas MAB_1211c is selectively induced by lanosterol, consistent with co-regulation within the CYP51–CYP123 operon (Fig. 2). MAB_0101 shows weak, non-significant responses. The cholesterol utilisation cluster that contains MAB_0611 and MAB_0613 carries the ring-degrading 3-oxosteroid 1-dehydrogenase together with the thiolase, acyl-CoA dehydrogenase and enoyl-CoA hydratase genes of side-chain β-oxidation. Bars represent mean expression from three biological replicates with individual replicate values overlaid as dots; error bars represent standard error of the mean.

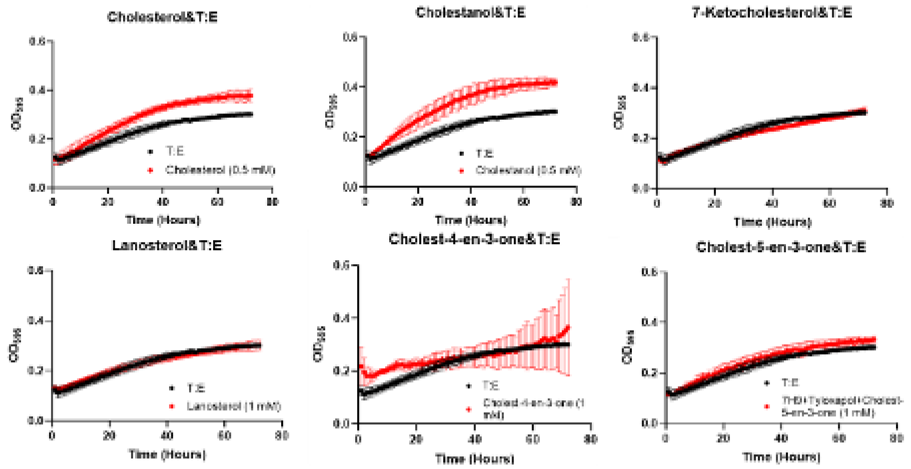


**Figure S8: Growth of *M. abscessus* on six different sterols vs growth on T:E (control).** Growth curves of *M. abscessus* cultured in 7H9 medium supplemented with 0.5% tyloxapol:ethanol (1:1) and individual sterols as potential carbon sources. Growth was monitored at $OD_{595}$ over 72 hours using a BMG FLUOstar microplate reader. The following conditions were tested; cholesterol (0.5 mM), cholestanol (0.5 mM), 7-ketocholesterol (0.5 mM), lanosterol (1 mM), cholest-4-en-3-one (1 mM), and cholest-5-en-3-one (1 mM). Black dots represent control cultures without added sterol (T:E only), and red dots represent cultures supplemented with the indicated sterol. Data points show mean values of three biological replicates with error bars representing standard error.

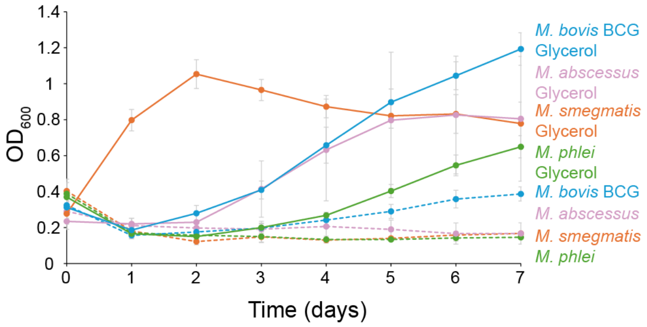


**Figure S9: Glycerol-supported growth of mycobacterial species in minimal medium.** Growth of *M. smegmatis* (orange), *M. phlei* (green), *M. abscessus* (pink) and *M. bovis var. BCG* (blue) in minimal medium without supplementation (dashed) and supplemented with 0.1% (v/v) glycerol (solid) as the sole carbon source. $OD_{600}$ was measured every 24 h; data represent the mean ± SD (M. smegmatis with and without glycerol; n=6; *M. phlei* with (n=8) and without (n=6) glycerol; *M. abscessus* with (n=8) and without (n=6) glycerol; *M. bovis* var. BCG with (n=8) and without (n=6) glycerol).

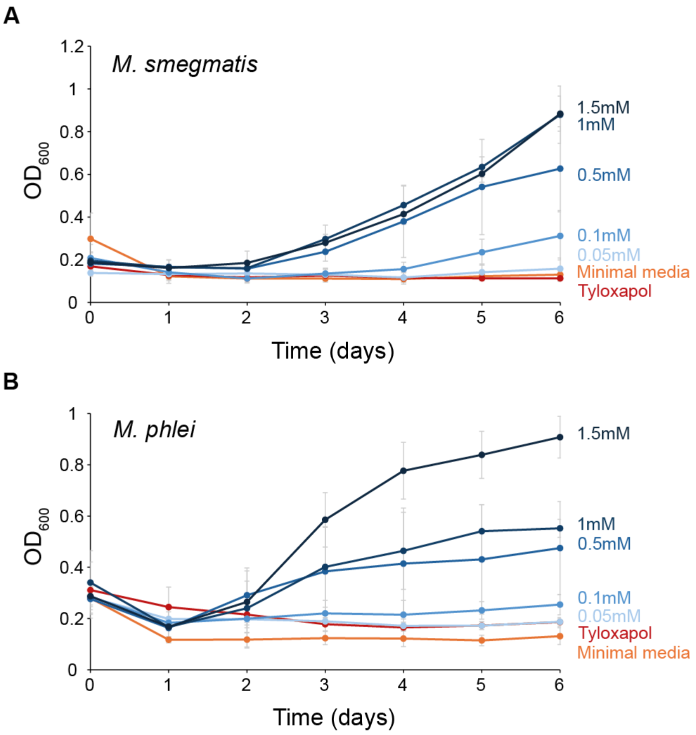


**Figure S10: Dose-dependent growth of mycobacteria on cholesterol.** Growth of *M. smegmatis* **(A)** and *M. phlei* **(B)** in minimal medium supplemented with cholesterol (0.05–1.5 mM) as the sole carbon source. $OD_{600}$ was measured every 24 h; data represent the mean ± SD of three technical replicates.

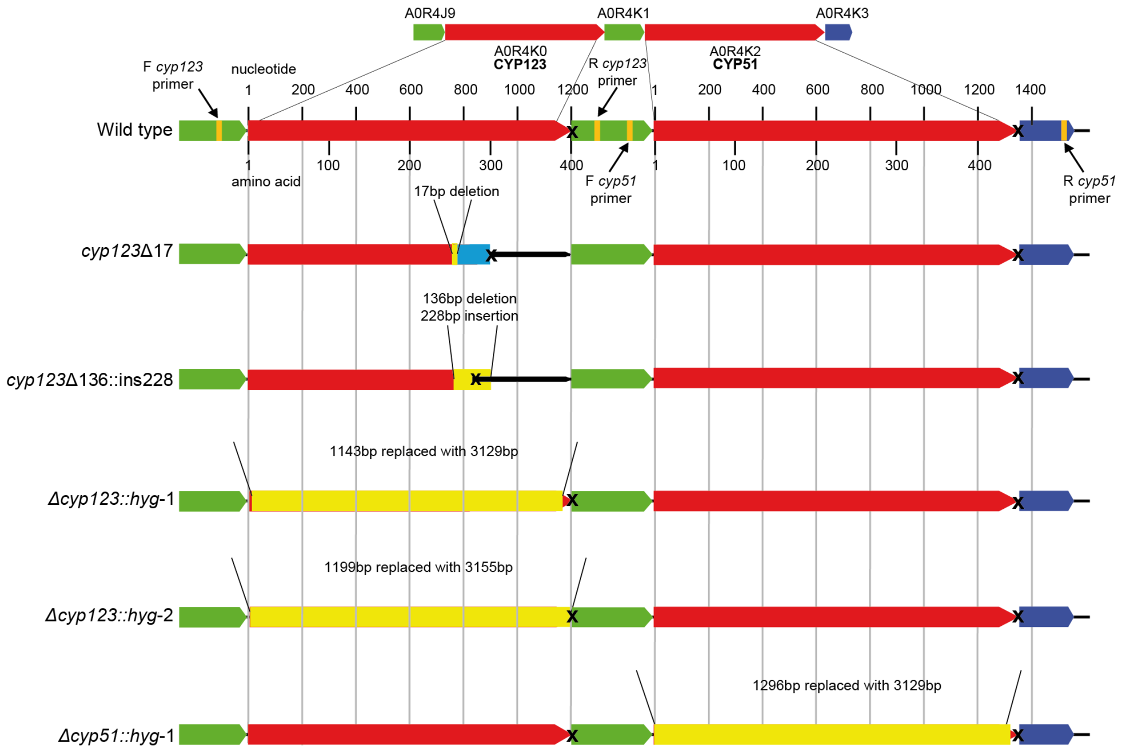


**Figure S11: CRISPR-Cas and ORBIT editing of the *M. smegmatis cyp123* gene.** The *M. smegmatis cyp123* gene was targeted by CRISPR-Cas gene editing at position 762. *M. smegmatis cyp123* and *cyp51* were also targeted by ORBIT gene editing. CRISPR-Cas and ORBIT mutants were confirmed by amplicon sequencing of the cyp123 gene. The ORBIT mutants are named Δ*cyp123::hyg-1*, Δ*cyp123::hyg-2* and Δ*cyp51::hyg-1* here, and Δ*cyp123-1*, Δ*cyp123-2* and Δ*cyp51-1* in the main text and Fig. 4.

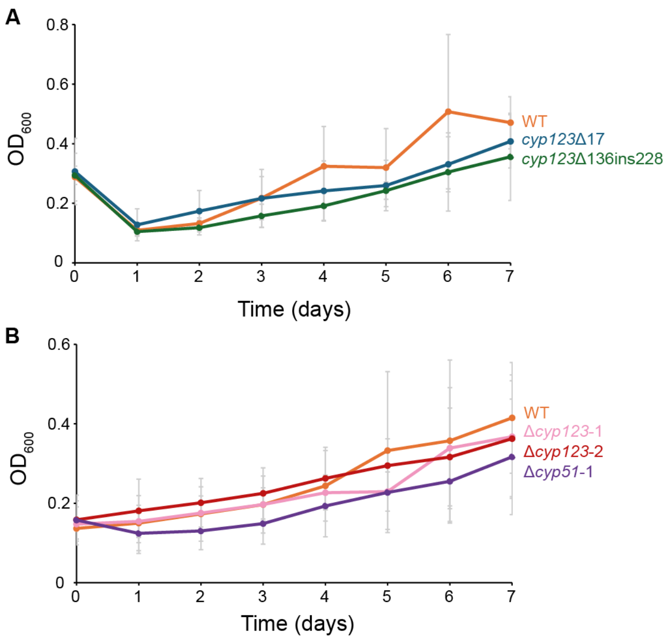


**Figure S12: Growth of *M. smegmatis cyp123* and *cyp51* mutants on lanosterol as a sole carbon source.** Growth of wild-type *M. smegmatis*, *cyp123* CRISPR (A) and *cyp123* and *cyp51* ORBIT (B) mutants in minimal medium supplemented with lanosterol (1 mM) as the sole carbon source, over seven days. $OD_{600}$ was measured every 24 h; data represent the mean ± SD of six technical replicates.

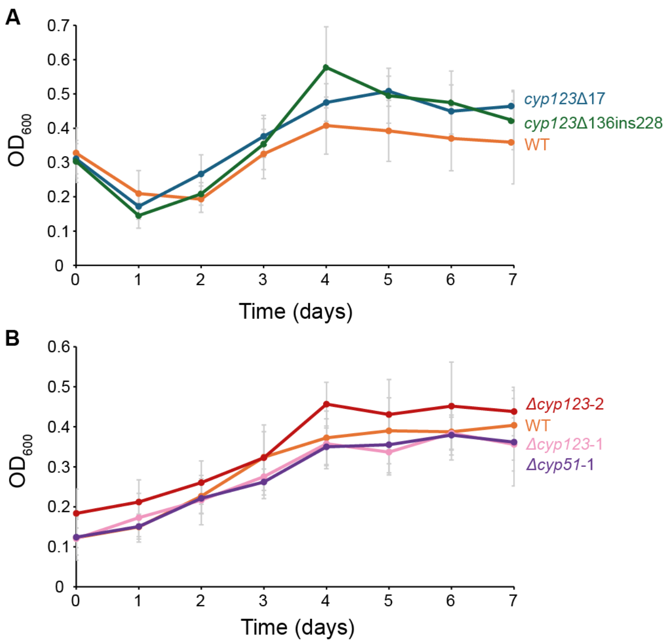


**Figure S13: Growth of *M. smegmatis cyp123* and *cyp51* mutants on cholesterol as a sole carbon source.** Growth of wild-type *M. smegmatis*, *cyp123* CRISPR (A) and *cyp123* and *cyp51* ORBIT (B) mutants in minimal medium supplemented with cholesterol (1 mM) as the sole carbon source. $OD_{600}$ was measured every 24 h; data represent the mean ± SD of six technical replicates.

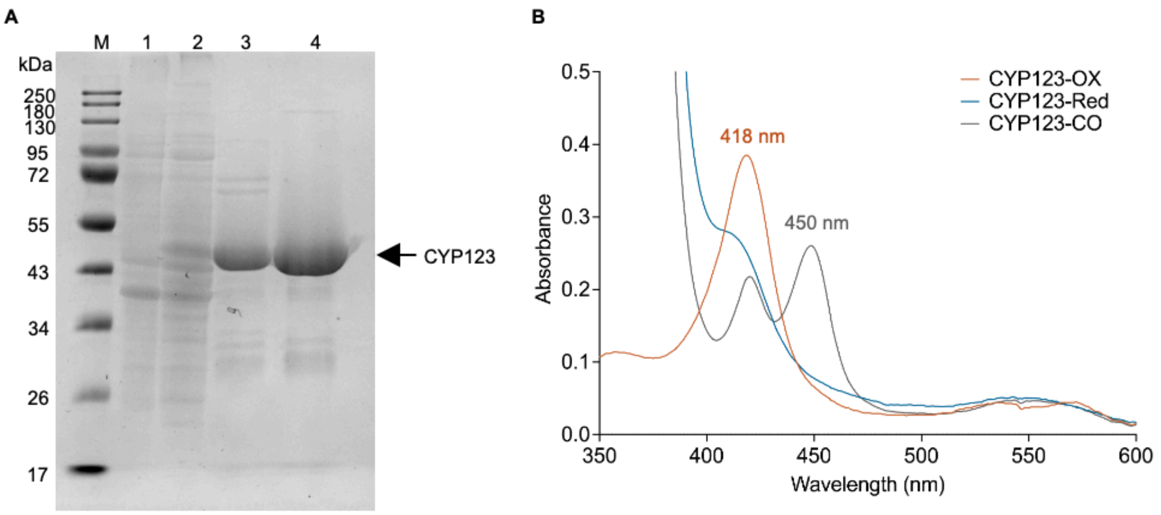


**Figure S14: Purification and spectroscopic characterisation of *M. smegmatis* CYP123. (A)** SDS-PAGE analysis of *M. smegmatis* CYP123 overexpression and purification. The gel lanes are: protein ladder (M), pre-induction cell pellets (lane 1), supernatant of protein-overexpressing cell pellets (lane 2), sample after Ni-NTA affinity chromatography purification (lane 3) and sample after size exclusion chromatography purification (lane 4), with the target CYP123 protein exhibiting a size of 45.9 kDa. **(B)** UV-visible spectra for different states of *M. smegmatis* CYP123. The ferric resting state (orange), the reduced enzyme (blue) and the reduced CO-bound (grey) spectra are shown. The enzyme has a split A450/A418 peak in the CO-bound state.

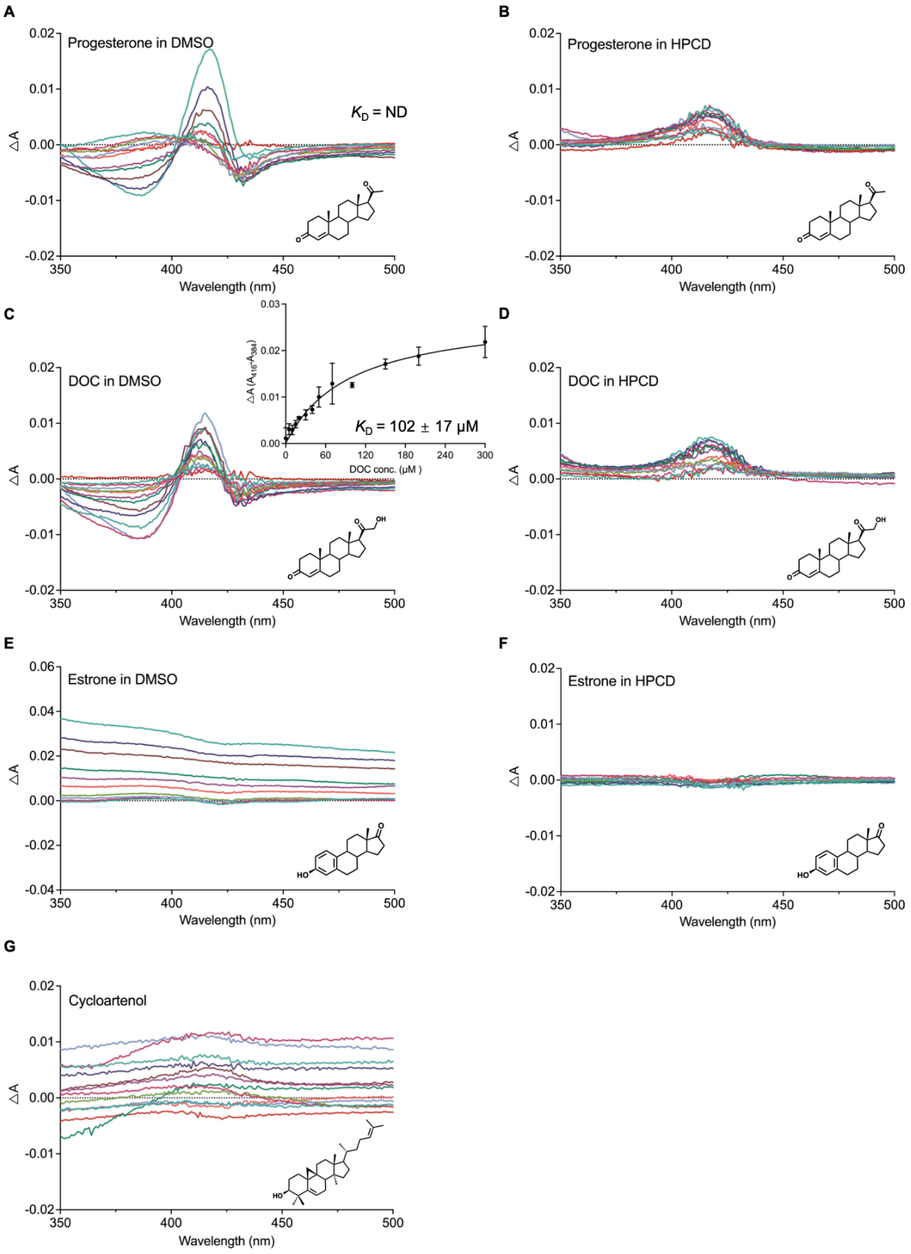


**Figure S15: Other steroids binding to CYP123 and their effect on substrate delivery method.** Difference spectra obtained upon titration of CYP123 with selected steroidal compounds using either DMSO or hydroxypropyl-β-cyclodextrin (HPCD) for substrate delivery. **(A, B)** Progesterone dissolved in DMSO and HPCD, respectively. Progesterone in DMSO showed a transition from a type I to a reverse type I spectral response with increasing ligand concentration, precluding reliable determination of a binding constant, whereas no characteristic spectral response was observed with HPCD delivery. **(C, D)** 11-Deoxycorticosterone (DOC) delivered in DMSO and HPCD, respectively. DOC in DMSO produced a reverse type I spectral response with a $K_D$ of 102 ± 17 µM, whereas no characteristic spectral response was observed with HPCD delivery. **(E, F)** Estrone delivered in DMSO and HPCD, respectively. Estrone in HPCD produced no detectable spectral response, whereas delivery in DMSO caused a baseline perturbation rather than a binding spectrum. **(G)** Cycloartenol delivered in HPCD. Cycloartenol in HPCD caused a baseline perturbation rather than a binding spectrum. Insets in **(C)** show binding curves used to determine $K_D$ values by plotting the peak-to-trough absorbance differences against substrate concentration.

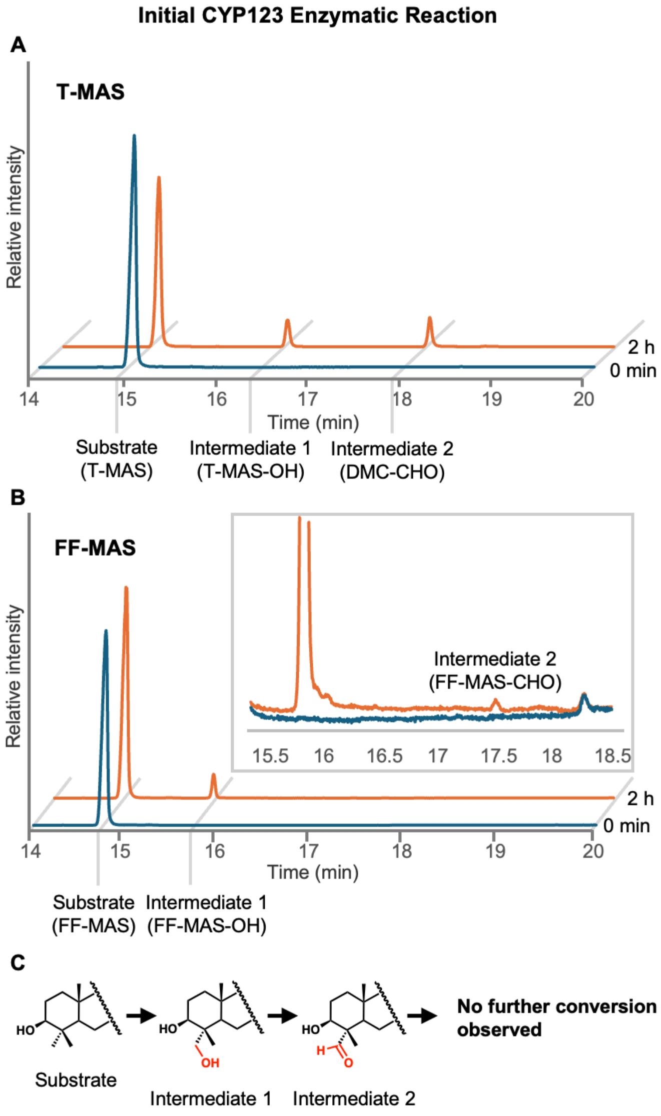


**Figure S16: CYP123 enzymatic reactions with 4,4-dimethylated sterol substrates using a single NADPH addition.** GC-MS total ion chromatograms showing oxidation of T-MAS **(A)** and FF-MAS **(B)** by CYP123. Reactions contained 5 µM CYP123, 0.25 mM substrate, 9 µM SoFdx, 1 µM SoFNR, and 10 mM NADPH. Substrate controls (blue) are compared with reacted samples (orange). Both substrates yielded hydroxylated (-OH) and aldehyde (-CHO) intermediates but no carboxylic acid product under these conditions, indicating that a single NADPH addition is insufficient for complete oxidation. **(C)** Proposed stepwise oxidation of the C4-methyl group of 4,4-dimethylated sterols by CYP123. GC-MS analyses were performed on an Agilent 7890B GC coupled to an Agilent 7200 Accurate-Mass Q-TOF, equipped with an Agilent VF-5ms column.

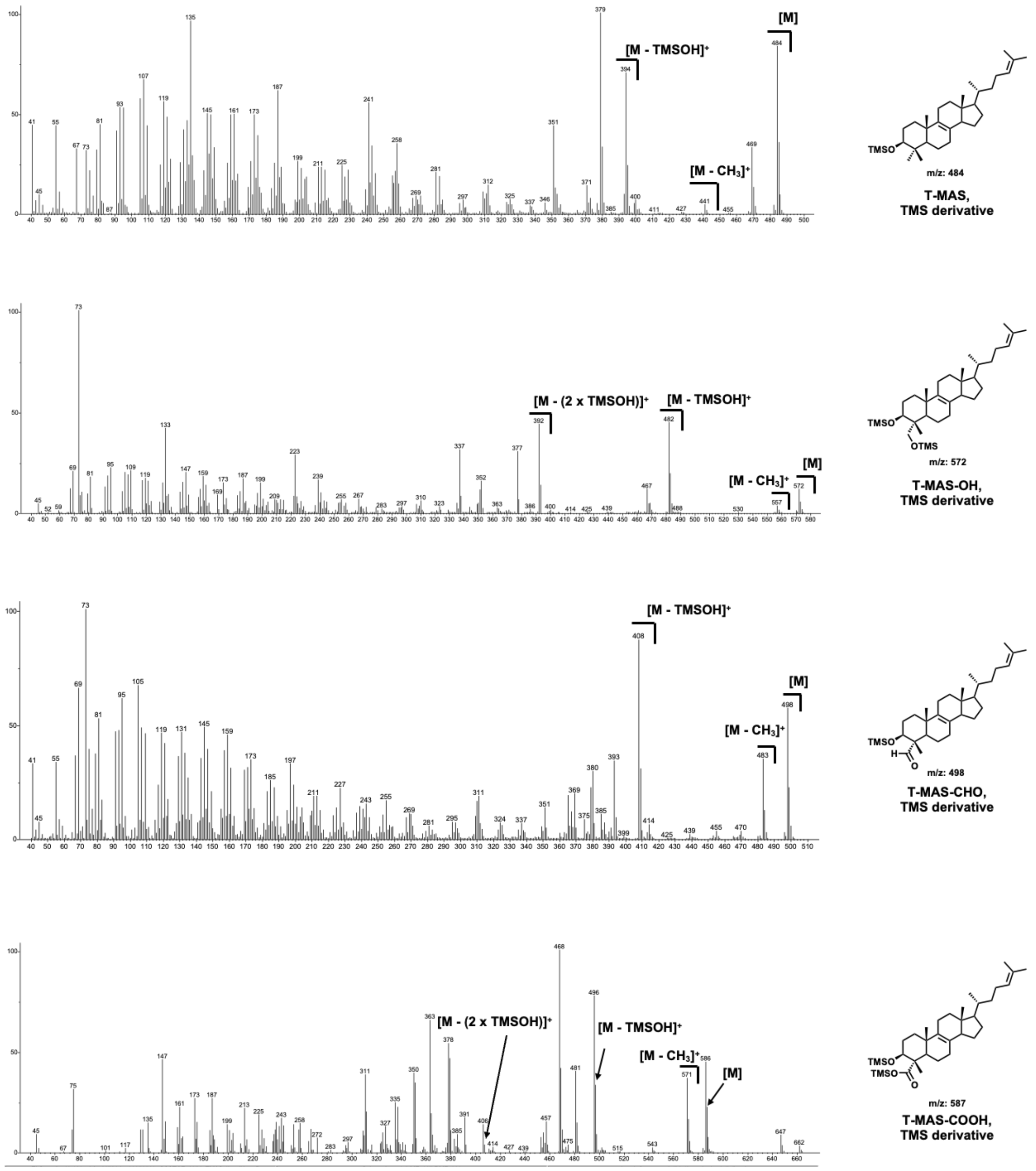


**Figure S17: Mass spectra of BSTFA-derivatised T-MAS and its CYP123-generated oxidation products, corresponding to the chromatographic peaks in Fig. 6A.** Spectra are shown for T-MAS (substrate), T-MAS-OH (C4-hydroxylated intermediate, +88 Da), T-MAS-CHO (C4-aldehyde intermediate, +14 Da), and T-MAS-COOH (C4-carboxylic acid product, +102 Da). Mass shifts are relative to the underivatised substrate and include TMS groups introduced during derivatisation.

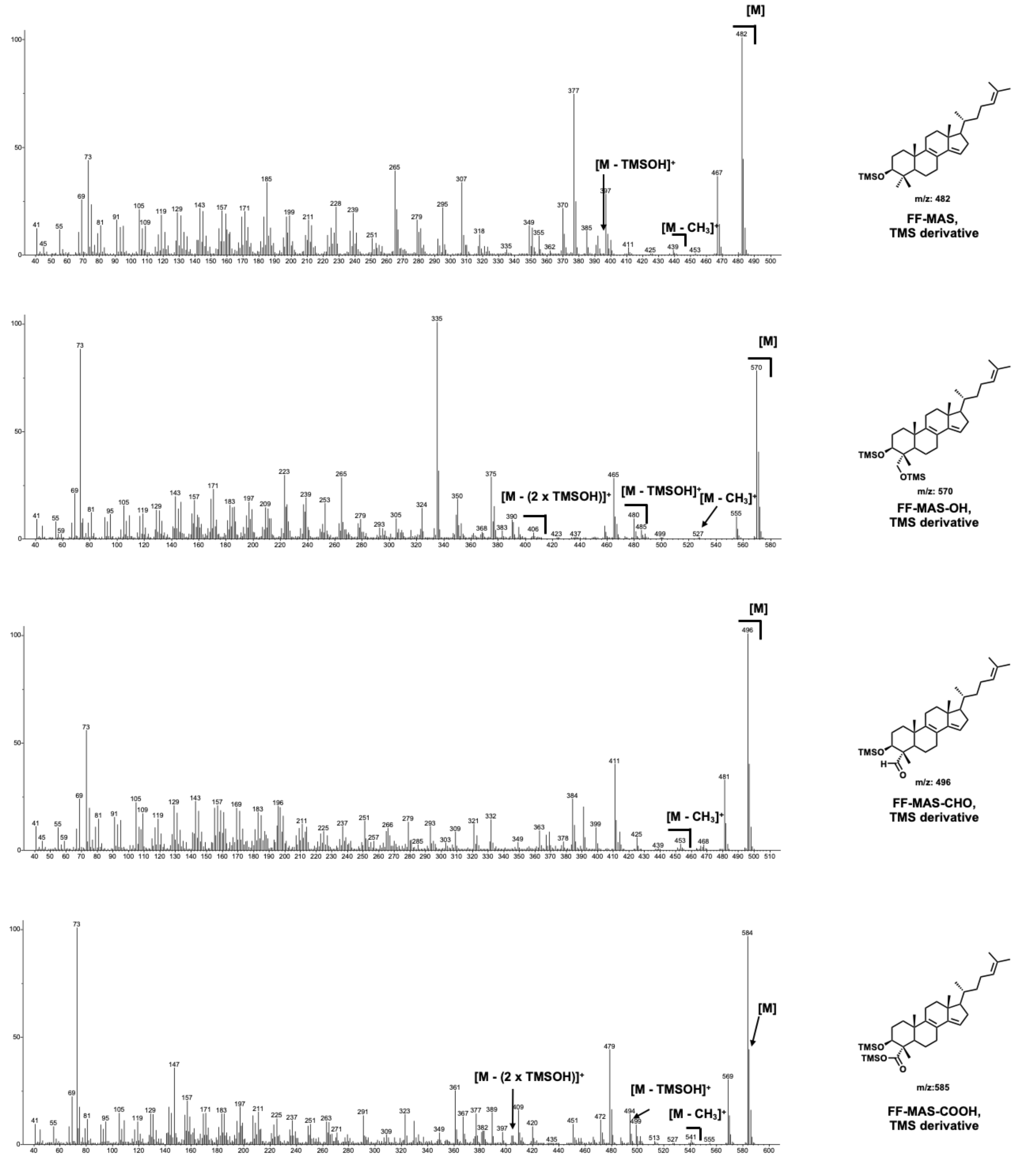


**Figure S18: Mass spectra of BSTFA-derivatised FF-MAS and its CYP123-generated oxidation products, corresponding to the chromatographic peaks in Fig. 6B.** Spectra are shown for FF-MAS (substrate), FF-MAS-OH (C4-hydroxylated intermediate, +88 Da), FF-MAS-CHO (C4-aldehyde intermediate, +14 Da), and FF-MAS-COOH (C4-carboxylic acid product, +102 Da). Mass shifts are as described for Fig. S17.

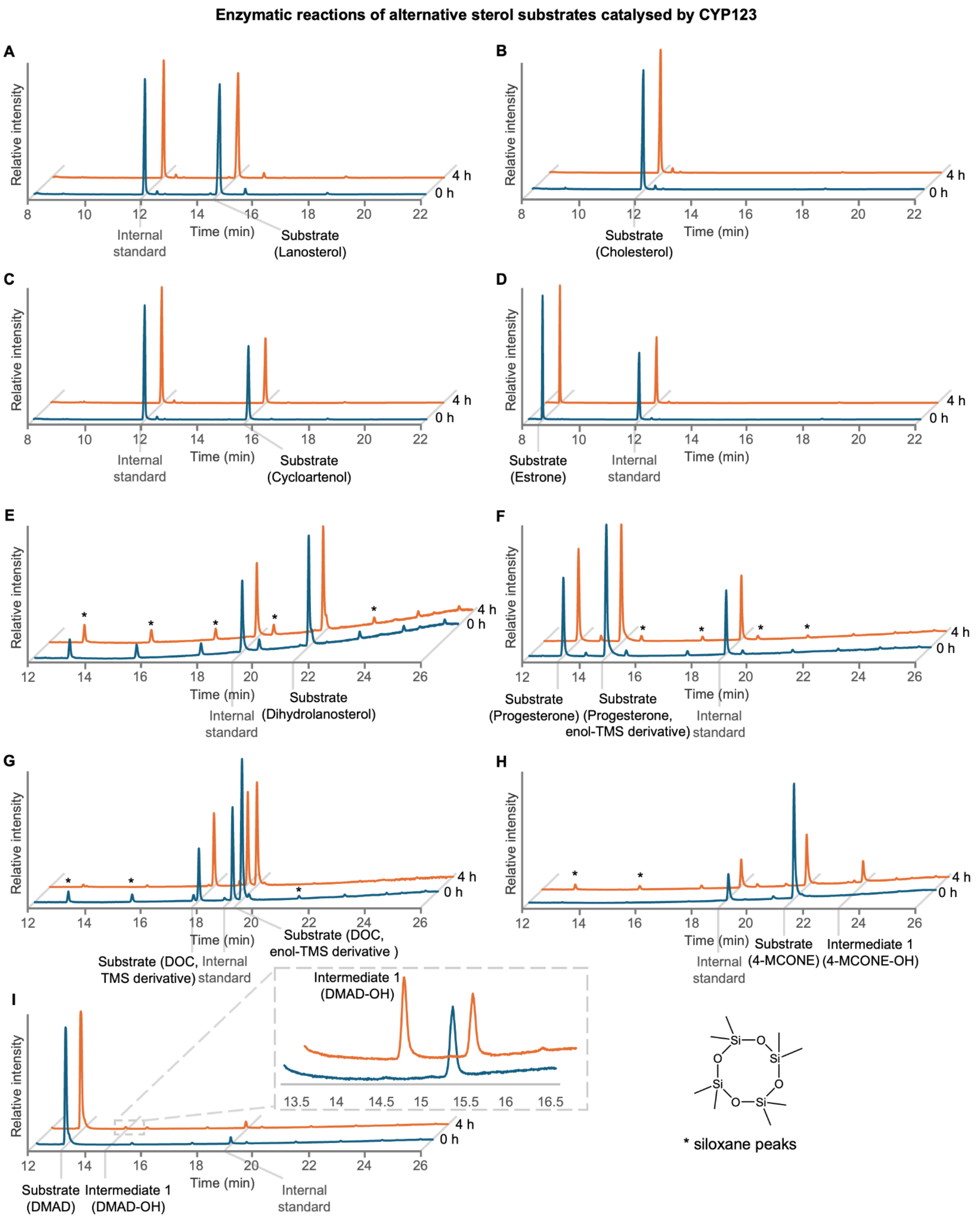


**Figure S19: Substrate specificity of CYP123: GC-MS analysis of sterols exhibiting limited detectable turnover by CYP123.** Total ion chromatograms of CYP123 reactions containing **(A)** lanosterol, **(B)** cholesterol, **(C)** cycloartenol, **(D)** estrone, **(E)** dihydrolanosterol, **(F)** progesterone, **(G)** 11-deoxycorticosterone (DOC), **(H)** 4-methylcholest-4-en-3-one (4-MCONE), and **(I)** 4,4-dimethyl-5α-androstane-3β,17β-diol (DMAD). Reactions contained 0.5 mM substrate, 5 μM CYP123, 9 μM SoFdx, 1 μM SoFNR, and 10 mM NADPH. All enzyme components and NADPH were resupplemented every 30 min to sustain turnover. Chromatograms at 0 h (blue) represent no-reaction controls; chromatograms at 4 h (orange) show reactions after incubation. No product peaks were detected for lanosterol, cholesterol, cycloartenol, estrone, dihydrolanosterol, progesterone or DOC (A–G); 4-MCONE and DMAD (H, I) gave only the corresponding hydroxylated intermediates. None of these sterols is a preferred substrate of CYP123 under the conditions tested. GC-MS analyses were performed on an Agilent 7890B GC coupled to an Agilent 7200 Accurate-Mass Q-TOF, equipped with an Agilent VF-5ms column.

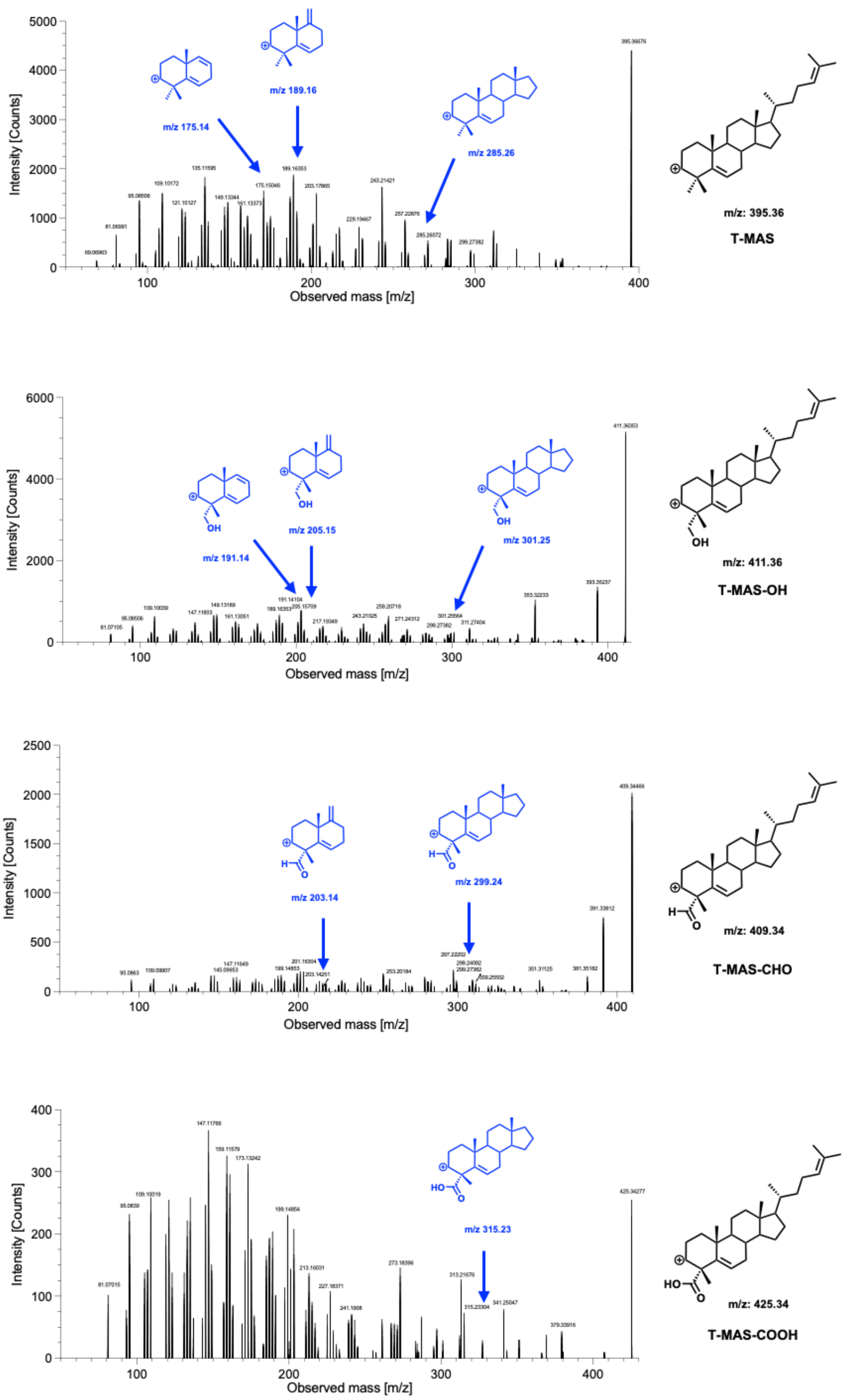


**Figure S20: Electrospray ionisation tandem mass spectrometry (ESI-MS/MS) fragmentation of T-MAS and CYP123-generated oxidation products.** ESI-MS/MS spectra of $[M{-}H_2O{+}H]^+$ ions from T-MAS and its oxidised products from the CYP123 reaction. Key diagnostic fragments are highlighted in blue.

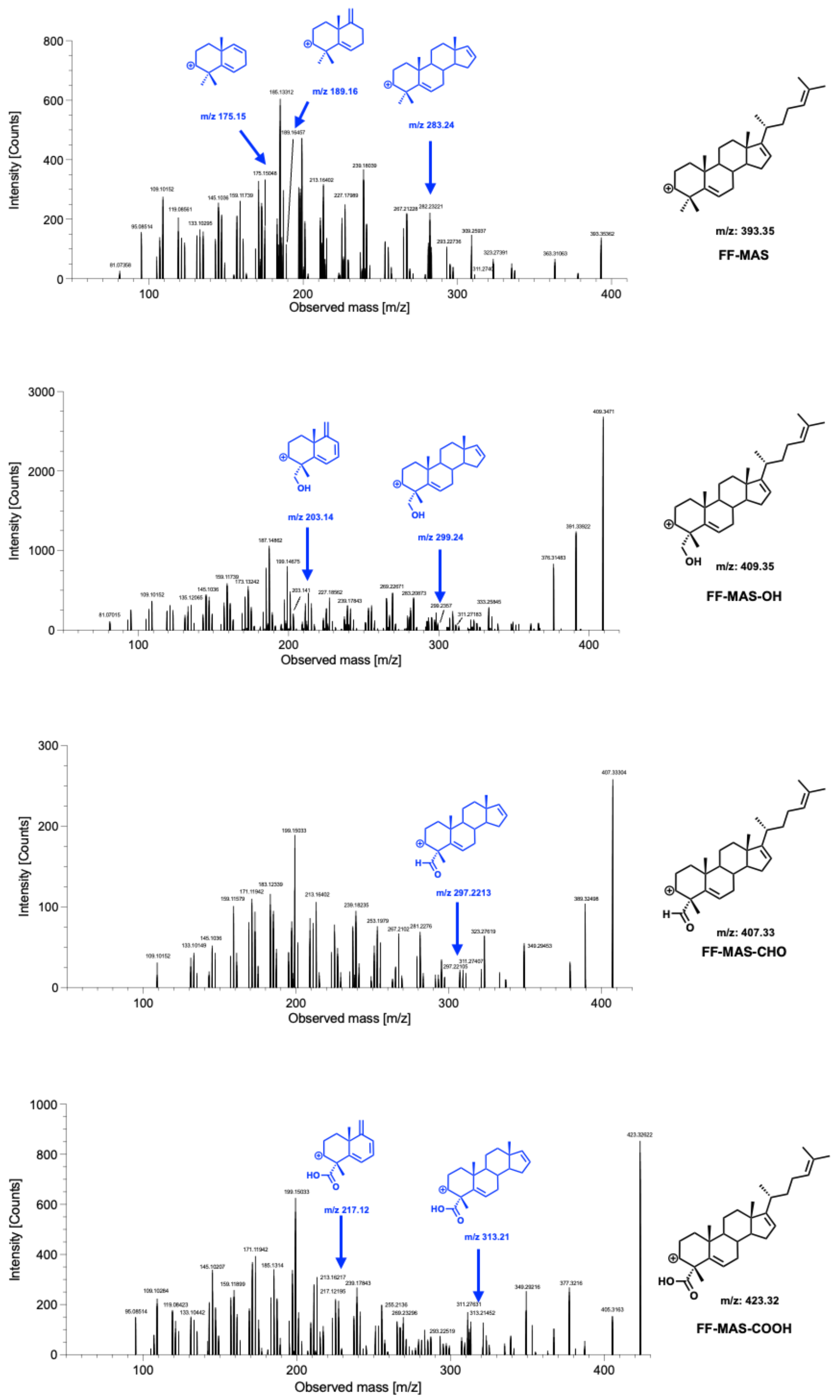


**Figure S21: LC-ESI-MS/MS fragmentation of FF-MAS and CYP123-generated oxidation products.** ESI-MS/MS spectra of $[M{-}H_2O{+}H]^+$ ions from FF-MAS and its oxidised products from the CYP123 reaction. Key diagnostic fragments are highlighted in blue.

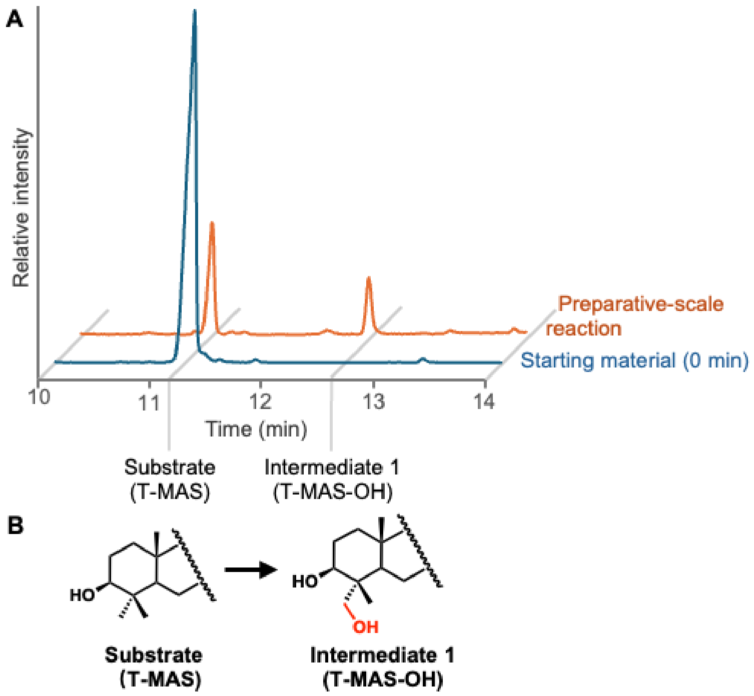


**Figure S22: Preparative-scale CYP123 enzymatic reaction with T-MAS as substrate. (A)** GC-MS chromatograms of the starting substrate T-MAS (blue) and the products obtained from a 10 mL CYP123 reaction (orange). The reaction contained 1 mg T-MAS, 5 mM NADPH, 5 μM CYP123, 9 μM SoFdx, 1 μM SoFNR, and 0.1 mg $mL^{-1}$ catalase, incubated at 30 °C for 10 h. A new peak corresponding to the alcohol intermediate was observed and extracted for NMR characterisation. **(B)** Proposed conversion of T-MAS to the monohydroxylated intermediate (T-MAS-OH) by CYP123. GC-MS analyses were performed on an Agilent 7890B GC coupled to an Agilent 5975 series MSD, equipped with an Agilent VF-5ht column.

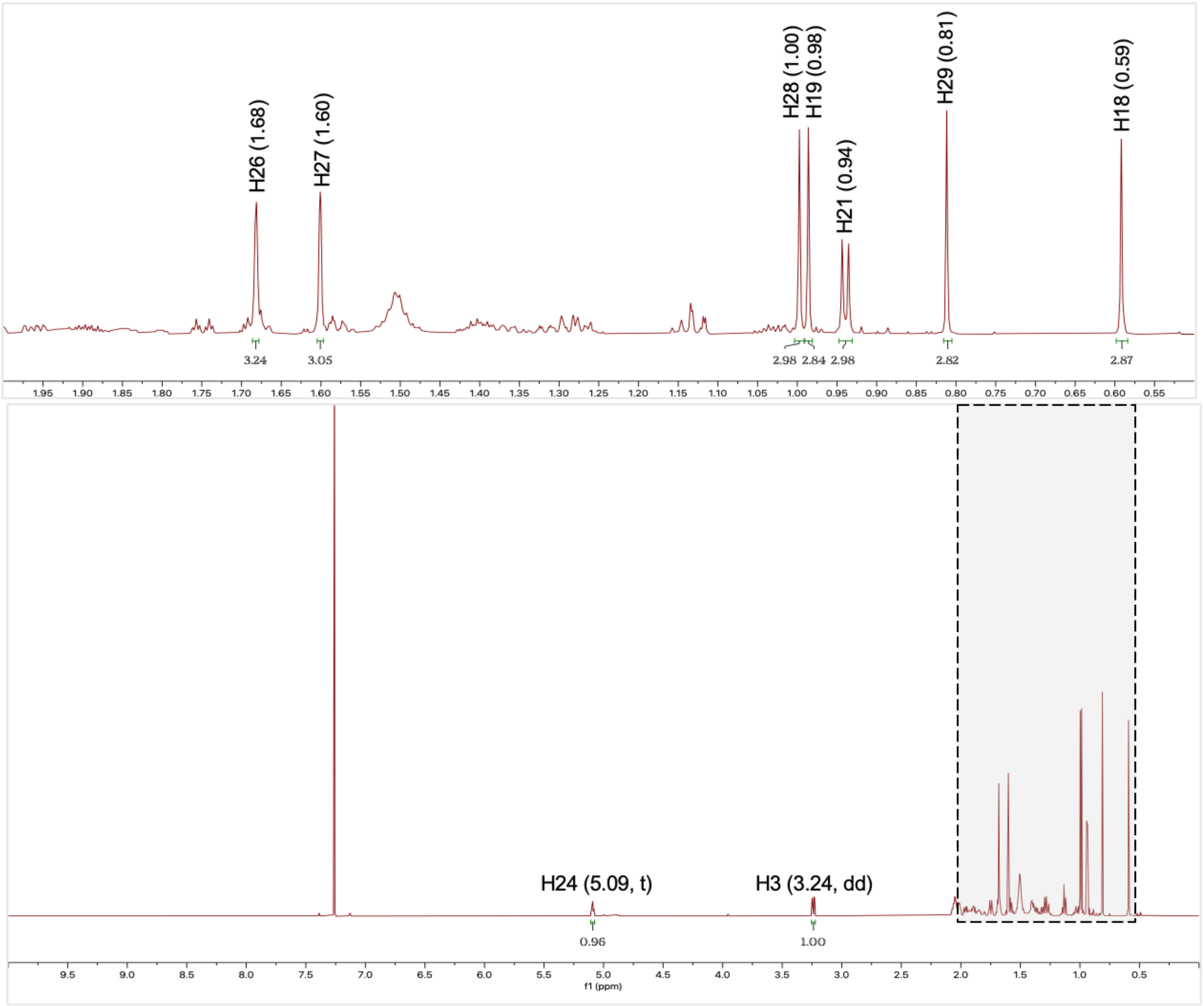


**Figure S23: $^{1}$H NMR spectrum of T-MAS.** $CDCl_3$, 800 MHz.

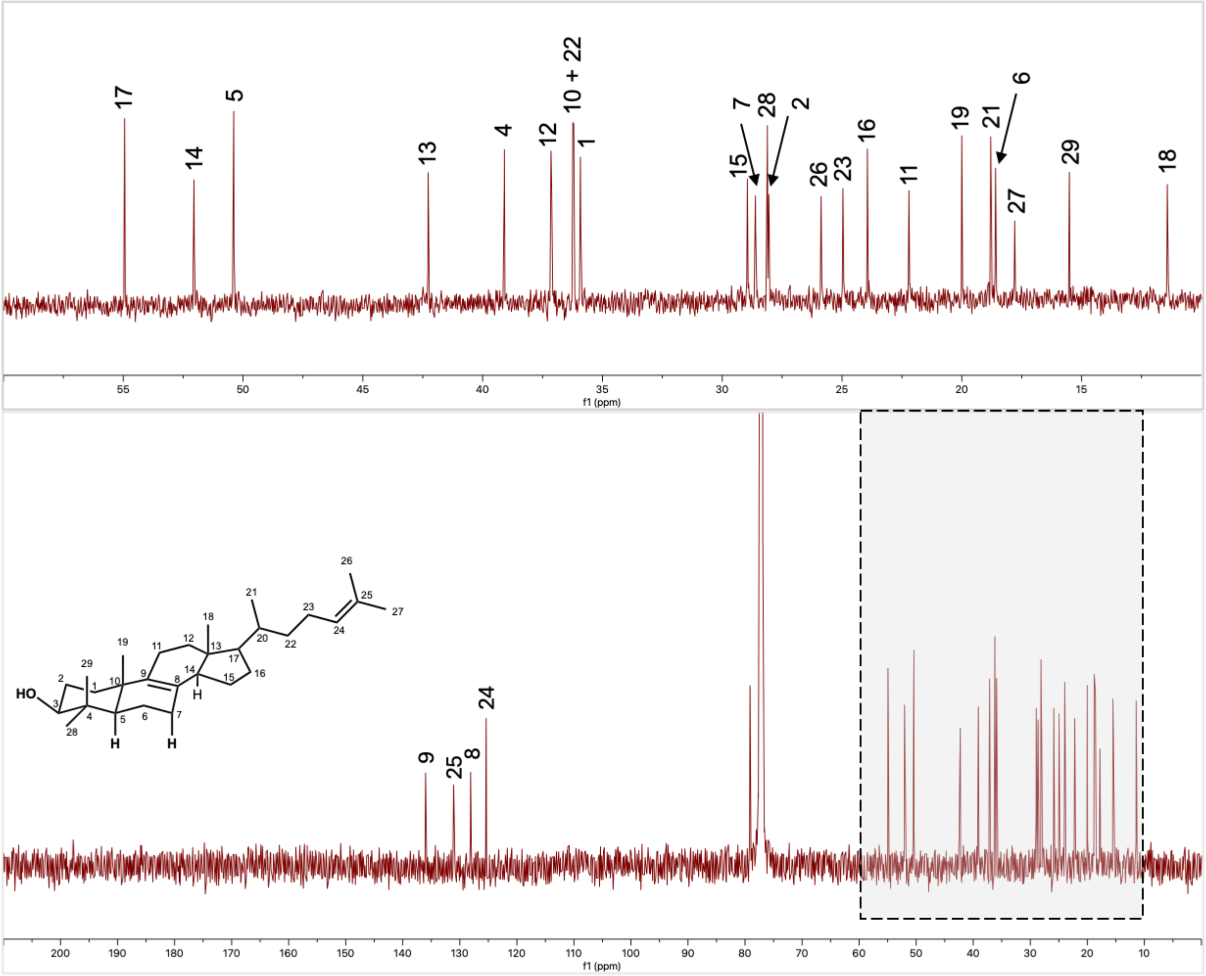


**Figure S24: $^{13}$C NMR spectrum of T-MAS.** $CDCl_3$, 800 MHz.

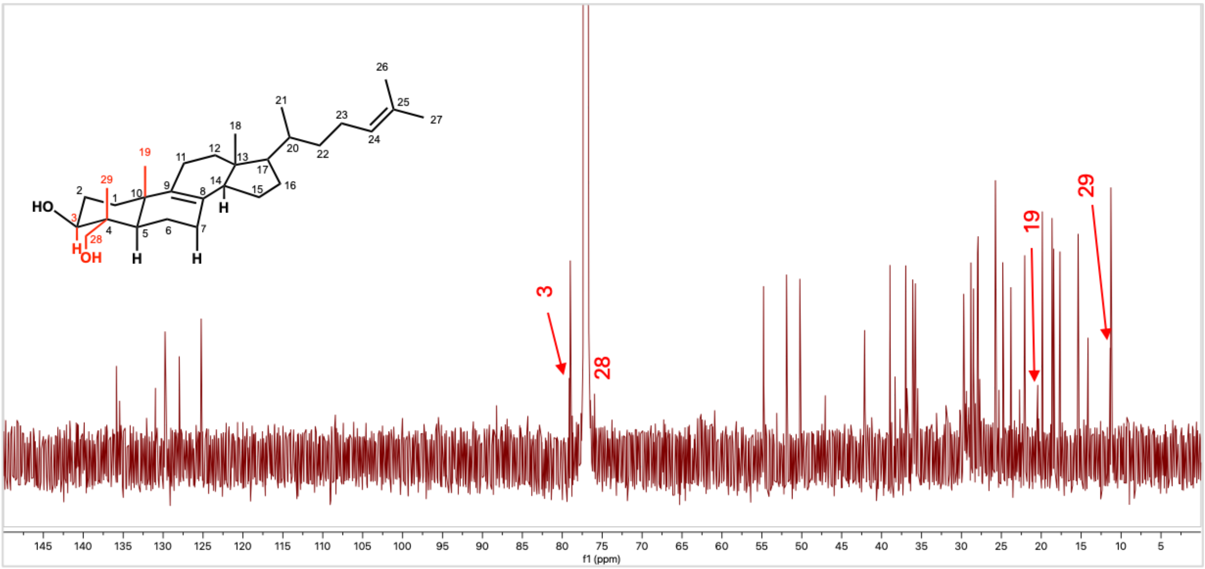


**Figure S25: $^{13}$C NMR spectrum of the T-MAS and T-MAS-OH mixture.** $CDCl_3$, 800 MHz. T-MAS-OH carbons are highlighted in red.

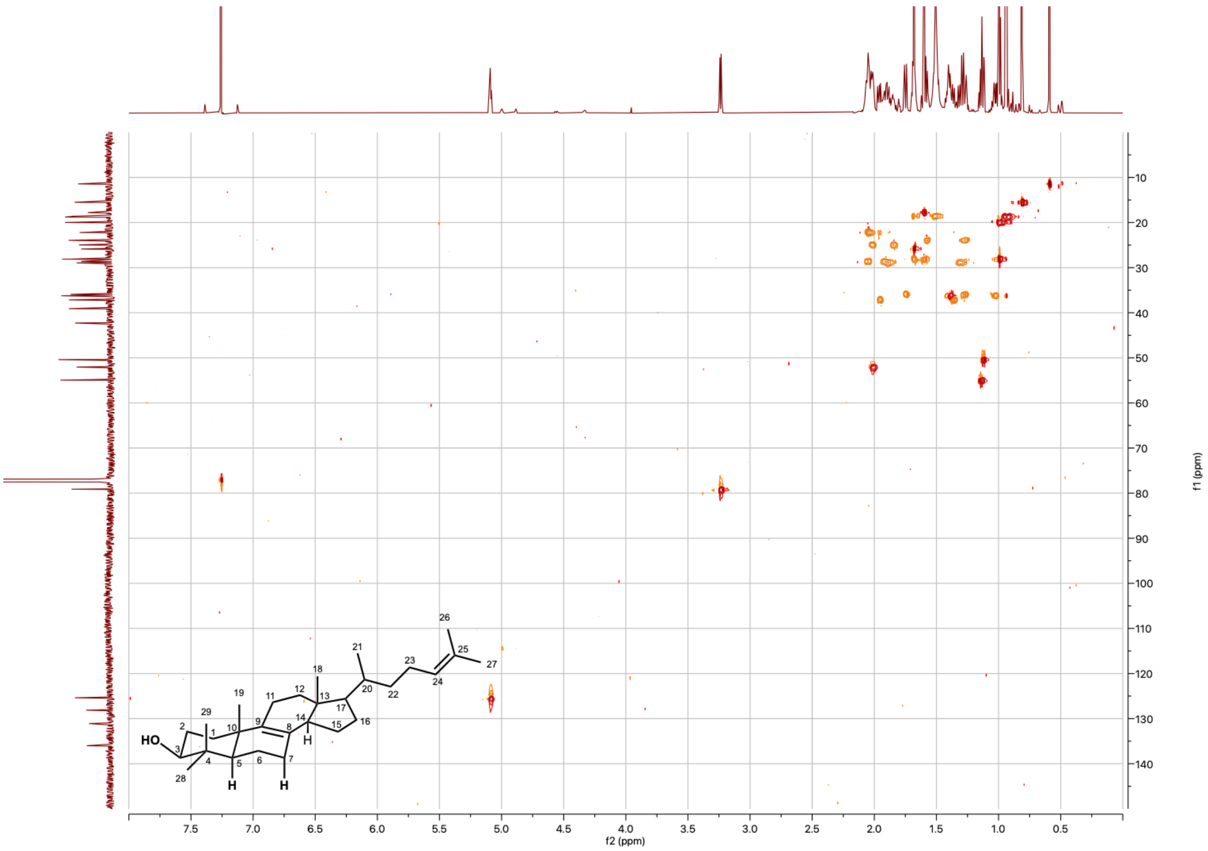

**Figure S26: Heteronuclear single quantum coherence (HSQC) NMR spectrum of T-MAS.** $CDCl_3$, 800 MHz.

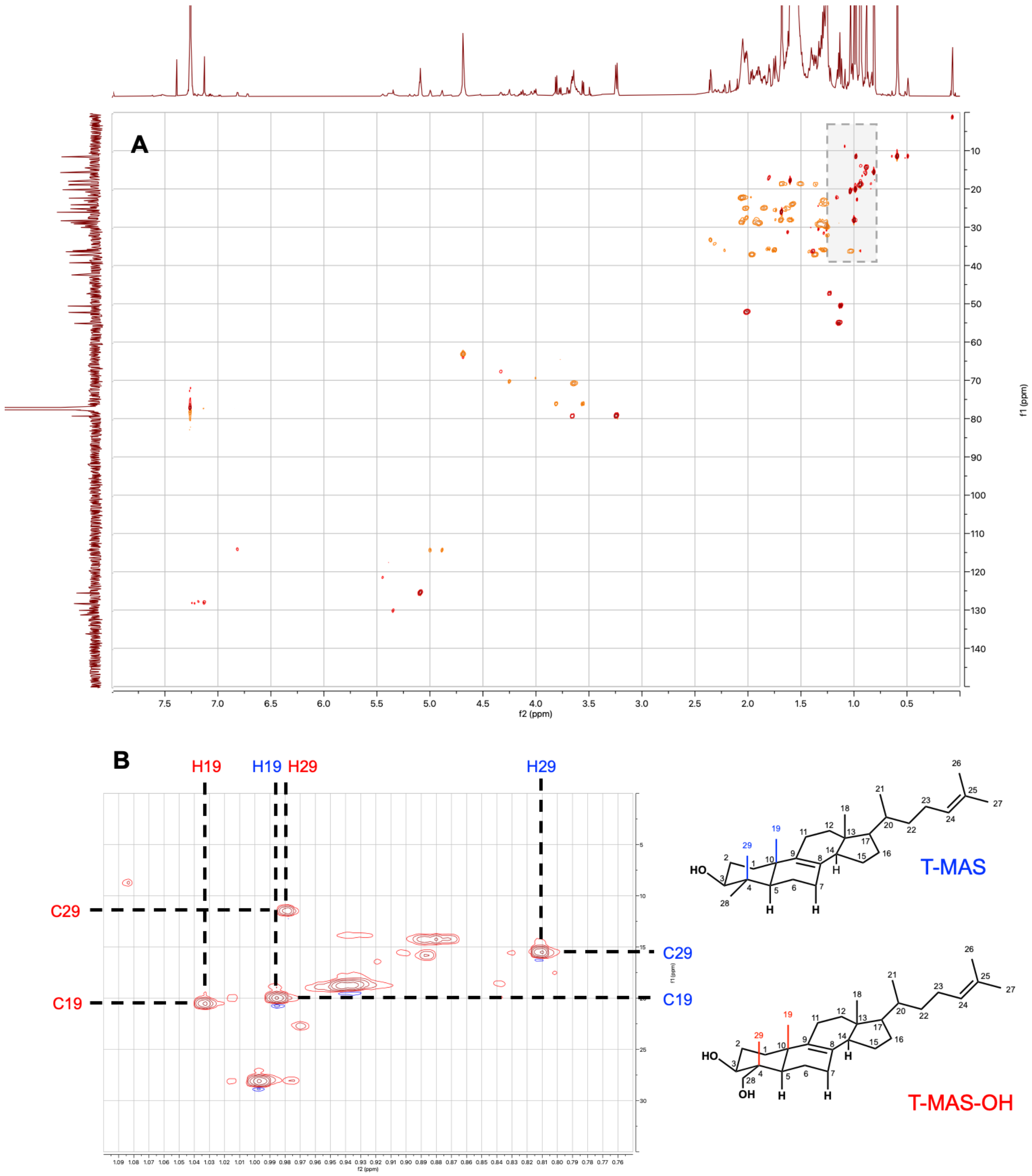


**Figure S27: HSQC NMR of the T-MAS and T-MAS-OH mixture.** $CDCl_3$, 800 MHz. **(A)** Full HSQC spectrum. **(B)** Expanded view showing HSQC correlations for T-MAS (blue) and T-MAS-OH (red) between methyl group carbons C-19 and C-29 and their corresponding protons H-19 and H-29.

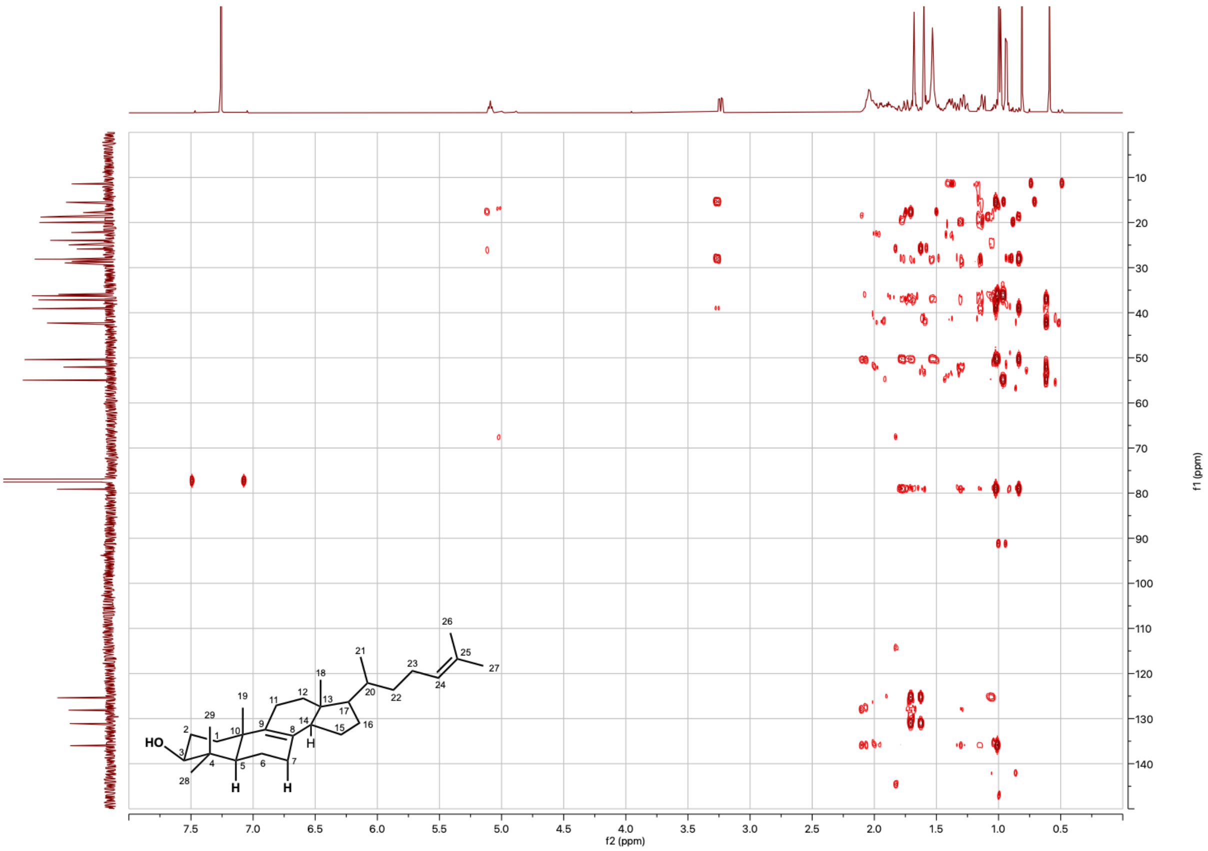


**Figure S28: Heteronuclear multiple bond correlation (HMBC) NMR spectrum of T-MAS.** $CDCl_3$, 500 MHz.

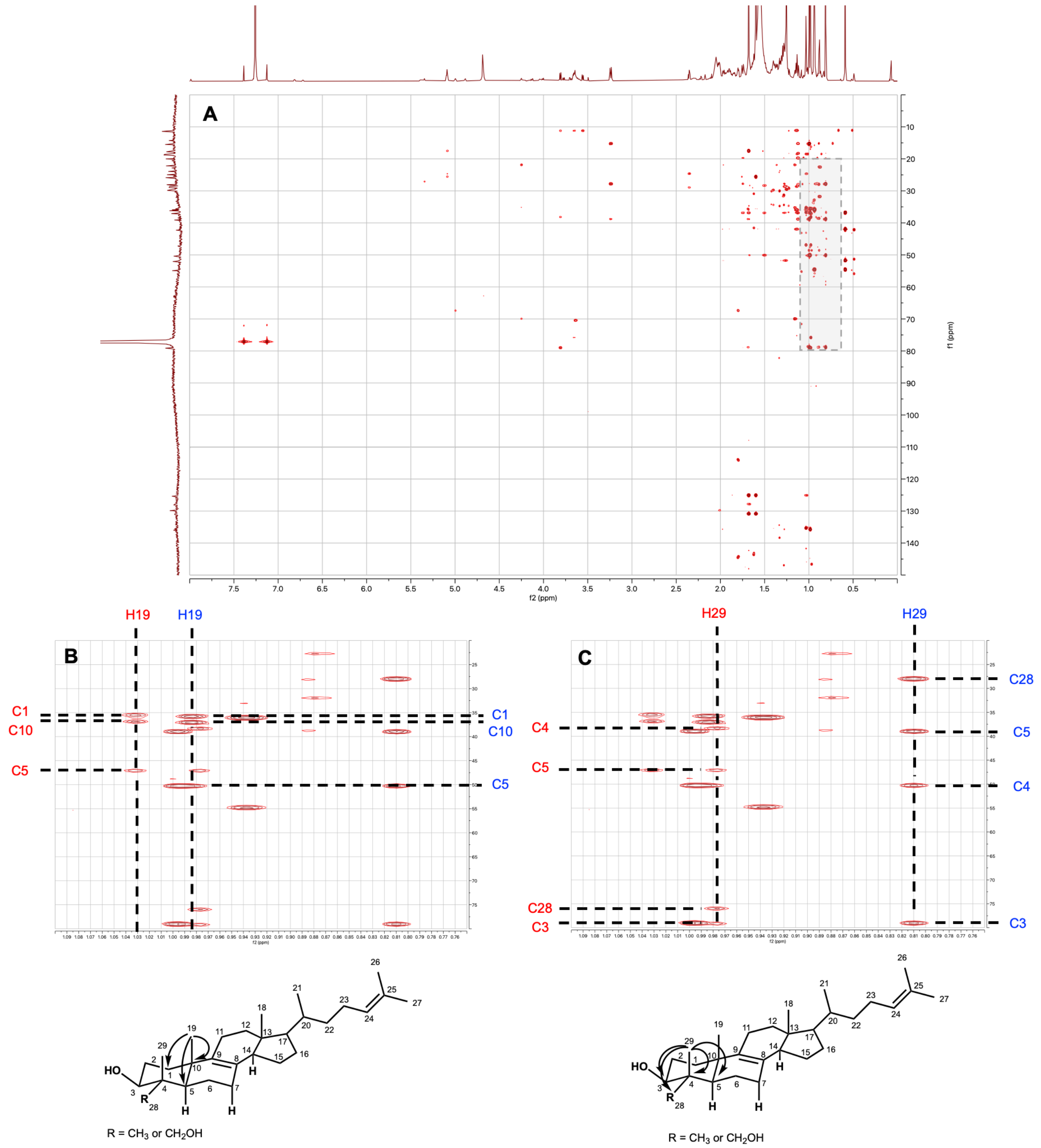


**Figure S29: HMBC NMR of the T-MAS and T-MAS-OH mixture.** $CDCl_3$, 800 MHz. **(A)** Full HMBC spectrum. **(B)** HMBC correlations for T-MAS (blue) and T-MAS-OH (red) showing connectivities of carbons C-1, C-10, and C-5 with methyl group protons H-19. **(C)** HMBC correlations for T-MAS (blue) and T-MAS-OH (red) showing connectivities of carbons C-4, C-5, C-28, and C-3 with methyl group protons H-29.

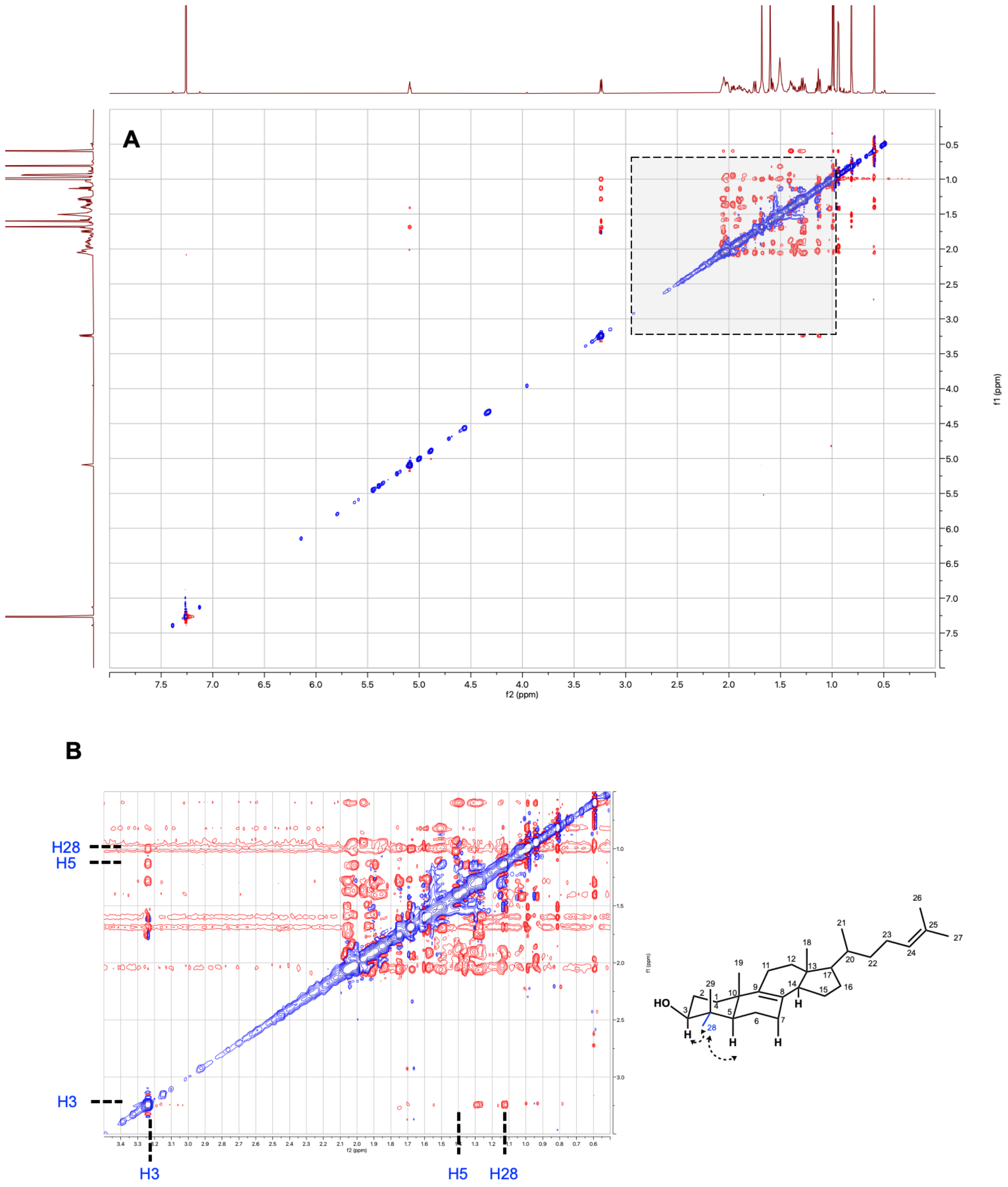


**Figure S30: Rotating frame Overhauser effect spectroscopy (ROESY) NMR of T-MAS.** $CDCl_3$, 800 MHz. **(A)** Full ROESY spectrum. **(B)** Nuclear Overhauser effect (NOE) correlations between H-28 and nearby protons H-3 and H-5, confirming the equatorial (coplanar) orientation of the C4α-methyl group.

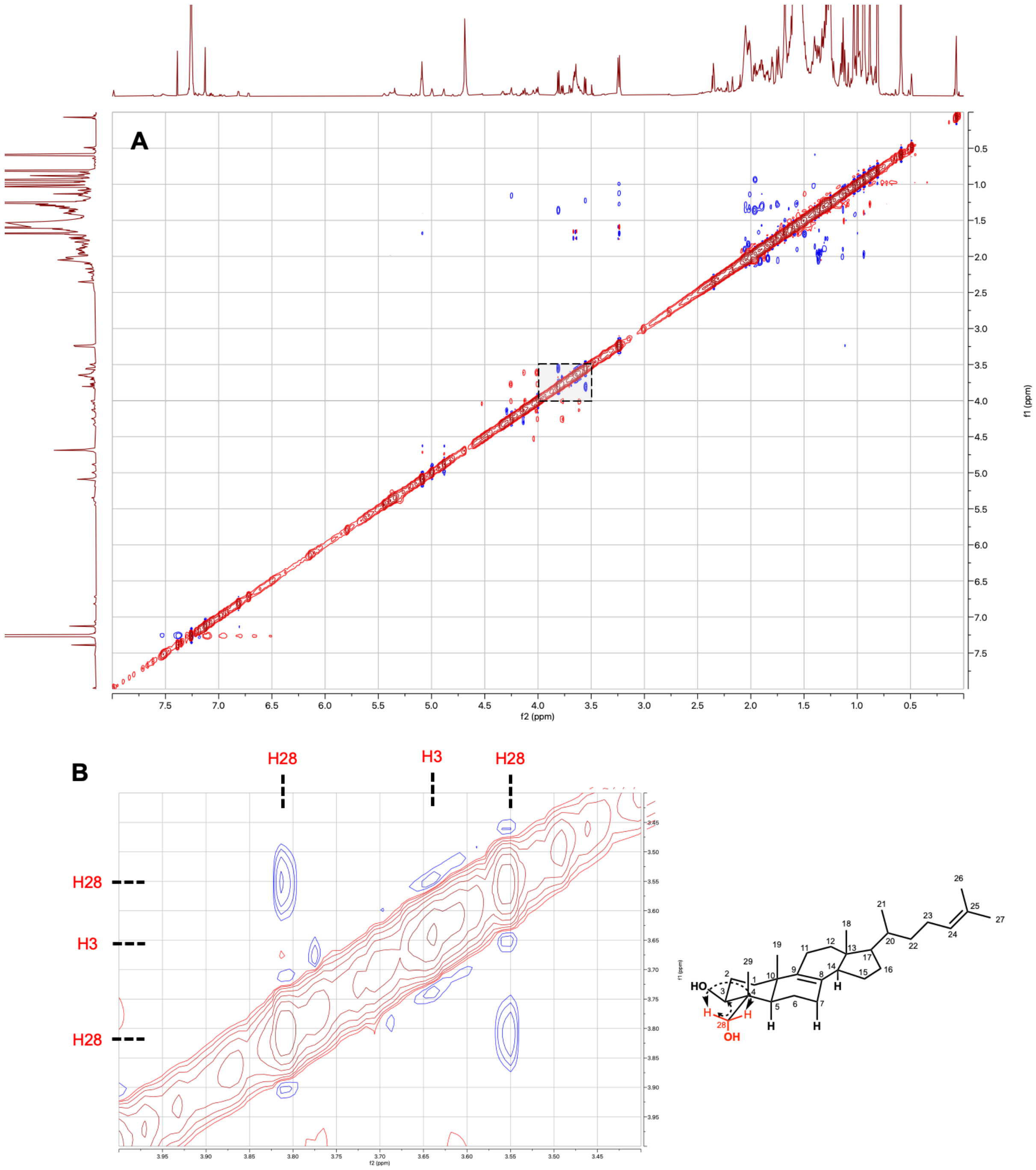


**Figure S31: ROESY NMR of the T-MAS and T-MAS-OH mixture.** $CDCl_3$, 800 MHz. **(A)** Full ROESY spectrum. **(B)** NOE correlations between the two H-28 $CH_2$ protons and nearby proton H-3, confirming oxidation at the equatorial C4α-methyl group.

**Table S3: $^{1}H$ and $^{13}C$ NMR chemical shift assignments for T-MAS.** Spectra recorded in $CDCl_3$ at 800 MHz. Chemical shifts (δ) in ppm; coupling constants (J) in Hz.

| | $^{1}H$ | $^{13}C$ |
|---|---|---|
| **1** | 1.74 (m) 1.26 (m) | 35.92 |
| **2** | 1.59 (m), 1.67 (m) | 28.06 |
| **3** | 3.24 (dd) 1H, 11.8, 4.4 Hz | 79.13 |
| **4** | - | 39.09 |
| **5** | 1.13 (d) 2.1 Hz | 50.39 |
| **6** | 1.68 (m), 1.50 (m) | 18.60 |
| **7** | 2.04 (m), 1.89 (m) | 28.61 |
| **8** | - | 128.11 |
| **9** | - | 136.01 |
| **10** | - | 36.04 |
| **11** | 2.04 (m) | 22.21 |
| **12** | 1.95 (m), 1.36 (m) | 37.15 |
| **13** | - | 42.27 |
| **14** | 2.03 (m) | 52.05 |
| **15** | 1.41 (m) | 28.95 |
| **16** | 1.26 (m), 1.57 (m) | 23.94 |
| **17** | 1.13 (m) | 54.94 |
| **18** | 0.59 (s) 3H | 11.26 |
| **19** | 0.98 (s) 3H | 20.00 |
| **20** | 1.37 (m) | 36.20 |
| **21** | 0.94 (d) 3H 6.5 Hz | 18.79 |
| **22** | 1.03 (m) | 37.21 |
| **23** | 1.8, 2.0 (m) | 24.96 |
| **24** | 5.09 (t) 1H 7.1 Hz | 125.38 |
| **25** | - | 131.09 |
| **26** | 1.68 (s) 3H | 25.87 |
| **27** | 1.60 (s) 3H | 17.79 |
| **28** | 1.00 (s) 3H | 28.11 |
| **29** | 0.81 (s) 3H | 15.51 |

**Table S4: $^{1}H$ and $^{13}C$ NMR chemical shift assignments for T-MAS-OH.** Spectra recorded in $CDCl_3$ at 800 MHz. Chemical shifts (δ) in ppm; coupling constants (J) in Hz.

| | $^{1}H$ | $^{13}C$ |
|---|---|---|
| **1** | 1.74 (m) 1.26 (m) | 35.37 |
| **2** | 1.64 (m), 1.74 (m) | 25.44 |
| **3** | 3.65 (dd) 1H 12.2 Hz, 4.2 Hz | 79.11 |
| **4** | - | 38.25 |
| **5** | 1.23 (d) | 47.01 |
| **6** | not assignable | |
| **7** | not assignable | |
| **8** | - | 128.11 |
| **9** | - | 135.53 |
| **10** | - | 35.89 |
| **11** | 2.04 (m) | 22.21 |
| **12** | 1.95 (m), 1.36 (m) | 37.15 |
| **13** | - | 42.27 |
| **14** | 2.03 (m) | 52.05 |
| **15** | 1.41 (m) | 28.95 |
| **16** | 1.26 (m), 1.57 (m) | 23.94 |
| **17** | 1.13 (m) | 54.94 |
| **18** | 0.59 (s) 3H | 11.49 |
| **19** | 1.03 (s) 3H | 20.37 |
| **20** | 1.37 (m) | 36.20 |
| **21** | 0.94 (d) 3H 6.5 Hz | 18.79 |
| **22** | 1.03 (m) | 37.21 |
| **23** | 1.8, 2.0 (m) | 24.96 |
| **24** | 5.09 (t) 1H 7.1 Hz | 125.38 |
| **25** | - | 131.09 |
| **26** | 1.68 (s) 3H | 25.87 |
| **27** | 1.60 (s) 3H | 17.79 |
| **28** | 3.80, 3.55 (2 x d) 10.1 Hz | 75.88 |
| **29** | 0.98 (s) 3H | 11.28 |

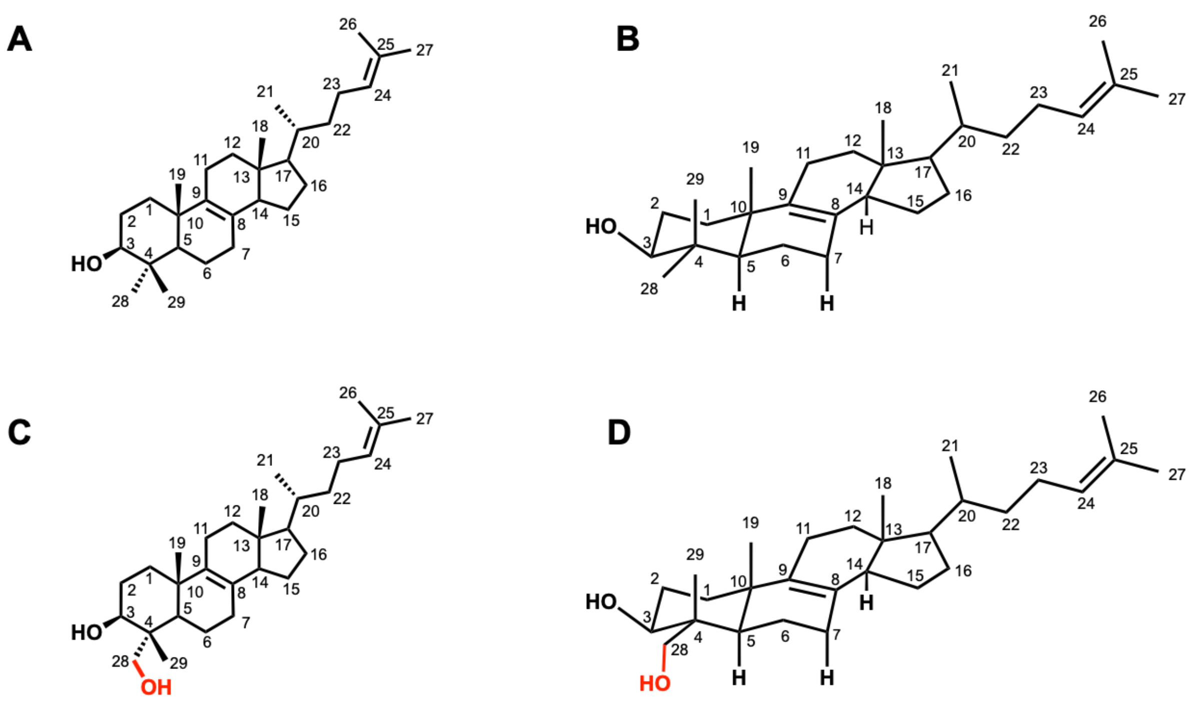


**Figure S32: Structures and conformational views of T-MAS and T-MAS-OH. (A)** Chemical structure of T-MAS. **(B)** Conformational view of T-MAS showing axial (C-29) and equatorial (C-28) C4-methyl groups. **(C)** Chemical structure of T-MAS-OH, the first enzymatic intermediate (C4α-alcohol) in the CYP123-catalysed three-step oxidation sequence. **(D)** Conformational view of T-MAS-OH.

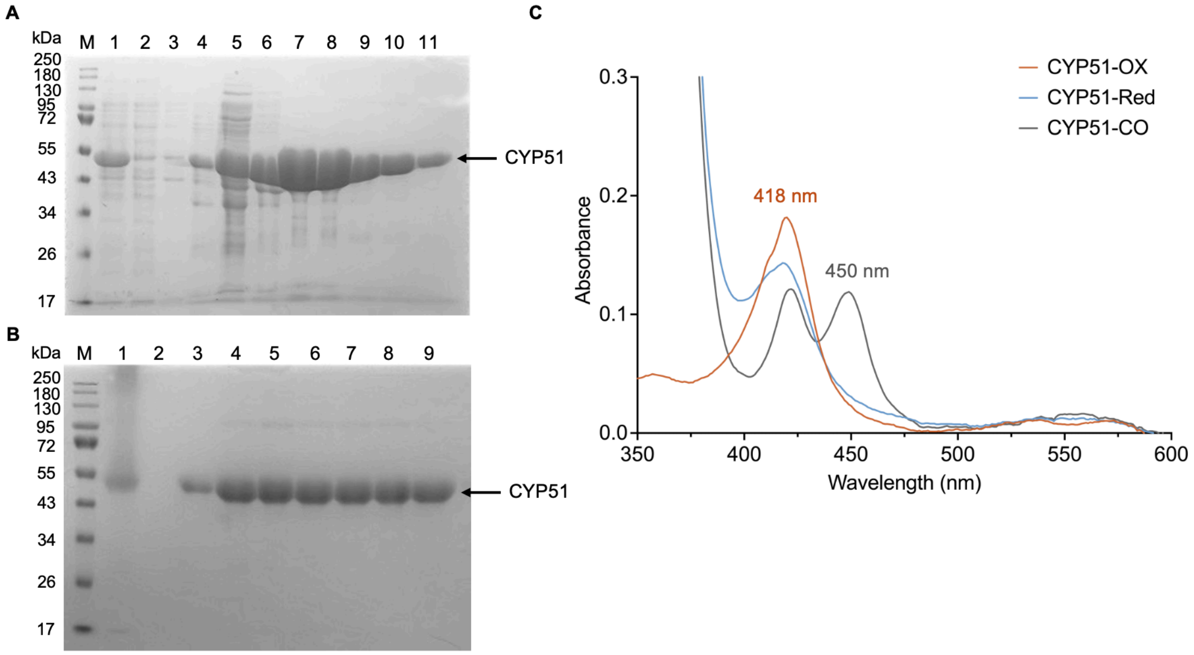


**Figure S33: Purification and spectroscopic characterisation of *M. smegmatis* CYP51. (A)** SDS-PAGE analysis of *M. smegmatis* CYP51 overexpression and Ni–NTA affinity purification. The gel lanes are: protein ladder (M), supernatant of protein-overexpressing cell pellets (lane 1), sample after washing the Ni–NTA affinity column (lane 2), fractions eluted from the Ni–NTA affinity column (lanes 3–11), with the target CYP51 protein exhibiting a size of 52.6 kDa. **(B)** SDS–PAGE analysis of samples collected during size-exclusion chromatography (SEC). The gel lanes are: protein ladder (M), concentrated Ni–NTA eluate (lane 1), fractions collected from the SEC peak (lanes 2–9). **(C)** UV-visible spectra for different states of *M. smegmatis* CYP51. The ferric resting state (orange), the reduced enzyme (blue) and the reduced CO-bound (grey) spectra are shown. The enzyme has a split A450/A418 peak in the CO-bound state.

**Table S5: X-ray data collection, processing and refinement statistics.**

| **PDB ID** | **9TJO**<br>**apo-CYP123** | **9TJP**<br>**PEG-CYP123** |
|---|---|---|
| **Data collection** | | |
| Space group | P 43 21 2 | P 43 21 2 |
| Cell dimensions *a, b, c* (Å) | 132.3, 132.3, 126.8 | 132.9, 132.9, 127.2 |
| Cell angles *α, β, γ* (°) | 90.0, 90.0, 90.0 | 90.0, 90.0, 90.0 |
| Resolution (Å) | 66.16-2.14<br>(2.20-2.14) | 66.45-2.19<br>(2.25-2.19) |
| Rmeas | 0.117 (3.161) | 0.173 (3.857) |
| Rsym or Rmerge | 0.115 (3.089) | 0.170 (3.785) |
| CC1/2 | 1.000 (0.512) | 0.999 (0.584) |
| $I / \sigma I$ | 17.5 (1.2) | 13.8 (1.1) |
| Completeness (%) | 100.0 (100.0) | 100.0 (100.0) |
| Redundancy | 27.2 (25.3) | 27.3 (27.0) |
| **Refinement** | | |
| Resolution (Å) | 66.25-2.14 | 66.54-2.19 |
| No. reflections | 62498 / 3239 | 59056 / 3072 |
| $R^{work}$ / $R^{free}$ | 0.220 / 0.262 | 0.210 / 0.272 |
| Ramachandran | | |
| Favoured (%) | 96.32% | 96.83% |
| Outlier (%) | 0.13% | 0.00% |
| No. atoms | | |
| Protein | 12297 | 12297 |
| Ligand/ion | 139 | 186 |
| Water | 189 | 167 |
| B-factors | | |
| Protein | 60.7 | 56.5 |
| Ligand/ion | 51.7 | 53.2 |
| Water | 49.9 | 46.4 |
| R.m.s. deviations | | |
| Bond lengths (Å) | 0.0044 | 0.0072 |
| Bond angles (°) | 1.251 | 1.671 |

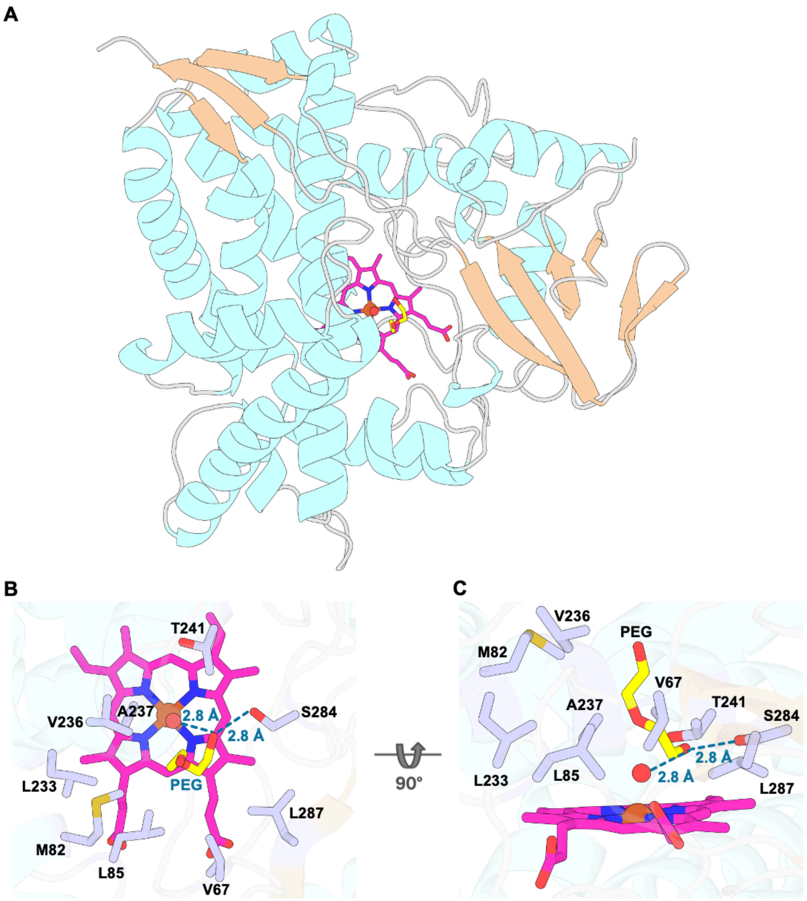


**Figure S34: Structural analysis of the PEG-bound CYP123 complex solved at 2.19 Å. (A)** Ribbon and surface diagram of CYP123 bound to a PEG ligand (in yellow sticks) introduced by the cryoprotectant. The protein is shown with α-helices in cyan, β-sheets in light orange, and loops in grey. The haem prosthetic group is shown in magenta sticks. The PEG ligand occupies a channel-like cavity leading into the active site. **(B)** Close-up view of the active site, highlighting key residues interacting with the PEG ligand. The ligand is primarily stabilised by a hydrophobic cluster (V67, M82, L85, L233, V236, A237, L287), with a potential hydrogen bond formed with S284 and the distal water molecule (red sphere), helping to orient PEG within the pocket. **(C)** Side view of the PEG-bound active site rotated 90° relative to panel B, showing the position of PEG within the binding pocket above the haem group, where it is stabilised by a water-mediated hydrogen-bonding network involving S284.

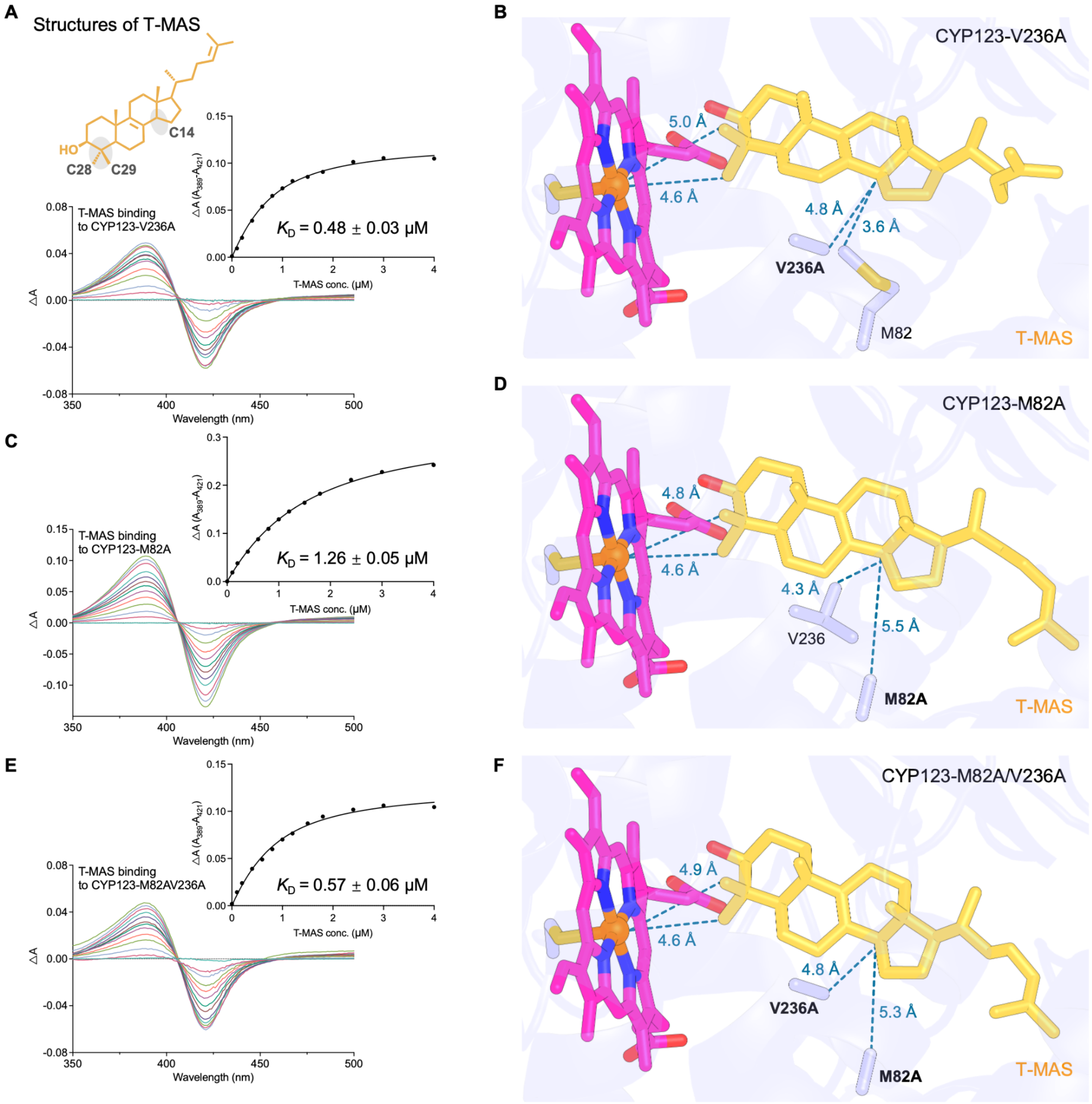


**Figure S35: Effects of CYP123 V236A and M82A variants on T-MAS binding. (A, C, E)** T-MAS binding to CYP123-V236A, CYP123-M82A and CYP123-M82A/V236A, respectively, with the corresponding $K_D$ values of 0.48 ± 0.03, 1.26 ± 0.05 and 0.57 ± 0.06 μM. Insets show the peak-to-trough absorbance differences plotted against T-MAS concentration. $K_D$ values are reported as best-fit ± SE. **(B, D, F)** Corresponding docking models showing the predicted binding poses of T-MAS relative to the haem, M82 and V236, or their alanine substitutions. Selected distances between T-MAS, the haem and active-site residues are indicated by dashed lines and reported in Å.

```
tr|A0A4S8NE54|A0A4S8NE54_9ACTN    ----------MVSTSSTTERPAAPLRFDPYDYDFHEDPYPTYQRLRDEDPVHHAPVDDIW 50
tr|J9SN57|J9SN57_9ACTN            MM----------TESTTVSGMEHELPFDPYAYGFHEDPYPTYHRLRTESPVYYHSGMDFW 50
tr|A0A1X1YJJ8|A0A1X1YJJ8_9MYCO    ----MASQAVNSETSKISE-TSDSPVLNPYDYDFHEDPFPYYRRLRDEAPVYHNEELNFW 55
tr|I7G922|I7G922_MYCS2            ----------------MSL-PTVEPVLDPYDYDFHEDPYPYYKRLRDEAPLYRNDERNFW 43
tr|A0A1E3RVP1|A0A1E3RVP1_9MYCO    -MKPRGGSLDEPGRTTVTV-PTSDLVLDPYDYDFHEDPYPYYQRLRDEAPLYRNDEIGFW 58
tr|A0A022LRJ6|A0A022LRJ6_9ACTN    ----------------MTTATPAPLVFSPYDYAVHEDPYPFYARLRAEAPVYHNPDLDFW 44
tr|A0A4R5BLK6|A0A4R5BLK6_9PSEU    --------------------MSSGVVFSPYDYAVHADPYPTYARLRDEAPVYHNAELDFW 40
tr|A0A7W7MZG0|A0A7W7MZG0_9ACTN    ------------------MAPAAQIFYSPFDYAVHEDPYPCYARLRDEAPLYRNDELGFW 42
tr|A0A0B8NAU3|A0A0B8NAU3_9NOCA    ----------------MTAASVEPLLFDPYDYRVHEDPYPVYQRLRAEAPVYHHPELDFW 44
tr|A0A562E8M0|A0A562E8M0_RHORH    MIQASPGSASHAGPASPDPPTLGPVIFNPYDYAFHEDPYPTYRRLREEDPAYHNPDLGFW 60
                                                             .*: * .* **:* * *** * * :     .:*

tr|A0A4S8NE54|A0A4S8NE54_9ACTN    VISRHEDVFDVLRRDDVFSNRMGVSLDATAWNPDAHKVMSFLGLDGAEQARVRKLVSAAF 110
tr|J9SN57|J9SN57_9ACTN            ALAGHADVRAAFRDTQRLSNSWGVSMDPSAYGPDAHKSMSFLAMDDPKHMRIRKLVSKGF 110
tr|A0A1X1YJJ8|A0A1X1YJJ8_9MYCO    AISRHQDVLHGFRNSDALSNRDGVSLDPVSRGPHAQLVMSFLAMDDPQHLRLRTLVSKGF 115
tr|I7G922|I7G922_MYCS2            ALSRHEDVLKGFRNSTALSNKHGVSLDPVSRNDEAHRVMSFLALDDPAHLRLRTLVSKGF 103
tr|A0A1E3RVP1|A0A1E3RVP1_9MYCO    ALSRHTDVLQGFRNSTTLSNQYGVSLDPASRGPHAAKTMSFLAMDDPAHLRLRTLVSKGF 118
tr|A0A022LRJ6|A0A022LRJ6_9ACTN    ALSRHADVREAFRAPDVFSNSMGVSLDPAAYGRHAYKTMSFLAMDDPRHARLRKLVSKGF 104
tr|A0A4R5BLK6|A0A4R5BLK6_9PSEU    ALSRYSDVAALFRDNTRLSNANGVTLDPAAWGPYAHKTMSFLALDDPRHNRLRSLVSRGF 100
tr|A0A7W7MZG0|A0A7W7MZG0_9ACTN    ALSRHADVSAAFRDHATFSNSHGVSLEPSAWGPQAHRTMSFLALDPPRHTRMRALVSKGF 102
tr|A0A0B8NAU3|A0A0B8NAU3_9NOCA    ALSRHVDVAAGFRDNVRLSSTNGVSLDPAAWGPHAWRTMSFLAMDDPRHLRLRKLVYQGF 104
tr|A0A562E8M0|A0A562E8M0_RHORH    ALSRHADVVAAFRDNTRLSSANGVSLDPSAYGPNAHKVMSFLAMDDPRHMRMRRLVSKGF 120
                                  .:: : **   :*    :*.  **:::  : .  *   ****.:*   : *:* **  .*

tr|A0A4S8NE54|A0A4S8NE54_9ACTN    TPRRVRELTPEVQQLTEKYLASTLAINA-EHGEADWIKDFAGKLPMDVISEMMGVPVSDR 169
tr|J9SN57|J9SN57_9ACTN            TPRRVNELTGRITALTHQHWSKCL-----DKGEFDFVHDFAALLPMDVVSELLGVPEADR 165
tr|A0A1X1YJJ8|A0A1X1YJJ8_9MYCO    TPRRIRELEPQVVRLARQHLDSAIQPDARGTCSFDFINDFAGKLPMDVISELMGVPEADR 175
tr|I7G922|I7G922_MYCS2            TPRRIRELEARVTEIAVQHLEIAL---Q--SDSFDFVDDFAGKLPMDVISELMGVPEEDR 158
tr|A0A1E3RVP1|A0A1E3RVP1_9MYCO    TPRRIRELEPRVTEIAVQHLDTMLEKAQ--SGTVDYVEEFAGKLPMDVISELMGVPEPDR 176
tr|A0A022LRJ6|A0A022LRJ6_9ACTN    TSRRVDALSEGIARISRRYWDECLATS-RDGDEFDFVQDYAGKLPMDVISDLMGVPESDR 163
tr|A0A4R5BLK6|A0A4R5BLK6_9PSEU    TPRRVRDLQPRILELARRHLHPAL-----ESGEFDFVADFAGKFPMDVISELIGVPEADR 155
tr|A0A7W7MZG0|A0A7W7MZG0_9ACTN    TPRRVKEMGDGIRALARRHLEPAL-----EKREFDFVSDFAGLLPMDVISEMMGVPEADR 157
tr|A0A0B8NAU3|A0A0B8NAU3_9NOCA    TPRRVTEMSARIREITLRYLEPAL-----AGGDFDWIDAVAGQVPMDVISELLGVPESDR 159
tr|A0A562E8M0|A0A562E8M0_RHORH    TPRRVADLEERILELTLRYLEPAV-----EAGEFDWISEFAGKLPMDVISELMGVPESDR 175
                                  * **:  :   :  :: ::    :          *::   *. .****:*:::***  **

tr|A0A4S8NE54|A0A4S8NE54_9ACTN    DEVRRLADLVVHREDGVRDVPVAGMEASMSLFGYYGDMVSQRRAQPTDDLTSALLAAEIE 229
tr|J9SN57|J9SN57_9ACTN            TRLRHQSDLLLHREDGDLDLPEAAVYAYIELHNYYSELIADRRKNPGEDLVSALIAAEID 225
tr|A0A1X1YJJ8|A0A1X1YJJ8_9MYCO    ARIRELADGVLHREDGVTDVPQSAIQASMDLMTYYNEMIADFKKNPADNLTSALLETEVD 235
tr|I7G922|I7G922_MYCS2            VRIRALADGVMHREDGVADVPASAIAASGELLVYYADMVKRRRRNVSDDLTSALVEAEID 218
tr|A0A1E3RVP1|A0A1E3RVP1_9MYCO    DQVRGWADDVMHRDDGVTDVPPAAVEASINLIVYYQGMVAERRKKPTDDLTTALLEAEID 236
tr|A0A022LRJ6|A0A022LRJ6_9ACTN    DELRRLADLVMHREEGGQDVPEDSMVAALDLITYYQGMVSERRRERTEDLTSALIDAEID 223
tr|A0A4R5BLK6|A0A4R5BLK6_9PSEU    DRLRELADTLVHREEGVFDVPQTAMEAALHLVAYYSDLLAERRRKRADDLTSALLDAEID 215
tr|A0A7W7MZG0|A0A7W7MZG0_9ACTN    VEVRRLADLVVHREEGLNDVPPAGMDAALTLVGYYQDMVAERRRRPADDLTSALLDAEID 217
tr|A0A0B8NAU3|A0A0B8NAU3_9NOCA    AELRRLADLLVHREEGVLDVPVAAMEAALQLAGYYNEVLAARKRQRTGDLTSALLDAEAD 219
tr|A0A562E8M0|A0A562E8M0_RHORH    SEIRRLADLVVHREEGVLDVPVAAMEASLHLVGYYADMLAERRKKETSDLTSALLEAEID 235
                                   .:*  :* ::**::*  *:*  .: *   *  **  ::   : .   :*.:**: :* :

tr|A0A4S8NE54|A0A4S8NE54_9ACTN    GE------KLRDGEVMAFLFLMVVAGNETTTKLLGNALFHLGAHPEQQAAVLGAPEDPDT 283
tr|J9SN57|J9SN57_9ACTN            DDETGEKTRLTEDEIVGFMVLMVVAGNETTTKLLANALYWGWRNPDELAKVFAD---PEV 282
tr|A0A1X1YJJ8|A0A1X1YJJ8_9MYCO    GD------RLGDQEILAFLFLMVIAGNETTTKLLGSSIYWGHRNRDQLTSVYAD---HSR 286
tr|I7G922|I7G922_MYCS2            GD------KLTDDEIMAFLFLMVVAGNETTTKLLANAAYWGFKNPDQLAPVFDD---HSQ 269
tr|A0A1E3RVP1|A0A1E3RVP1_9MYCO    GD------RLTDDEVLGFMFLMVIAGNETTTKLLANAAFWGHKNPDQLVPVYAD---LER 287
tr|A0A022LRJ6|A0A022LRJ6_9ACTN    GE------KLDEDEILSFLFLMVVAGNETTTKLLANAMFWGHRNPDQSQLVIED---PER 274
tr|A0A4R5BLK6|A0A4R5BLK6_9PSEU    GD------RLSDEEIIAFLVLMVVAGNETTTKLLANAAYWAHRNPEEQAKPLID---PGR 266
tr|A0A7W7MZG0|A0A7W7MZG0_9ACTN    GD------RLDDTEVIAFLFLMVVAGNETTTKLLANALYWASLNPEQAAKPFGD---PGR 268
tr|A0A0B8NAU3|A0A0B8NAU3_9NOCA    GD------RLTDEEILGFMFLMVVAGNETTTKLLGNALYWAGRNPGEYAKVAGQ---PEL 270
tr|A0A562E8M0|A0A562E8M0_RHORH    GD------RLTDDEIIGFMFLMVVAGNETTTKLLGNALYWGAHNKSEVAQVLHD---PAR 286
                                  .:      :* : *::.*:.***:**********..: :    :  :

tr|A0A4S8NE54|A0A4S8NE54_9ACTN    LVVPWIEETLRHDTSSQMLARYVTEDTEVAGITIPGGSKALVLLGSANRDERVFG-DPTV 342
tr|J9SN57|J9SN57_9ACTN            VVPDWTEETLRYDNSTQIVVRRVVEDAPYGDFVIPAGDRVLLLVGSANRDVEVFGADADR 342
tr|A0A1X1YJJ8|A0A1X1YJJ8_9MYCO    -IPLWVEETLRYDTPSQILARTAAQDYTLHGTTIPEGAVVLLLPGSGNRDERVFD-DPDA 344
tr|I7G922|I7G922_MYCS2            -IPLWVEETLRYDTSSQILARAVVEDLTFYGTTVPAGDVLVLLAGSANRDERAFD-DPDE 327
tr|A0A1E3RVP1|A0A1E3RVP1_9MYCO    -VPLWVEETLRYDTSSQILARTVVGDLTLYDTKIPDGDVVLLLPGSGHRDERVFD-EPDR 345
tr|A0A022LRJ6|A0A022LRJ6_9ACTN    -VSEWVEETLRYDTSSQLVLRTTMQDVTVRGVTIPADQKVLLLIGAANRDEEVFE-DPET 332
tr|A0A4R5BLK6|A0A4R5BLK6_9PSEU    -VTDWIEETLRYDTSTQVIARTATEEIRLHETTIPTGARVLLMPGSANRDPRAFD-DADR 324
tr|A0A7W7MZG0|A0A7W7MZG0_9ACTN    -VDDWVEETLRYDTSSQMLARLVTTDIELHGQRVPAGDRMLLLVGSANRDPRVFE-DADS 326
tr|A0A0B8NAU3|A0A0B8NAU3_9NOCA    -VEDWVEETLRYDASSQILARTAAEDLEYYGQTIPQGSKVLLLIGSANRDSAVFH-DADR 328
tr|A0A562E8M0|A0A562E8M0_RHORH    -APQWTEETLRYDTSSQIVARTAVADIELHDATIRSGDKVLLLIGSANRDSDVFD-AADD 344
                                      * *****:*  :*:: * .  :       :  .   ::: *:.:**  .*

tr|A0A4S8NE54|A0A4S8NE54_9ACTN    FNVHREKSELSQILSFGTGRHFCLGANLARLEAKVVLNELVRQVDGFQVHADRAVRVHST 402
tr|J9SN57|J9SN57_9ACTN            YEIGRDCG--QALMSFGLGAHFCLGAHLARLEANIGLTEVVKSIRAVDIDADKAVRVHSV 400
tr|A0A1X1YJJ8|A0A1X1YJJ8_9MYCO    FRIGRDIG--SKLLSFGSGAHFCLGAHLARMEARVALDELFNRISDYQVDEANAVRVHSS 402
tr|I7G922|I7G922_MYCS2            YRIGRDIG--SKLVSFGSGAHFCLGAHLARMEARVALAELFKRIRGYDVDESASVRVHSS 385
tr|A0A1E3RVP1|A0A1E3RVP1_9MYCO    FLIGRDIG--AKLLSFGSGAHFCLGAHLARMEARVALTELLKRIRRYEVDEAGAVRVHSS 403
tr|A0A022LRJ6|A0A022LRJ6_9ACTN    FRLGRDTS---ASLSFGQGPHFCLGAHLARLEGRIALDQISRSITGYEIL-PGAERVHSS 388
tr|A0A4R5BLK6|A0A4R5BLK6_9PSEU    YDLDRSAD--TKIVSFGGGTHFCLGAHLARLEARIALTEFVELVKSYDVDTDRAQRVHSS 382
tr|A0A7W7MZG0|A0A7W7MZG0_9ACTN    YDLDRDT---SQLVSFGGGRHFCLGANLARLEARIALGELVRRVSSFEVDHAGAERVHSV 383
tr|A0A0B8NAU3|A0A0B8NAU3_9NOCA    YDIERPDK--GGLISFGRGVHFCLGAHLARLEASIVLREFVARVRGYGIGESGIQRVHSV 386
tr|A0A562E8M0|A0A562E8M0_RHORH    FRIGRDSS--QKIASFGGGVHFCLGAHLARLEANIALEQFARRVADYEIDEDRCERVHST 402
                                  : : *         *** * ******:***:*. : * :.   :    :      ****

tr|A0A4S8NE54|A0A4S8NE54_9ACTN    SVRGFASLPVTFRPRSA--- 419
tr|J9SN57|J9SN57_9ACTN            NVRGFAELPVKVTVR----- 415
tr|A0A1X1YJJ8|A0A1X1YJJ8_9MYCO    NVRGFADLPITVKEA----- 417
tr|I7G922|I7G922_MYCS2            SVRGFAHLPITVEKR----- 400
tr|A0A1E3RVP1|A0A1E3RVP1_9MYCO    NVRGFAHLPITLEIN----- 418
tr|A0A022LRJ6|A0A022LRJ6_9ACTN    NVRGFAHLPMRVEVRRS--- 405
tr|A0A4R5BLK6|A0A4R5BLK6_9PSEU    NVRGFATLPVRVESR----- 397
tr|A0A7W7MZG0|A0A7W7MZG0_9ACTN    NVRGFAALPVRVVGRGRKES 403
tr|A0A0B8NAU3|A0A0B8NAU3_9NOCA    NVRGFAHLPVSVEVR----- 401
tr|A0A562E8M0|A0A562E8M0_RHORH    NVRGFAALPVEVTERHA--- 419
                                  .***** **: .
```

**Figure S36: Comparative sequence alignment of CYP123 enzymes across diverse bacterial species.** Multiple sequence alignment of CYP123 enzymes from phylogenetically diverse actinobacteria: *Nocardioides caeni* (A0A4S8NE54), *Gordonia* sp. KTR9 (J9SN57), *Mycolicibacter longobardus* (A0A1X1YJJ8), *M. smegmatis* mc$^2$155 (I7G922), *Mycolicibacterium holsaticum* (A0A1E3RVP1), *Dietzia* sp. UCD-THP (A0A022LRJ6), *Saccharopolyspora karakumensis* (A0A4R5BLK6), *Actinomadura livida* (A0A7W7MZG0), *Nocardia seriolae* (A0A0B8NAU3), and *Rhodococcus rhodochrous* J45 (A0A562E8M0). The *M. smegmatis* CYP123 sequence serves as the reference; other sequences were selected from the 472 members of SSN Cluster 24 (Fig. S4) whose genome neighbourhoods carry the conserved gene set of Table S6, using CD-HIT v4.8.1 as representatives of groups sharing ≥70% identity over ≥80% of their length (cd-hit -c 0.7 -n 5 -aS 0.8); 472 sequences were clustered into 9 groups. Highlighted in yellow: the haem-binding motif FGSGAHFCLG (residues 341–350 in *M. smegmatis*), matching the canonical cytochrome P450 PROSITE signature FGxGxHxCxG and containing the axial cysteine ligand (Cys348). Arg96 and Arg289, which may also participate in haem coordination, are highlighted in yellow. Highlighted in blue: the I-helix motif LMVVAGNETTTKLL (residues 233–246), located deep within the binding pocket adjacent to the haem, potentially involved in substrate coordination and positioning for successive oxidations. Also notable are well-conserved binding-pocket residues Val67 and Leu85, positioned on opposite sides of the active-site cavity. The alignment was generated and visualised using EMBL-EBI Clustal Omega (v1.2.4).

**Table S6: Conserved genome neighbourhood of bacterial CYP123.** Frequency of neighbouring Pfam families within ten open reading frames of each query, determined with the EFI Genome Neighbourhood Tool (neighbourhood size 10, co-occurrence threshold 20%), for the 472 Cluster 24 sequences from which the Figure S36 representatives were selected and for the remaining 427 members of Cluster 24. PF00106/PF13561, PF03446/PF14833 and PF00107/PF08240 are domain pairs of single proteins.

| Pfam | Family | n = 472 | % | n = 427 | % |
|---|---|---|---|---|---|
| PF00106 | Short-chain dehydrogenase/reductase | 472 | 100.0 | 290 | 67.9 |
| PF13561 | Short-chain dehydrogenase/reductase, C-terminal | 472 | 100.0 | 222 | 52.0 |
| PF00067 | Cytochrome P450 | 471 | 99.8 | 239 | 56.0 |
| PF00440 | TetR-family regulator, HTH domain | 471 | 99.8 | 306 | 71.7 |
| PF02627 | Carboxymuconolactone decarboxylase-like | 468 | 99.2 | 185 | 43.3 |
| PF03446 | NAD(P)-binding oxidoreductase, N-terminal | 468 | 99.2 | 198 | 46.4 |
| PF14833 | NAD(P)-binding oxidoreductase, C-terminal | 468 | 99.2 | 179 | 41.9 |
| PF00171 | Aldehyde dehydrogenase | 466 | 98.7 | 230 | 53.9 |
| PF13459 | Ferredoxin | 466 | 98.7 | 213 | 49.9 |
| PF00107 | Zn-binding alcohol dehydrogenase, C-terminal | 327 | 69.3 | 49 | 11.5 |
| PF08240 | Zn-binding alcohol dehydrogenase, GroES-like | 327 | 69.3 | 53 | 12.4 |

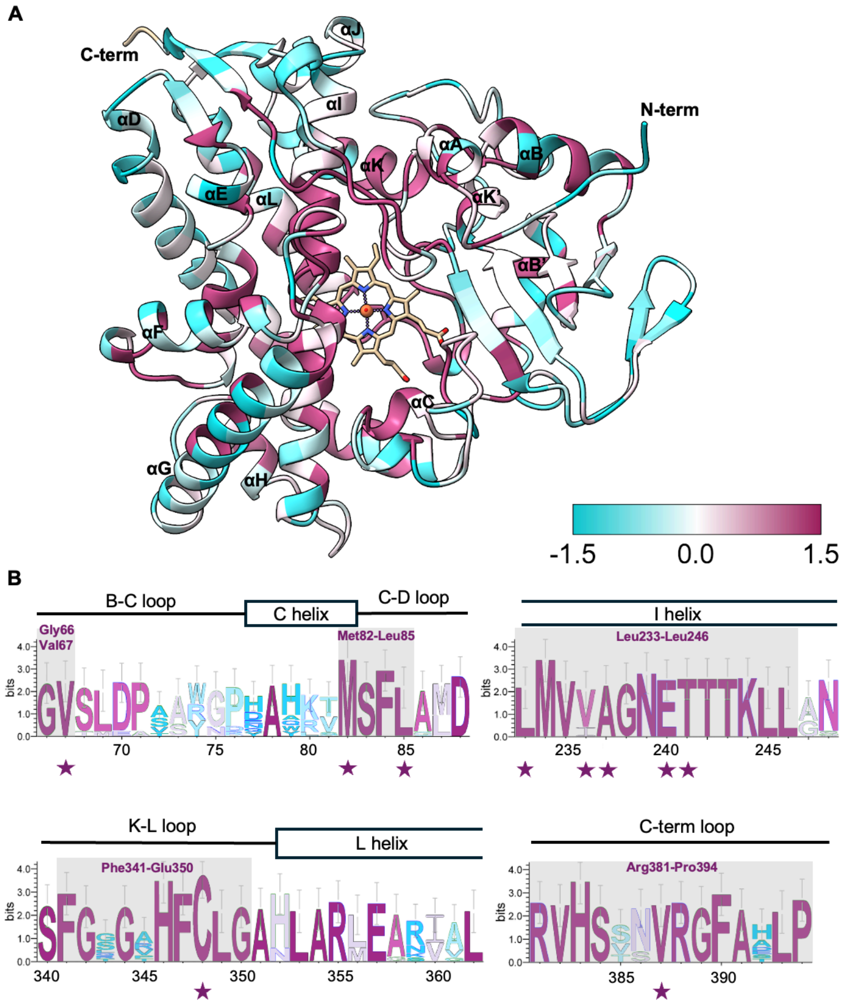


**Figure S37: Sequence conservation mapped onto the CYP123 structure. (A)** Per-residue conservation derived from the multiple sequence alignment of Fig. S36, mapped onto the *M. smegmatis* CYP123 crystal structure (PDB: 9TJO) in UCSF ChimeraX using the Coloring by Sequence Conservation function and shown as a cyan-to-maroon gradient (range −1.5 to +1.5; maroon = higher conservation). Highly conserved regions highlighted in grey include: the I-helix core (Leu233–Leu246), the haem-binding motif (Phe341–Gly350, including the axial ligand Cys348), and a C-terminal loop (Arg381–Pro394) near the substrate channel. Key residues potentially involved in substrate binding and catalysis (Val67, Met82–Leu85, Leu233–Ala237, Glu240–Thr241, and Val387) are also highlighted. **(B)** Sequence logo of the same alignment, generated with WebLogo and coloured to match the gradient in (A).

**Table S7: Bacterial strains and plasmids used in this study.**

| Strains/Plasmids | Description | Source |
|---|---|---|
| **Strains** | | |
| *E. coli* DH5α | supE44 ΔlacU169 (ϕ80 lacZΔM15) hsdR17 *recA*1 endA1 gyrA96 thi-1 relA1 | New England Biolabs |
| *E. coli* BL21 (DE3) | F- ompT hsdSB (rB- mB-) gal dcm (DE3) | New England Biolabs |
| *E. coli* Rosetta2 (DE3) | F- ompT hsdSB (rB- mB-) gal dcm (DE3) pRARE2 (CamR) | Sigma-Aldrich |
| *Mycobacterium smegmatis* mc$^2$155 (ATCC 700084) | Type strain | ATCC |
| *Mycobacterium abscessus* (ATCC 19977) | Type strain | UKHSA |
| *Mycobacterium phlei* (DSM 43239) | Type strain | DSMZ |
| *Mycobacterium bovis* var. BCG (NCTC 14044) | Reference strain | UKHSA |
| *Gordonia bronchialis* (NCTC 10667) | | UKHSA |
| *Rhodococcus coprophilus* (NCTC 10994) | Type strain | UKHSA |
| *Mycobacterium marinum* (NCTC 2275) | Type strain | UKHSA |
| *Nocardia africana* (NCTC 13184) | Type strain | UKHSA |
| *M. smegmatis cyp123*Δ17 | CRISPR edited *cyp123*, 17 bp deletion at 742–758 | This study |
| *M. smegmatis cyp123*Δ136ins228 | CRISPR edited *cyp123*, 228 bp insertion at 758, 136 bp deletion at 759–894 | This study |
| *M. smegmatis Δcyp123::hyg*-1 (Δ*cyp123*-1) | ORBIT edited *cyp123*, 1142 bp replaced by 3143 bp cassette | This study |
| *M. smegmatis Δcyp123::hyg*-2 (Δ*cyp123*-2) | ORBIT edited *cyp123*, *cyp123* replaced by 3142 bp cassette | This study |
| *M. smegmatis Δcyp51::hyg*-1 (Δ*cyp51*-1) | ORBIT edited *cyp51*, 1294 bp replaced by 3127 bp cassette | This study |
| *M. smegmatis cyp123*Δ17 + *cyp123* | CRISPR edited *cyp123*, 17 bp deletion at 742–758, complemented with *cyp123* | This study |
| *M. smegmatis cyp123*Δ136ins228 + *cyp123* | CRISPR edited *cyp123*, 228 bp insertion at 758, 136 bp deletion at 759–894, complemented with *cyp123* | This study |
| **Plasmids** | | |
| pET28a (+) | N-terminal 6xHis tag, T7 promoter, KanR, MCS, lacI repressor | EMD Biosciences |
| pET28-CYP123 | pET28a vector containing *cyp123* (MSMEG_5861) cloned into NcoI-XhoI, leading to a C-terminal $His_6$ tag | This study |
| pET28-CYP51 | pET28a vector containing *cyp51* (MSMEG_5863) cloned at the NcoI site by Gibson assembly, leading to a C-terminal $His_6$ tag | This study |
| pET28-CYP123-M82A | pET28-CYP123 carrying the M82A mutation in *cyp123*; constructed by site-directed mutagenesis | This study |
| pET28-CYP123-V236A | pET28-CYP123 carrying the V236A mutation in *cyp123*; constructed by site-directed mutagenesis | This study |
| pET28-CYP123-M82A/V236A | pET28-CYP123 carrying the M82A and V236A mutation in *cyp123*; constructed by site-directed mutagenesis | This study |
| pYC2085 | CRISPR, helper plasmid expresses Sth1 Cas9 and the cognate sgRNA | Addgene #158721 (10.1128/mBio.02364-19) |
| pNHEJ-recAmu-sacB | CRISPR, helper plasmid expresses NHEJ machinery of *M. marinum* and recAR61C allele | Addgene #158717 (10.1128/mBio.02364-19) |
| pNHEJ-recX-sacB | CRISPR, helper plasmid expresses NHEJ machinery of *M. marinum* and recX | Addgene #158718 (10.1128/mBio.02364-19) |
| pNHEJ-recX-sacB-d1311 | pNHEJ-recX-sacB derivative recovered from *cyp123*Δ136ins228, carrying a 1,311 bp deletion that removes *sacB* | This study, GenBank QB217605 |
| pKM461 | ORBIT helper plasmid; expresses Che9c RecT annealase and Bxb1 integrase under tet control, with *sacB* for counter-selection | Addgene #108320 (10.1128/mBio.01467-18) |
| pKM464 | ORBIT payload plasmid for gene deletion; carries a 38 bp Bxb1 *attB* site, *hyg* for selection in mycobacteria and *cat* for selection in *E. coli* | Addgene #108322 (10.1128/mBio.01467-18) |
| pML1335 | L5 integration vector for mycobacteria; carries *int* L5, the *gfpm2*+ and *xylEm* reporters and a hygromycin resistance marker | Addgene #32377 (10.1016/j.gene.2010.07.012) |
| pCYP123-comp | pML1335 derivative in which the *gfpm2*+ cassette is replaced by *cyp123*, integrating at the L5 *attB* site under the Psmyc promoter | This study, GenBank QB217606 |
| pBS-Int | Integration plasmid | Addgene #50000 (10.1016/S1438-4221(01)80004-7) |

**Table S8: Primers used throughout the study.**

| Function | Primer | Sequence (5'-3') | Notes |
|---|---|---|---|
| Cloning | pET28 Mtb *cyp123* Forward | CATGTCATGACCGTCCGCGTCGG | Restriction site cloning |
| | pET28 Mtb *cyp123* Reverse | ATAAGAATGCGGCCGCCCTGGCCTGCACGCT | |
| | pET28 Na *cyp123* Forward | CATGTCATGACTGTGGCATCTGCG | |
| | pET28 Na *cyp123* Reverse | AAGGAAAAAAGCGGCCGCCGAAACCTCCACGGTGAC | |
| | pET28 Rc *cyp123* Forward | CATGTCATGATTCAGGCCTCCCG | |
| | pET28 Rc *cyp123* Reverse | AAGGAAAAAAGCGGCCGCGGCATGTCGATCGGTGAC | |
| | pET28 Mma *cyp123* Forward | CATGTCATGAGCGTGCACGTGG | |
| | pET28 Mma *cyp123* Reverse | AAGGAAAAAAGCGGCCGCGGCGACCTCCACGGT | |
| | pET28 Gb *cyp123* Forward | CATGTCATGACCGAGATCGCCGT | |
| | pET28 Gb *cyp123* Reverse | AAGGAAAAAAGCGGCCGCGCGCACCTGCACCTTC | |
| | pET28 Ms *cyp123* Forward | CATGCCATGGGTTTGCCGACCGTCGA | |
| | pET28 Ms *cyp123* Reverse | CCGCTCGAGGCGTTTCTCCACTGTGATCG | |
| | pET28 MbB *cyp123* Forward | CATGTCATGACCGTCCGCGTCGGT | $T_a$ 61 °C, Expected size 1230 bp |
| | pET28 MbB *cyp123* Reverse | AAGGAAAAAAGCGGCCGCCCTGGCCTGCACGC | |
| | pET28 Forward | CTCGAGCACCACCACCACCACCACTG | Gibson Assembly cloning |
| | pET28 Reverse | CCATGGTATATCTCCTTCTTAAAGTTAAAC | |
| | pET28 Ms CYP51 Forward | CTTTAAGAAGGAGATATACCATGACCACGCTCCACGACGTACCGC | |
| | pET28 Ms CYP51 Reverse | GTGGTGGTGGTGGTGCTCGAGCCTGACCCGCCTGCGGTAGCGCAC | |
| | pET28 MsCYP123-M82A Forward | GTTTCTCGCACTCGACGATCCGGCACACCTGCGGTTGC | Site-directed mutagenesis |
| | pET28 MsCYP123-M82A Reverse | GATCGTCGAGTGCGAGAAACGACGCCACGCGATGCGCCTCGTCGTTG | |
| | pET28 MsCYP123-V236A Forward | CTCATGGTCGCGGCCGGTAACGAGACCACCACCAAACT | |
| | pET28 MsCYP123-V236A Reverse | CGTTACCGGCCGCGACCATGAGGAACAGGAACGCCATG | |
| | T7 Forward | TAATACGACTCACTATAGGG | Sequence verification |
| | T7 Reverse | GCTAGTTATTGCTCAGCGG | |
| CRISPR Cas | Ms *cyp123* pos762 Forward | GGGAGAATGCCGCCTATTGGGGTTT | |
| | Ms *cyp123* pos762 Reverse | AAACAAACCCCAATAGGCGGCATTC | |
| | *cyp123* Ontarget Guide RNA 762 Left | CGGTGAACTGCTCGTGTACT | Designed using CRISPOR, Ta 60 °C, Expected size 343 bp |
| | *cyp123* Ontarget Guide RNA 762 Right | TCGTGCCGTAGAACGTCAAA | |
| | pML1335 Forward | ATTTAAATAAGCTTTAGCTAATTAATTG | Gibson Assembly |
| | pML1335 Reverse | TATTAACCTCCTTTCTGTTAATTAAG | |
| | pML1335 Ms *cyp123* Forward | AACAGAAAGGAGGTTAATAATGAGTTTGCCGACCGTC | |
| | pML1335 Ms *cyp123* Reverse | ATTAGCTAAAGCTTATTTAAATTTAGCGTTTCTCCACTGTG | |
| ORBIT | Ms *cyp123* ORBIT oligo 1 | CGTGGTCGGGCTCATCCTGGGAGAGGACCGATGAGTTTGCCGACCGTCGAACCCGTACTCGG TTTGTCTGGTCAACCACCGCGGTCTCAGTGGTGTACGGTACAAACCAGAAAGGAGGAAGGACA CCTTCCGATCACAGTGGAGAAACGCTAAATGGCCCGTTTTGAACCCCACCCTGCCCGC | Used for *M. smegmatis* Δ*cyp123::hyg*-1 |
| | Ms *cyp123* ORBIT oligo 2 | CACCAGATCGCCGACCGCCTCACCTACGTGGTCGGGCTCATCCTGGGAGAGGACCGATGAGG TTTGTCTGGTCAACCACCGCGGTCTCAGTGGTGTACGGTACAAACCAGAAAGGAGGAAGGAAT GGCCCGTTTTGAACCCCACCCTGCCCGCAGGCCAGCGATCGTGGCCGGCGCGTCCTCC | Used for Δ*cyp123::hyg*-2 |
| | Ms *cyp123* ORBIT Detection Forward | GCCCTGGAGATGACGTTCTT | Amplification from either side of *cyp123*, |

| | | | |
|---|---|---|---|
| | Ms *cyp123* ORBIT Detection Reverse | AGTTCCTGGCACTTCTCGAC | $T_a$ 56 °C, Expected size for WT 1468 bp, for sequencing |
| | Ms *cyp51* ORBIT oligo | TGCCCTCGGTGAAGAAGGGATGCCGAACCAGTGACCACGCTCCACGACGTACCGCGGGTTG GTTTGTCTGGTCAACCACCGCGGTCTCAGTGGTGTACGGTACAAACCAAGGTGCGCTACCGCA GGCGGGTCAGGTAGTCCCATGGGTTGCTACCGCATCGAGCTCGA | Used for Δ*cyp51::hyg*-1 |
| | Ms *cyp51* ORBIT Detection Forward | TTGAGGATTGGGCCAAGTGG | Amplification from either side of *cyp51*, Expected size for WT 1750 bp, for sequencing |
| | Ms *cyp51* ORBIT Detection Reverse | ATTGACAGTGCTCGGGTGG | |
| qPCR | Ma 0101 Forward | TCTCATTCAGCACCGCATCC | Ta 59 °C, Expected size 77 bp |
| | Ma 0101 Reverse | GTTGCGCATTCCACCAGATG | |
| | Ma 0611 Forward | GGCAGAAGTTGTCCGACGAT | Ta 59 °C, Expected size 77 bp |
| | Ma 0611 Reverse | GTTGCGGGTTGTCTCATTGC | |
| | Ma 0613 Forward | GAGCAGGTGTCGTGTGAGTT | Ta 59 °C, Expected size 89 bp |
| | Ma 0613 Reverse | GACCAGTTAAAGAGCTTGCCG | |
| | Ma 1211c Forward | CGTCGTCATGCTCTATCCGT | Ta 59 °C, Expected size 71 bp |
| | Ma 1211c Reverse | TGATGTCGAAGGTATGCGGG | |
| | Ma *recA* Forward | CGTAAGTCCGGTTCCTGGTT | Ta 59 °C, Expected size 70 bp |
| | Ma *recA* Reverse | AGTTACGGGCGTTCTCCTTG | |
| | Ma 1214c Forward | CCGAGCTGACCATCTACACC | Ta 59 °C, Expected size 142 bp |
| | Ma 1214c Reverse | ATCTCCAGGTAGGGGTCCAC | |
| | Ma 1216c Forward | TAGCCGAAACGTCAGCAGTA | Ta 59 °C, Expected size 67 bp |
| | Ma 1216c Reverse | TACGGGTCTTCGTGGAAGTC | |

Ms- *Mycobacterium smegmatis*, Mtb - *Mycobacterium tuberculosis*, Na - *Nocardia africana*, Rc - *Rhodococcus coprophilus*, Mma - *Mycobacterium marinum*, Gb - *Gordonia bronchialis*, MbB - *Mycobacterium bovis* var. BCG, Ma - *Mycobacterium abscessus*, $T_a$ - annealing temperature.
